# Ganglioglioma deep transcriptomics reveals primitive neuroectoderm neural precursor-like population

**DOI:** 10.1101/2022.12.17.520880

**Authors:** Joshua A Regal, María E Guerra García, Vaibhav Jain, Vidyalakshmi Chandramohan, David M Ashley, Simon G Gregory, Eric M Thompson, Giselle Y López, Zachary J Reitman

**Affiliations:** Department of Radiation Oncology, Duke University, Durham, NC 27710, USA; Duke Molecular Physiology Institute, Duke University, Durham, NC 27710, USA; Department of Neurosurgery, Duke University, Durham, NC 27710, USA; Department of Pathology, Duke University, Durham, NC 27710, USA

**Keywords:** Single cell RNA-sequencing, CITE-seq, spatial transcriptomics, ganglioglioma, glioneuronal tumors, brain tumors, genetic signatures

## Abstract

Gangliogliomas are brain tumors composed of neuron-like and macroglia-like components that occur in children and young adults. Gangliogliomas are often characterized by a rare population of immature astrocyte-appearing cells expressing CD34, a marker expressed in the neuroectoderm (neural precursor cells) during embryogenesis. New insights are needed to refine tumor classification and to identify therapeutic approaches. We evaluated five gangliogliomas with single nucleus RNA-seq, cellular indexing of transcriptomes and epitopes by sequencing, and/or spatially-resolved RNA-seq. We uncovered a population of CD34+ neoplastic cells with mixed neuroectodermal, immature astrocyte, and neuronal markers. Gene regulatory network interrogation in these neuroectoderm-like cells revealed control of transcriptional programming by TCF7L2/MEIS1-PAX6 and SOX2, similar to that found during neuroectodermal/neural development. Developmental trajectory analyses place neuroectoderm-like tumor cells as precursor cells that give rise to neuron-like and macroglia-like neoplastic cells. Spatially-resolved transcriptomics revealed a neuroectoderm-like tumor cell niche with relative lack of vascular and immune cells. We used these high resolution results to deconvolute clinically-annotated transcriptomic data, confirming that CD34+ cell-associated gene programs associate with gangliogliomas compared to other glial brain tumors. Together, these deep transcriptomic approaches characterized a ganglioglioma cellular hierarchy - confirming CD34+ neuroectoderm-like tumor precursor cells, controlling transcription programs, cell signaling, and associated immune cell states. These findings may guide tumor classification, diagnosis, prognostication, and therapeutic investigations.

## Introduction

Glioneuronal tumors (GNTs) are brain tumors that are composed of both neoplastic neuron-like and macroglial-like components[40]. GNTs account for approximately 2% of all primary brain tumors and most often occur in children and young adults[40]. GNTs include ganglioglioma (GG)[40]. The majority of GNTs are slow-growing tumors that do not result in death of the patients[40]. Nevertheless, GNTs can be associated with significant morbidity due to associated seizure disorders, risks of surgical resection, and tumor recurrence[40]. The standard of care therapy is surgical resection[40]. However, new treatment strategies are needed for patients with anaplastic, recurrent, or progressive GNTs[40]. Genomic alterations in ganglioglioma are minimal[23, 58, 68]. Nonetheless, at least 90% of GGs have driver alterations in the BRAF/MAPK pathway, with about half of GGs carrying BRAF V600E[23, 58, 68]. Additional recurrent alterations include chromosome 7 gains with *BRAF*, *KIAA1549*, and *EGFR* copy number gains as well as homozygous *CDKN2A/B* loss[23, 58, 68]. BRAF/MEK targeted therapies are undergoing clinical testing for BRAF-mutant glioneuronal and other brain tumors, with early evidence suggesting clinically-significant anti-tumor activity[36, 47]. However, further understanding of how to molecularly target GNTs is needed. Additionally, immunotherapeutic advances have resulted in significant breakthroughs for many malignancies, but immunotherapies have not yet improved outcomes for GNTs[4, 29, 40, 84]. Better understanding of tumor microenvironment may help to identify opportunities for effective immunotherapy.

Opportunities exist to further understand GNT development and maintenance and thereby improve upon GNT treatment. There is now decades of work supporting the existence of cancer stem cells for myriad cancer types, and putative primitive progenitor cell types have been identified in other low grade brain tumors, such as pilocytic astrocytoma[8, 67]. Understanding cellular hierarchy is important for understanding tumorigenesis, tumor maintenance, and tumor treatment, but little is understood about the potential for or nature of GNT stem cells. ScRNA-seq approaches have revealed that many subtypes of brain tumors contain tumor cells that transcriptionally resemble normal brain progenitor cell subtypes[11, 20]. However, this approach has not yet been used to investigate GNTs. Many pediatric low grade gliomas such as gangliogliomas contain tumor cells that express CD34, (an endothelial and hematopoietic stem cell marker but also) a transient marker of neuroectodermal neural precursor cells during neural development[12, 21, 26]. The precise nature of these cells is not well understood. However, because of their primitive neuroectodermal neural precursor cell marker expression and histologic appearance, it is tempting to hypothesize that GNT CD34+ cells are neoplastic stem or precursor cells that transcriptionally resemble normal neuroectodermal neural precursor cells.

In addition to the therapeutic challenges, classification of GNTs is a major diagnostic challenge with resultant impacts on understanding of prognosis and medical decision making[40]. Brain tumor classification has evolved enormously over the last several years with the gradual incorporation of tumor genetic features leading to significant improvements in tumor classification and medical decision making[7, 44]. Tumor heterogeneity and associated under-sampling are notorious issues with brain tumors, resulting in significant rates of under-grading and under-treatment but can sometimes be compensated for to a large extent by accounting for genetic factors[89]. Many types of pediatric brain tumors are now being understood as separate entities, with implications for prognosis and medical decision making[7, 44].

Single nucleus (or cell) RNA-sequencing (snRNA-seq), cellular indexing of transcriptomes and epitopes by sequencing (CITE-seq), and spatial transcriptomics (stRNA-seq) have been carried out on a number of brain tumor subtypes to provide insights to guide the development of new therapeutic strategies and new classification schemes for the tumors[11, 20, 48, 50, 62, 66, 67, 78]. However, these high-resolution transcriptomic techniques have not yet been published for GNTs. As outlined above, a deep understanding of GNT cellular composition is needed to generate hypotheses for therapeutic interventions that may benefit select patients with GNT and to provide insights that may refine the GNT pathologic classification. We hypothesize that the GNT CD34+ cells transcriptionally resemble embryonic neuroectodermal (neural precursor) cells that may serve as progenitors for the other neuron-like and macroglia-like tumor cells. We additionally hypothesize that a detailed definition of GNT immune cell composition and activation status will further the understanding of the utility of immunotherapeutic targeting for GNT. Consequently, we endeavored to characterize a major GNT subtype (ganglioglioma) using snRNA-seq, CITE-seq, and stRNA-seq.

## Results

### Neoplastic neuron-like, macroglia-like, and *CD34*+ neuroectoderm-like components in GNTs

To elucidate normal and neoplastic cell states in GNTs, single nucleus RNA-seq was carried out on 4 GG tumors (**sTable 1**). All tumors had a BRAF V600E driver. To our knowledge, no clinical testing revealed additional protooncogene or tumor suppressor mutation or copy number variation. SnRNA-seq yielded high-quality RNA-seq profiles from 34,907 nuclei. Nuclei were subjected to non-linear dimensional reduction and clustered based on k-nearest neighbor search[15, 32, 70, 74] yielding 30 clusters (**Figure 1A**). Nuclei largely segregated by tumor-of-origin except for select normal-appearing clusters (**sFigure 1A-G**). Cell types were identified using k-nearest neighbors clustering, hypothesis-driven markers, analysis of top differentially expressed genes, inference of chromosomal copy number variations, and Seurat-based label transfer.

**Figure 1:**
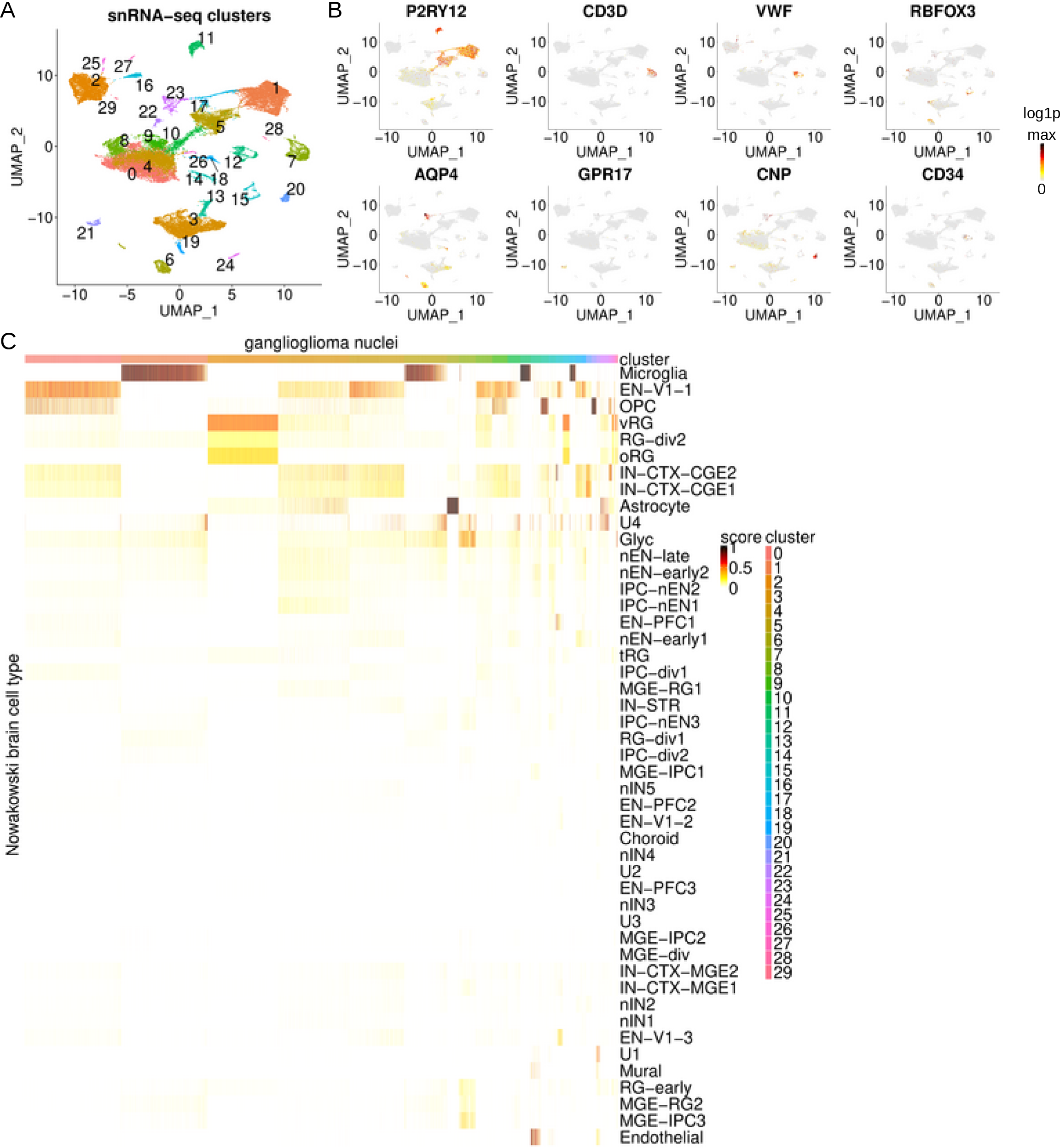
Ganglioglioma single nucleus RNA-seq. (A) UMAP plot of SCTransformed data from 34,907 nuclei, colored by Seurat cluster. (B) Feature plots of log1p[RNA] of select cell typing markers, *P2RY12* (microglia), *CD3D* (T lymphocyte), *VWF* (endothelial), *RBFOX3* (pan-neuron), *AQP4* (astrocyte), *GPR17* (oligodendrocyte precursor cell, OPC), *CNP* (oligodendrocyte), and *CD34* (endothelial and stem cell). Colored from 0 (grey), to maximum log1p for the transcript plotted (black), via yellow, then orange, then red. (C) Heatmap of Seurat label transfer prediction scores (from 0 to 1) for ganglioglioma nuclei queried against developing brain (Nowakowski *et al*.) reference atlas with query nuclei as columns and reference annotations as rows. Note cluster colors in legend should be the same as cluster colors throughout. EN-V1-1=early born deep layer/subplate excitatory neuron V1; OPC=oligodendrocyte precursor cell; vRG=ventricular radial glia; RG-div2=dividing radial glia (S-phase); oRG=outer radial glia; IN-CTX-CGE1 or 2=CGE/LGE-derived inhibitory neurons; U1, U2, U3, and U4= unknown 1, 2, 3, and 4; glyc=glycolysis; nEN-late=newborn excitatory neuron - late born; nEN-early1 or 2=newborn excitatory neuron - early born; IPC-nEN1, 2, or 3=intermediate progenitor cells excitatory neuron-like; EN-PFC1=early born deep layer/subplate excitatory neuron prefrontal cortex; tRG=truncated radial glia; IPC-div1=dividing intermediate progenitor cells radial glia-like; MGE-RG1=medial ganglionic eminence radial glia 1; IN-STR=striatal neurons; RG-div1=dividing radial glia (G2/M-phase); IPC-div2=intermediate progenitor cells radial glia-like; MGE-IPC1, 2, or 3=medial ganglionic eminence progenitors; nIN1, 2, 3, 4, or 5=medial ganglionic eminence newborn neurons; EN-PFC2 or 3=early and late born excitatory neuron prefrontal cortex; EN-V1-2=early and late born excitatory neuron V1; MGE-div=dividing medial ganglionic eminence progenitors; IN-CTX-MGE2=medial ganglionic eminence-derived Ctx inhibitory neuron, cortical plate-enriched; IN-CTX-MGE1=medial ganglionic eminence-derived Ctx inhibitory neuron, germinal zone enriched; EN-V1-3=excitatory neuron V1 - late born; RG-early=early radial glia; MGE-RG2=medial ganglionic eminence radial glia 2.

Normal brain, immune cell, and vascular clusters expressed classic cell type markers (**Figure 1B and sFigure 1H**). Normal clusters included macroglia comprising astrocytes (*GFAP*-, *AQP4*-, and *ALDH1L1*-expressing, cluster 23), oligodendrocytes (*MBP*-, *MOG*-, *CNP*-, and *MOBP*-expressing, cluster 20), and oligodendrocyte precursor cells (OPCs, *GPR17*-, *CSPG4*-, *OLIG1*-, and *OLIG2*-expressing, cluster 21)[37, 63, 92]. Cluster 15 was composed of neurons defined by *RBFOX3* expression including *CUX2*-, *NRGN*-, *SLC17A7*-, and/or *TBR1*-expressing excitatory neuron subclusters and *GAD1*-, *SLC32A1*-, *SST*-, and/or *VIP*-expressing inhibitory neuron subclusters[63, 92, 93]. Immune cell clusters included a microglial supercluster (*PTPRC*-, *CD14*-, and *P2RY12*-expressing, clusters 1, 5, 11, and 17) and lymphocyte clusters (*PTPRC*-expressing but not *CD14*-expressing, clusters 7 and 28 with cluster 7 nuclei expressing T cell marker (CD3, CD4, and CD8)-encoding transcripts)[63, 92]. We further identified an endothelial cluster (*VWF*-, *PECAM1*-, and *CD34*-expressing, cluster 12) and vascular leptomeningeal cells (VLMCs, cluster 22, *DCN*-, *COL1A1*-, *COL1A2*-expressing)[19, 63, 92, 94]. After identifying brain stromal cells with high confidence, we turned our attention to the remaining putative neoplastic cell clusters.

The remaining clusters contained abnormal appearing macroglia- and neuronal-like cells (clusters 0, 2-4, 6, 8-10, 13, 14, 16, 18, 19, 24-27, and 29). These clusters contained abnormal combinations of markers (despite doublet removal during preprocessing, **Figure 1B and sFigure 1H**). For instance, cluster 6 cells tended to express *CD34* (an endothelial, hematopoietic stem, and neuroectodermal neural precursor cell marker), *RBFOX3* (normally a pan-neuron-specific marker), and *AQP4* (normally an astrocyte-specific marker)[12, 19, 63, 92, 94]. Among these putative neoplastic cells, cluster 6 was the most enriched in *CD34*. In terms of cell type proportions, 22932/34907=66% of cells were neoplastic-appearing. Of these neoplastic cells, 694/22932=3.0% were in cluster 6, 259/22932=1.1% were *CD34*+, and 168/22932=0.7% were cluster 6, *CD34*+ cells. Smaller proportions of *CD34*+ cells were present among clusters 24 (5% *CD34*+), 3 (1.6% *CD34*+), 19 (1.2% *CD34*+), 14 (0.9%*CD34*+), 10 (0.5% *CD34*+), 8 (0.4% *CD34*+), 13 (0.2% *CD34*+), and 0 (0.02% *CD34*+). All cluster 6 *CD34*+ cells (n=168) were from tumor 5. Outside of cluster 6, the neoplastic-appearing *CD34*+ cells were from tumor 1 (n=66 *CD34*+), tumor 5 (n=11 *CD34*+), and tumor 4 (n=10 *CD34*+). Hence, analysis of cell markers identified the majority of ganglioglioma cells as neoplastic-appearing with a rare *CD34*-rich population possibly representing tumor neuroectoderm neural precursor cell-like stem cells.

Preliminary normal and neoplastic cell identification was confirmed by analysis of top differentially expressed genes (**Supplementary Information** and **sTable 2**), inference of copy number variation, and Seurat-based label transfer. Inference of copy number variations using inferCNV[55] identified clonal del1p and subclonal del14 exclusively in the putative neoplastic cell clusters in tumor 4 (**sFigure 1I**), providing support that these clusters represented neoplastic cells. Note that most gangliogliomas do not carry large CNVs[23, 58], so the absence of such alterations for snRNA-seq cells from the other tumors does not rule out their assignment as neoplastic. We next mapped snRNA-seq profiles to transcriptional atlases from developing brain[14, 25, 34, 52, 91], adult brain[6, 49, 93], primary brain tumors[20, 67, 82], and tumor-associated immune cells[5, 18, 51, 90]. This mapping reinforced and refined stromal cell type assignments (examples in **Figure 1C** and **sFigure 1J-L**). Nuclei of the suspected neoplastic clusters generally had much more mixed and uncertain mapping, consistent with these representing abnormal, neoplastic cells. Interestingly, the *CD34*-rich cluster 6 nuclei were confidently identified as immature astrocytes where represented in reference atlases. For instance, cluster 6 mapped confidently to first-/second-trimester (i.e. immature) astrocytes in a developmental brain atlas (**Figure 1C**)[52], and cluster 6 was confidently identified as protoplasmic/immature astrocyte by mapping to a glioblastoma atlas (**sFigure 1L**)[82]. Other neoplastic-appearing clusters tended to map to either neurons or macroglia at different developmental stages. For instance, clusters 0, 4, 8, 9, 10, 18, and 19 mapped to first-/second-trimester neurons while cluster 13 mapped to developing OPCs and clusters 2, 16, and 29 mapped more closely to ventricular radial glia (vRG; **Figure 1C**)[52]. Thus, snRNA-seq results support the presence of large neuron-like and macroglia-like populations in addition to a smaller neuroectoderm neural precursor cell-like compartment among neoplastic GG cells.

### GG *CD34*-associated neuroectodermal neural precursor-like cells exhibit stem cell states

Many cancers, including gliomas, are composed of a cellular hierarchy containing stem-like cells capable of self-renewal, differentiation, tumorigenecity, tumor progression, or resilience in the face of anti-tumor therapy and ultimately tumor regrowth[50, 55, 61, 78]. Understanding these cells is of special interest because their identification could be used for better tumor classification, prognostication, and therapeutic targeting. Cell typing and mapping above identified a neoplastic, *CD34*-rich cluster 6. Such CD34+ cells have previously been hypothesized to represent ganglioglioma tumor precursor/stem cells, but their transcriptomic profile and location in the tumor cell hierarchy was not previously well understood.

Given the working hypothesis that gangliogliomas arise from neuroectodermal neural precursor-like cells, neoplastic cells were next interrogated for individual neuroectodermal markers in addition to *CD34*[80]. Interestingly, cluster 6 identity and *CD34* expression were each associated with expression of neuroectodermal neural precursor cell markers *PAX6*, *SOX2*, and *MSI1* (**Figure 2A**)[38, 72, 88]. Hence, analysis of cell markers supported our hypothesis that neoplastic *CD34*+ cells resemble primitive neuroectoderm neural progenitor cells.

**Figure 2:**
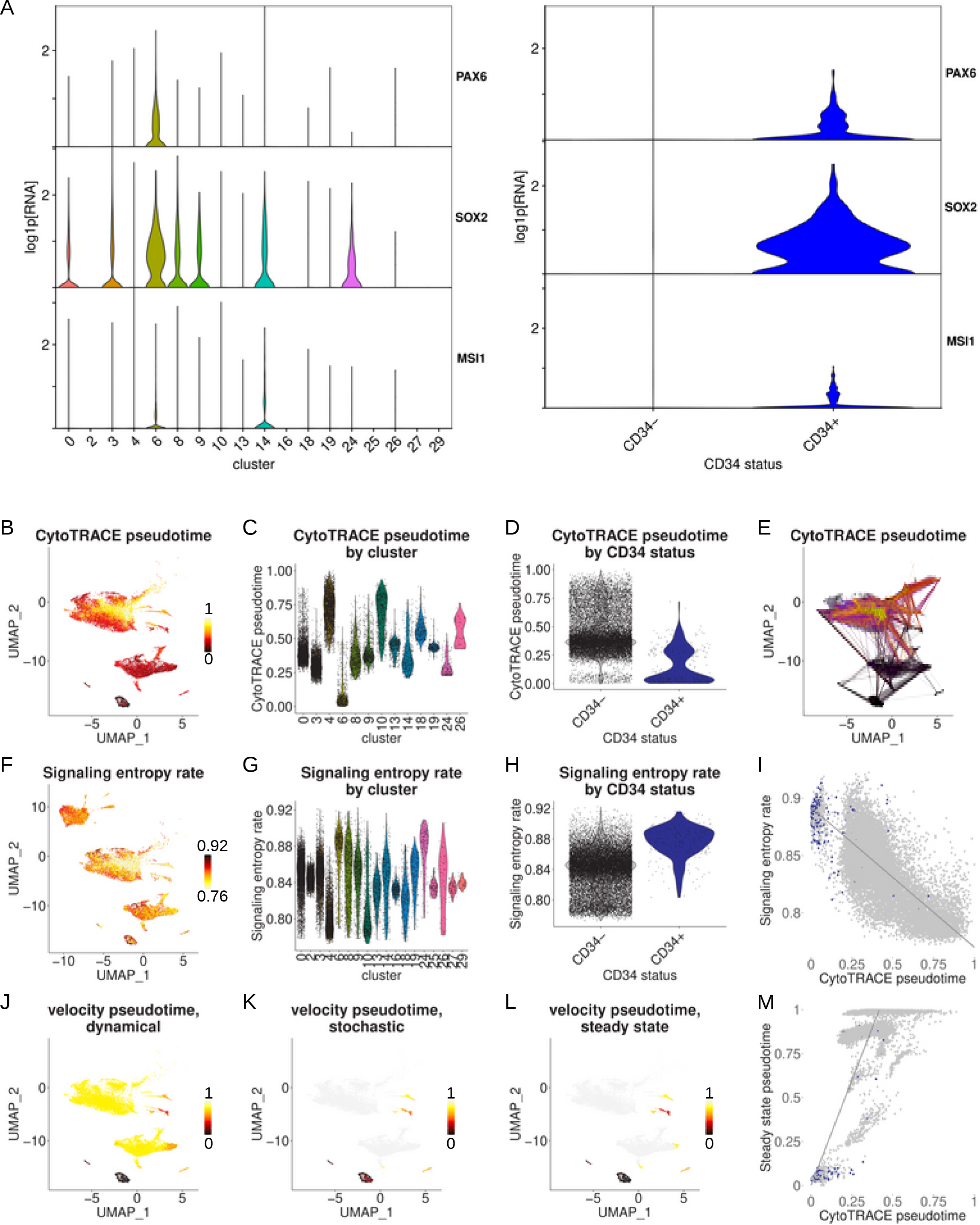
Ganglioglioma neoplastic cellular hierarchy. (A) Violin plots of neoplastic cell expression (log1p[RNA], base 2) of individual neuroectodermal markers by cluster (left) or by *CD34* status (right). (B) Feature plot of ganglioglioma neoplastic tumor 1, 4, and 5 nuclei CellRank/CytoTRACE pseudotime (tumor 3 nuclei excluded due to low complexity which resulted in aberrantly high calculated pseudotime for tumor 3 nuclei). (C-D) Violin plots of CytoTRACE pseudotime by ganglioglioma cluster (C) or *CD34* status (D). (E) Random walk of earliest pseudotime cells (black dots) via CellRank/CytoTRACE pseudotime to predicted terminal states (yellow dots). (F-H) Differentiation potency was also inferred via SCENT signaling entropy rate (SR) for ganglioglioma neoplastic nuclei. Feature and violin plots of SR shown analogous to B-D above for CytoTRACE pseudotime. (I) Signaling entropy rate versus CytoTRACE pseudotime for ganglioglioma neoplastic nuclei (except tumor 3), colored by *CD34* status (*CD34+* in blue, *CD34*-gray). Gray line represents linear least-squares fit to (all of the neoplastic) data, with gray shading representing the 95% confidence interval. (J-L) Neoplastic cell scVelo RNA velocity-derived pseudotimes by dynamical (J), stochastic (K), and steady state (L) modes. (M) RNA velocity steady state pseudotime vs CytoTRACE pseudotime, colored by *CD34* status (*CD34+* in blue, *CD34*-gray). Gray line represents linear least-squares fit to (all of the neoplastic) data, with gray shading representing the 95% confidence interval.

We next sought to identify where the neuroectoderm neural precursor-like cells fall within the ganglioglioma neoplastic cell developmental hierarchy. First, we used CellRank/CytoTRACE to infer pseudotime[31, 42]. Analysis of our neoplastic-appearing ganglioglioma nuclei in this manner assigned cluster 6 nuclei or *CD34*+ nuclei (and particularly cluster 6, *CD34*+ nuclei) the earliest pseudotimes (**Figure 2B-E**). We next used SCENT to calculate signaling entropy rates (SR), which closely approximate stemness[77]. This method largely recapitulated the results with CellRank/CytoTRACE (**Figure 2F-I**). Finally, we used RNA velocity to infer temporal relationships between cell states[9, 16]. Multiple RNA velocity methods consistently agreed that cluster 6 and/or *CD34*+ neoplastic cells appeared to be the most primordial of neoplastic cells (**Figure 2J-M**). Therefore, several orthogonal methods showed that neoplastic cells represented in cluster 6 and/or *CD34*+ appeared particularly potent (i.e. stem cell-like), consistent with these representing primitive, neoplastic stem/precursor cells.

To further pin-down stem cell-like states and possibly the cell type of origin for ganglioglioma, we combined analysis of neuroectodermal neural precursor markers and cellular hierarchy results. Co-expression of *CD34*, *PAX6*, *SOX2*, and *MSI1* (after stringent counter selection against vascular cells using *PECAM1*, *VWF*, *DCN*, and *COL1A2*), was particularly strongly associated with ganglioglioma neoplastic cell stemness (**sFigure 2A**). To evaluate the timing of disappearance of such cell states during normal development, we tested existing single cell brain atlases for the presence of *CD34+PAX6+SOX2+MSI1+PECAM1-VWF-DCN-COL1A2-* cells. We first noted that prenatal brain reference atlases we probed all contained small but appreciable numbers of these cells: 14/2394 in Zhong *et al*.[91], 10/4129 in Nowakowski *et al*.[52], 7/25161 in van Bruggen *et al*.[14], and 9/48215 in Eze *et al*.[25] Considering data drop out, the actual potent precursor cells of the intended type are likely somewhat more prevalent. In contrast, adult brain reference atlases had no *CD34+PAX6+SOX2+MSI1+PECAM1-VWF-DCN-COL1A2-* cells detected: E.g. 0/47432 in the Allen Institute multiple cortical areas atlas[93] and 0/78886 cells in Nagy *et al*.[49] Interestingly, a very large, recently published brain single cell atlas spanning the third trimester to 40 years of age helped to fill the gaps in developmental data during childhood and adolescence[34]; 0 out of 154748 cells in the Herring *et al*. atlas were *CD34+PAX6+SOX2+MSI1+PECAM1-VWF-DCN-COL1A2-.* A potential confounder is the possible spatial variability of this cell type. However, when limiting our analysis to the pre-frontal cortex (due to there being robust data spanning development for this region), we still found an apparent disappearance of *CD34+PAX6+SOX2+MSI1+PECAM1-VWF-DCN-COL1A2-* cells during fetal development. Prenatal pre-frontal cortex *CD34+PAX6+SOX2+MSI1+PECAM1-VWF-DCN-COL1A2-* cells were found by Nowakowski *et al*. (2 cells out of 1076)[52] and Zhong *et al*. (14 cells out of 2394)[91]. No such cells were detected in the pre-frontal cortex third trimester to 40 years old (0/154748)[34] or a purely adult pre-frontal cortex atlas (0/78886)[49]. Hence, these results overall support the disappearance of the neuroectodermal neural precursor cells that ganglioglioma stem-like cells most resemble during normal brain development (fetal development for the pre-frontal cortex) and suggest that the ganglioglioma cell of origin may in fact arise during fetal brain development.

### Ligand-receptor analysis reveals PTN-PTPRZ1, FGF family, and PDGF family communication among ganglioglioma neoplastic cells

Intercellular communication pathways were interrogated in an unbiased manner using CellChat[35] (**sFigure 3A-B, sTables 3-5**). This analysis suggested neoplastic tumor cell PTPRZ1 targeting by PTN, the latter largely produced by the neoplastic cells (**Figure 3A and sFigure 3C-D**). 57% of neoplastic cluster-neoplastic cluster pairs exhibited a PTN-PTPRZ1 interaction (p-value <0.01). Every neoplastic cluster had at least one cluster by which it was targeted (p-value<0.01). Cluster 6 was particularly active in signaling, as a source for 17/18 neoplastic clusters (p<0.01, all except cluster 18) and as a target from 10/18 neoplastic clusters (p<0.01). PTPRZ1 is a receptor tyrosine phosphoprotein phosphatase[86]. PTN binding antagonizes PTPRZ1, which has been shown to have pleiotropic effects including activation of MAPK and AKT (PKB) signaling axes[86]. A mouse model of BRAF-V600E neuroectodermal tumors required a second hit with increased AKT/mTOR signaling to produce ganglioglioma-like tumors[17]. These results identify a neoplastic-cell-to-neoplastic-cell interaction that is known to drive the signaling axes needed for gangliogliomagenesis as a major feature of ganglioglioma in general, and of ganglioglioma cluster 6 stem/neuroectoderm neural precursor-like cells in particular.

**Figure 3:**
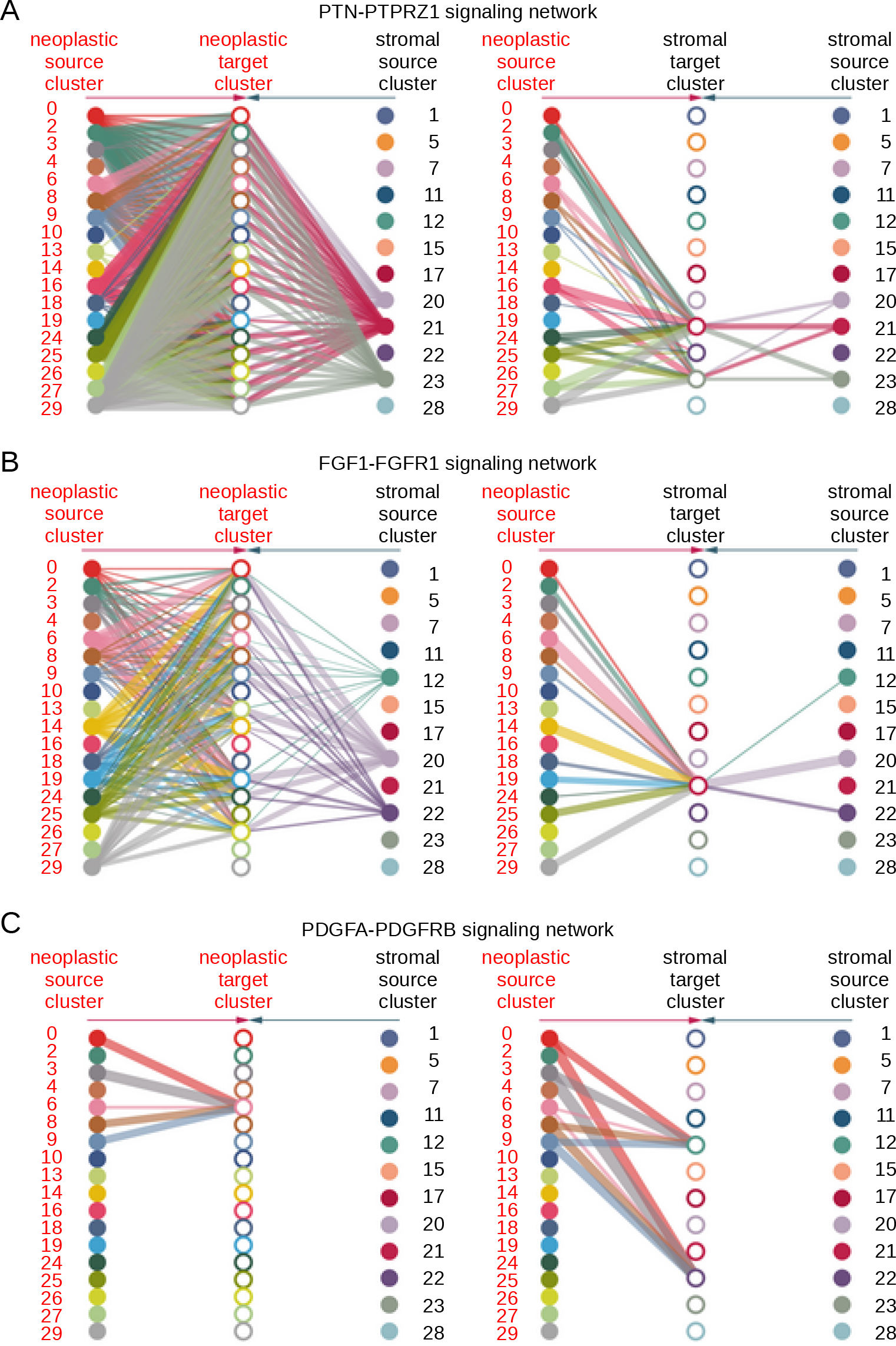
Ganglioglioma neoplastic cell communications of interest by snRNA-seq evaluated with CellChat. (A) Significant PTN-PTPRZ1 interaction as nominated by CellChat with results shown as hierarchy plots. Line thickness reflects strength of observed interaction. Interactions targeting neoplastic cells on the left plot and interactions targeting stromal cells on the right plot. Note that within each plot, neoplastic cluster sources are on the left and stromal cluster sources are on the right. Neoplastic targets are in the same order as neoplastic sources in the left plot, and stromal targets are in the same order as stromal sources in the right plot. (B-C) Analogous to (A) except for FGF1-FGFR1 (B) and PGFA-PDGFRB (C).

We also localized neoplastic cell compartments that take part in FGF and PDGF family ligand-receptor interactions. FGFR alteration is associated with a substantial minority of pediatric low grade gliomas[23, 58, 68]. PDGF(R) aberrant hyperactivity is potently oncogenic, more typically associated with high grade gliomas[65]. FGF1-FGFR1 interactions occurred between most clusters of neoplastic cells or OPCs (p<0.01 for 37% of possible neoplastic cluster-neoplastic cluster interactions, cluster 6 as a source p<0.01 for 10/18 neoplastic clusters including itself, cluster 6 as a target p<0.01 for 12/18 neoplastic clusters; **Figure 3B** and **sTables 4-5**). Other FGF-FGFR family members exhibited highly significant interactions between multiple neoplastic clusters (**sFigure 3E-I** and **sTables 4-5**). The PDGF family pathway was also a top hit (**sTable 3**). For instance, we identified interactions between PDGFA-PDGFRB in cluster 6 (p<0.01 for cluster 6 targeting by 5/18 neoplastic clusters, including itself, with no other significant neoplastic cluster-neoplastic cluster interaction (p=1)) as well as endothelial cluster 12 and VLMC cluster 22 (**Figure 3C** and **sTables 4-5**). We also detected PDGFB-PDGFRB and PDGFD-PDGFRB targeting directed at cluster 6 and vascular cells (**sFigure 3J-K** and **sTables 4-5**). These results pinpoint ganglioglioma neoplastic cell states that take part in FGF and PDGF signaling interactions, and reveal that such processes are especially active in neuroectoderm neural progenitor cell-like cluster 6 cells.

### BRAF and AKT pathway signaling in neoplastic cells

We next sought to identify signaling programs that may function downstream of the ligand-receptor interactions to drive growth and maintenance of the neoplastic cells. To ascertain important cellular signaling pathways in ganglioglioma, neural cells were subject to gene set enrichment analysis (GSEA) using clusterProfiler for gene ontology (GO), KEGG, and Wiki pathways[75, 85].

To uncover neoplastic cluster-associated pathway suppression or activation, neoplastic-appearing ganglioglioma clusters were compared to normal-appearing neural clusters (clusters 15, 20, 21, and 23). Consistently among the most depleted pathways (in neoplastic cells) were various pathways responsible for oxidative phosphorylation or ribosomal function, whether comparing by individual cluster or using neoplastic-appearing cells altogether (**Figure 4A and sFigure 4A-F**). Oxidative metabolism suppression is consistent with the Warburg effect typical of neoplastic cells and offers further support for our identification of cells as neoplastic or stromal[83].

**Figure 4:**
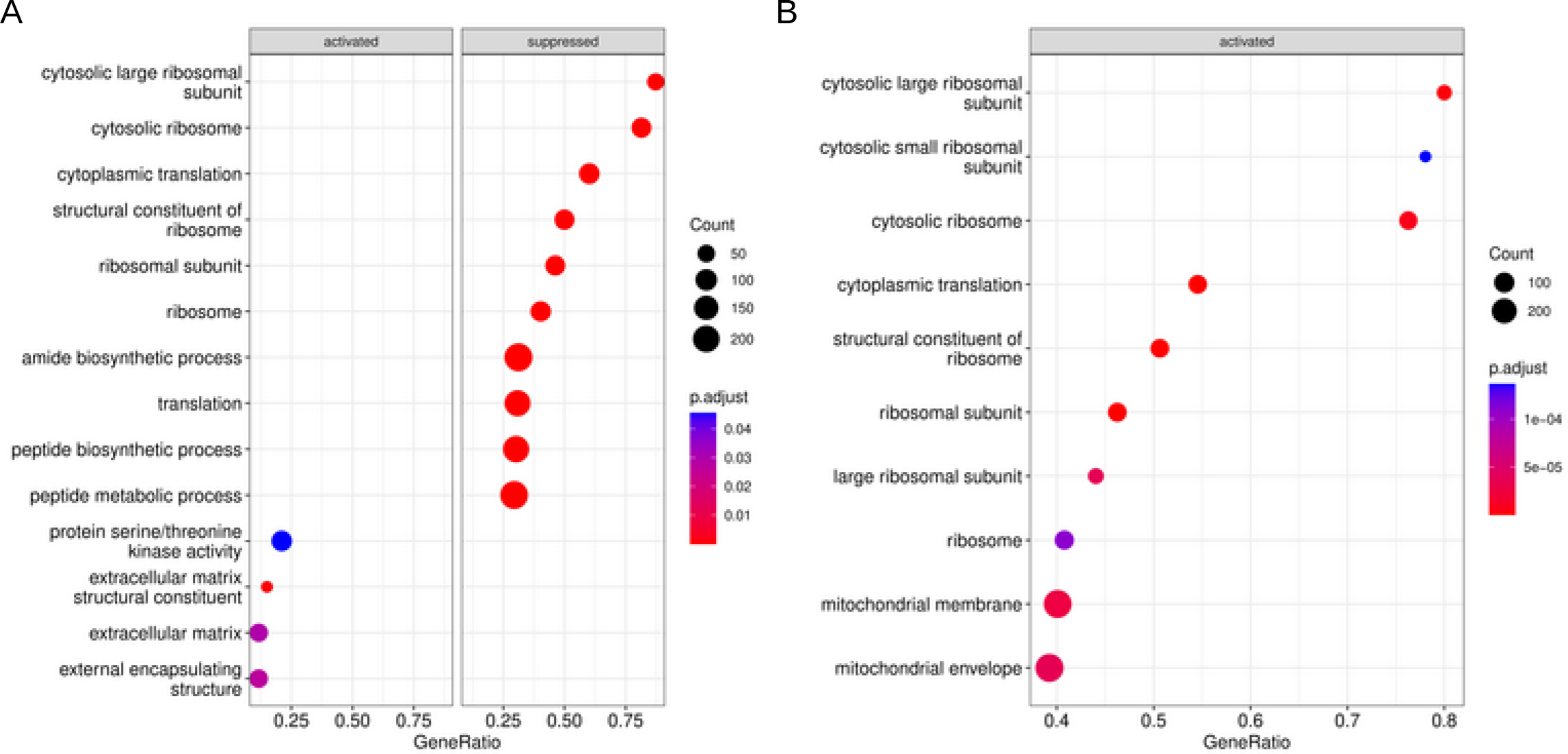
Ganglioglioma snRNA-seq gene set enrichment analysis. (A) Dotplots of top 10 (or fewer if Bonferroni-adjusted p-value>0.05) most activated or suppressed GO pathways when comparing neoplastic-appearing cells as a whole to normal-appearing neural cells. Bonferroni-adjusted-p-value shown as well as gene count in each pathway. (B) Dotplots of top 10 (or fewer if Bonferroni-adjusted p-value>0.05) most activated or suppressed GO pathways when comparing *CD34*+ neoplastic-appearing cells as a whole to *CD34*- neoplastic-appearing cells. Bonferroni-adjusted-p-value shown as well as gene count in each pathway.

In order to ascertain the pathways significant to *CD34*+ neoplastic cells in particular, we compared neoplastic *CD34*+ cells to neoplastic *CD34*-cells. The most *CD34*+ cell-enriched pathways were those involving oxidative phosphorylation and ribosomal machinery (**Figure 4B and sFigure 4G-L**). Unsurprisingly given neoplastic *CD34*+ cells’ promiscuous use of signaling nodes, they had very little in the way of pathway suppression (recall the high signaling entropy rate discussed above, **Figure 4B**, and **sFigure 4G-L**). Nearly identical results were obtained when comparing cluster 6 *CD34*+ cells to the remaining neoplastic-appearing cells. Close inspection of KEGG pathways of interest showed especially brisk expression of the PKB/AKT pathway machinery from FGF through downstream AKT effectors in neoplastic *CD34*+ versus neoplastic *CD34*-cells (**sFigure 4K**). Additionally, there was generally higher expression of BRAF pathway machinery-encoding transcripts in neoplastic *CD34*+ cells, including *BRAF* itself (**sFigure 4K**). Pluripotency machinery-encoding transcripts were generally greatly enriched in the neoplastic *CD34*+ cells, including the transcription factor-encoding *SOX2* (**sFigure 4L**). SOX2 is a SOX B1 member which contributes to embryonic, neural stem, and progenitor cell self-renewal and pluripotency (very early)[72]. These results were overall congruous with the preceding analysis. Particularly interesting is the persistent theme of signaling via AKT in addition to BRAF, particularly in the most stem-like of the neoplastic cells, the neoplastic *CD34*+ and/or cluster 6 cells. For the AKT pathway, there may very well be contribution by autocrine signaling via FGFs and/or PTN.

### Neoplastic cell gene regulatory networks and putative drivers

We next characterized transcription factors that may account for compartmental expression of the gene programs identified above. To do so, gene regulatory networks were reconstructed in an unbiased manner using SCENIC[69]. Top candidate transcription factors for both cluster 6 and neoplastic *CD34*+ cells included paired box 6 (PAX6) and its transcriptional activators myeloid ectopic viral integration site homeobox 1 (MEIS1) and transcription factor 7 like 2 (TCF7L2; **sFigure 5A-D**)[53, 57]. In the healthy adult brain, these transcription factors are not normally coexpressed[63, 92]. More interestingly, PAX6 is a neuroectodermal neural precursor cell marker, and the MEIS1-PAX6 cascade appears critical to early neuroectodermal cell fate determination[25, 53, 88]. In addition, TCF7L2 coordinates PAX6 activation in neural cells and modulates MYC as part of the WNT pathway[57]. *PAX6*, *MEIS1*, and *TCF7L2* concentrations as well as SCENIC-calculated regulon scores correlated strongly with ganglioglioma neoplastic cell stemness by CellRank/CytoTRACE pseudotime or SCENT signaling entropy rate (**Figure 5A** and **sFigure 5E-F**). SOX2 also appeared as a factor with significantly increased activity (and transcript concentration) among certain neoplastic cells, particularly those most stem-like, including cluster 6 and/or *CD34*+ cells (**Figure 5B**). SOX2 contributes to embryonic, neural stem, and progenitor cell self-renewal and pluripotency (very early, including in the neuroectoderm) and is not normally appreciably expressed in the adult brain[63, 72, 92]. These observations implicate PAX6, MEIS1, TCF7L2 and SOX2 programs in the more primitive, neuroectoderm neural precursor-like (cluster 6 and/or *CD34*+) tumor cells, and these programs are possibly controlling tumor cell potency/stemness/differentiation.

**Figure 5:**
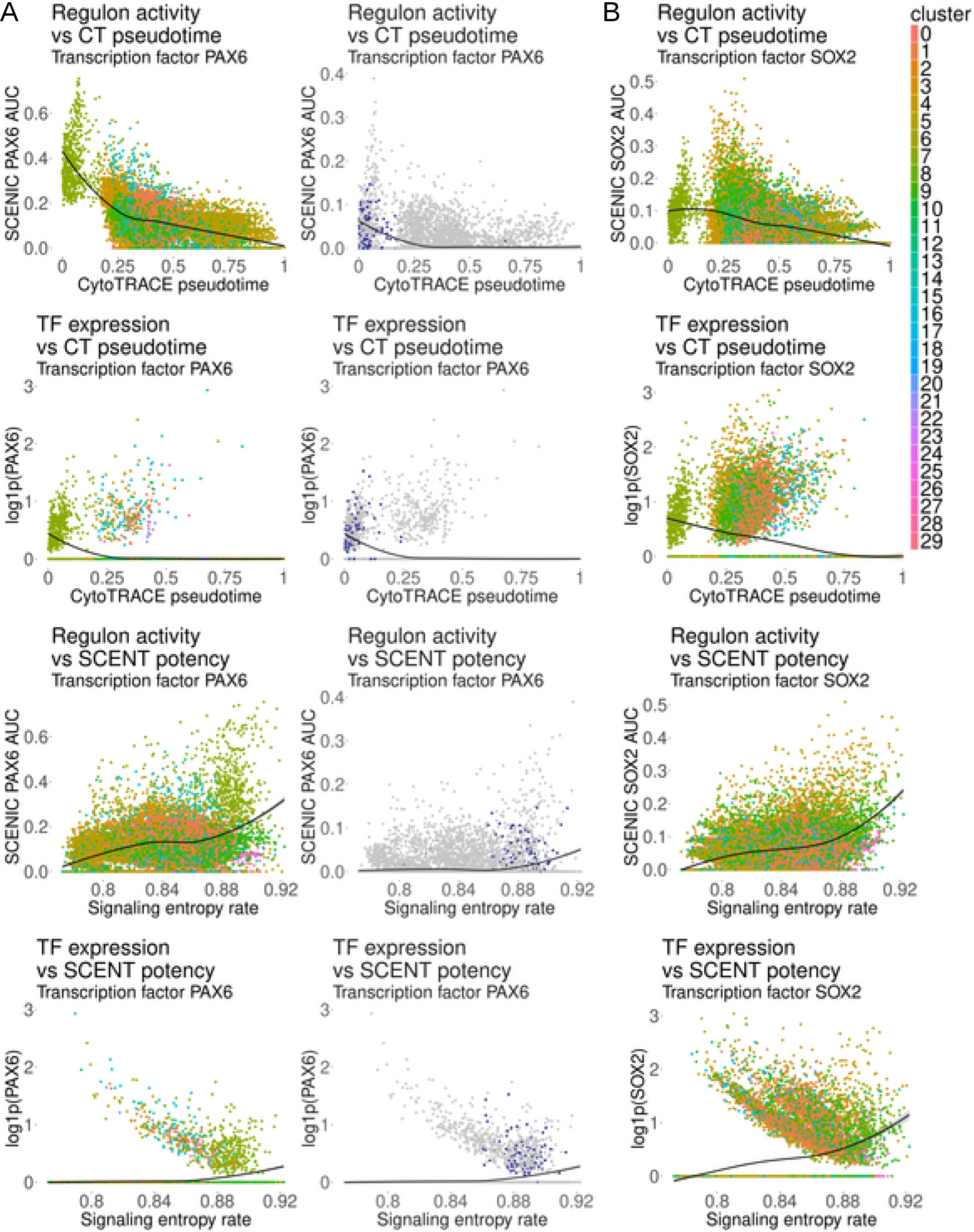
Ganglioglioma significant transcription factors. (A) Left: Scatterplots of PAX6 cluster-based SCENIC AUC and transcription factor expression vs CytoTRACE pseudotime or SCENT signaling entropy rate (SR) with data colored by cluster (legend to the topright of the figure). Loess best fit shown as a black curve; gray shading for the 95% confidence interval. Right: *CD34* status-based SCENIC AUC and transcription factor expression vs CytoTRACE pseudotime or SCENT signaling entropy rate (SR) with data colored by *CD34* status (*CD34*+ blue, *CD34*-gray). Loess best fit shown as a black curve; gray shading for the 95% confidence interval. (B) Analogous results to (A) are shown for SOX2 (cluster-based results only due to *CD34*-based regulon not meeting filters).

### Immune landscape

#### Myeloid cells

In healthy brain, myeloid cells are predominantly resident microglia; however, others have established the presence of bone marrow derived macrophages associated with brain tumors[48]. To determine the nature of immune cells in gangliogliomas, we first used markers of *a priori* interest to further decipher cell states. Ganglioglioma clusters 1, 5, 11, and 17 were *PTPRC*-, *ITGAM*-, *CD14*-, and *P2RY12*-expressing consistent with these representing microglia (**Figure 6A** and recall **Figure 1B** and **sFigure 1H**)[1, 63, 92]. Myeloid cells lacked appreciable expression of the classical pro-inflammatory markers *IL1A*, *IL6*, *TNF*, and *CD40* (all myeloid cells *IL6*-; otherwise **Figure 6A**)[5]. In terms of anti-inflammatory activity, they did not express appreciable levels of *IL10*, but there was diffuse expression of *TGFB1* (**Figure 6A**)[5]. Moreover, when looking at the broader cellular context, lymphocytes from clusters 7 and 28 expressed significant *TGFB1* and tumor clusters 3, 8, and 14 expressed substantial *CSF1*, which promote immunosuppressive and pro-inflammatory properties in macrophages, respectively (**Supplementary Information** and **sTable 2**)[5]. Clusters 1, 5, 11, and 17 all appeared to express significant amounts of *CD163* (except cluster 11) and *MSR1* (encodes CD204) and variable *MRC1* (encodes CD206), classic markers of immunosuppressive/tissue reparative/tumor promoting M2 macrophages (**Figure 6A**)[5]. In terms of classic pro-inflammatory M1 markers, myeloid cells expressed *TLR2* highly, *CD86*, variable *HLA-DR* (most in cluster 17), and very little *NOS2* (**Figure 6A**)[5]. This co-expression of M1 and M2 markers appears consistent with prior observations suggesting that the M1 versus M2 dichotomy does not seem to hold on the single cell level in tumor microenvironments[5]. Instead, M1 and M2 traits tend to be strongly positively co-varying in individual tumor-associated cells[5, 48]. More recently, with the help of single cell typing of tumor-associated immune cells, it has been suggested that C1Q and SPP1 status best differentiate different types of tumor-associated macrophages and that these markers are also associated with prognosis[43, 87]. C1Q expression appears to be associated with T lymphocyte recruitment and activation whereas SPP1 expression appears to be associated with tumor growth and metastasis. *C1Q* expression was variable in ganglioglioma myeloid cells, with high concentrations in cluster 17 and consistently elevated concentrations in cluster 11 (**Figure 6A**). *C1Q*+ cells were not distinct from *SPP1*+ cells, with *SPP1* expression high in cluster 11 and parts of clusters 17 and 5 (**Figure 6A**). These results overall support the presence of tumor-associated, aberrantly activated microglia.

**Figure 6:**
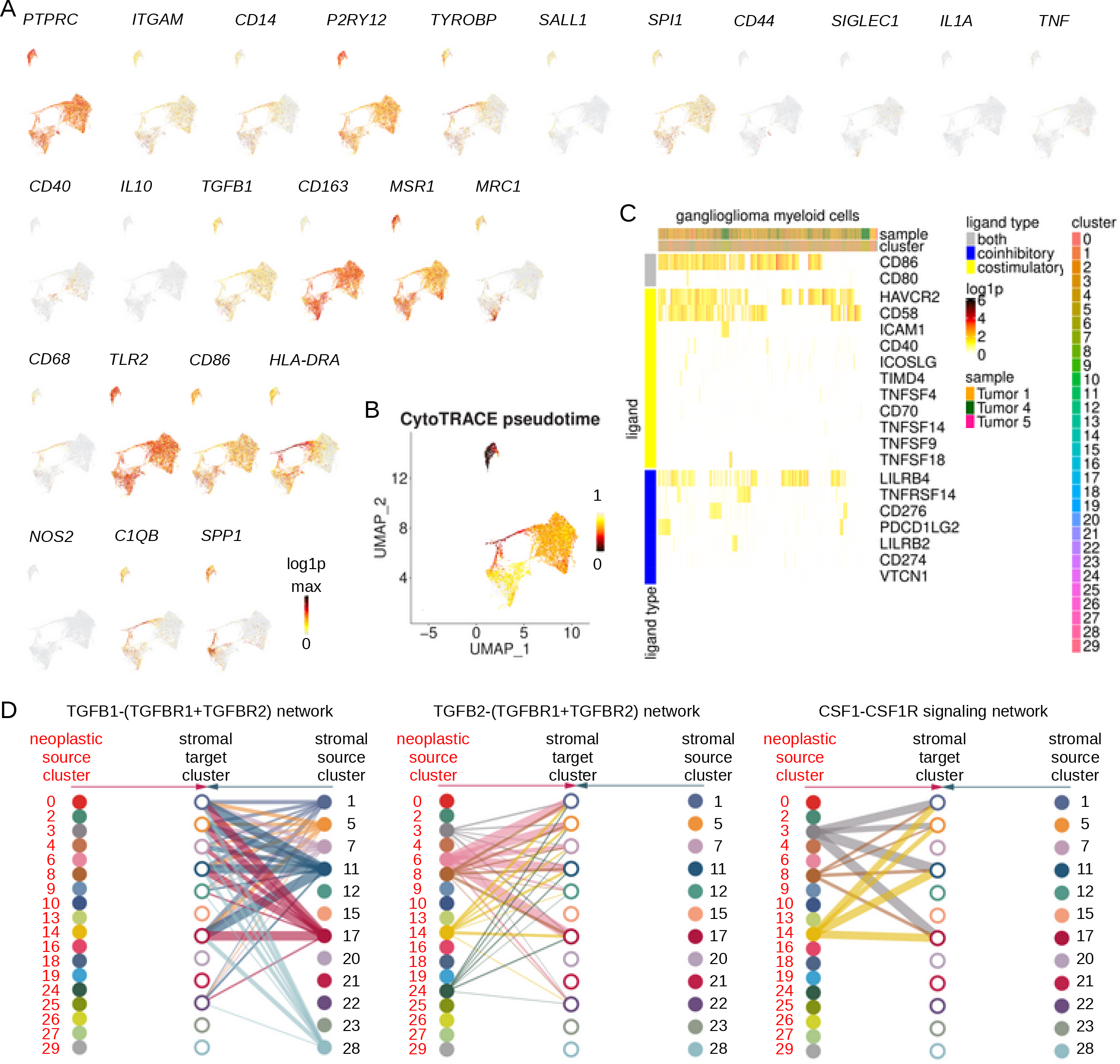
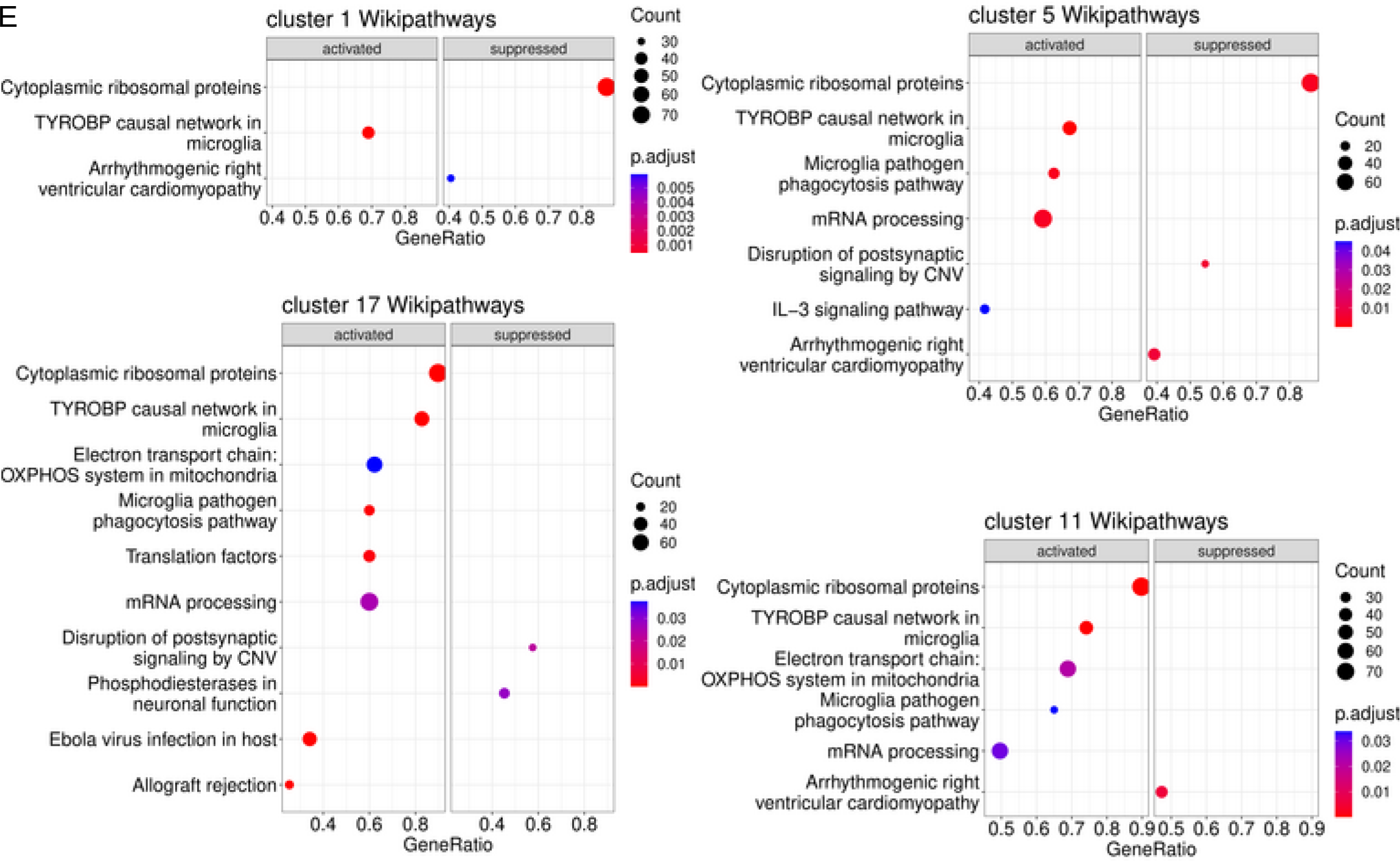
Ganglioglioma-associated myeloid cells. (A) Myeloid supercluster feature plots of RNA concentrations (log1p, base 2) for key general myeloid (*PTPRC*, *ITGAM*, *CD14*), microglia (*P2RY12*, *TYROBP*, *SALL1*, *SPI1*), macrophage (*CD44*, *SIGLEC1*), classical proinflammatory (*IL1A*, *TNF*, *CD40*; no expression for *IL6*, so not shown), anti-inflammatory (*IL10*, *TGFB1*), tissue reparative M2 (*CD163*, *MSR1*, *MRC1*, *CD68*), and M1 proinflammatory (*TLR2*, *CD86*, *HLA-DRA*, *NOS2*) markers as well as *C1QB* and *SPP1*. (B) Microglia feature plot of cellular hierarchy by CellRank/CytoTRACE pseudotime. (C) Heatmap of microglia expression of transcripts encoding lymphocyte stimulatory and inhibitory ligands of interest *a priori*. (D) Hierarchy plots of CellChat results showing significant involvement of microglia in TGFB and CSF pathways. Lines weighted by interaction significance. (E) Most (top 10 or fewer if Bonferroni-adjusted-p-value>0.05) activated or suppressed Wikipathways compared to the bulk ganglioglioma population for each of the microglial clusters.

We attempted to use differential gene expression analysis, multi-gene signature scoring, and Seurat-based label transfer to further type these cells. Top microglial cluster-overexpressed genes were enriched for complement-encoding transcripts (particularly C1Qs) and MHC class II-encoding transcripts, consistent with their expected roles in the complement cascade and professional antigen presentation, respectively (**sFigure 6A**). To confirm ganglioglioma myeloid cells represented microglia, Seurat-based label transfer was performed[32]. Using annotated tumor reference atlases differentiating microglia from macrophages[67, 82] for label transfer, the bulk of myeloid cells were consistently more strongly identified with an overall microglial rather than macrophage signature (recall **sFigure 1K-L**). When ganglioglioma-associated myeloid cells were subjected to multi-gene signature scoring (by UCell) based on established M1 and M2 signatures[5], the two scores appeared positively correlated both within each cluster and among cells of the supercluster, consistent with the aberrant activation that was found at the individual marker level and further supporting the lack of this dichotomy in the context of some tumors (**sFigure 6B-D**). Overall, differential gene expression analysis, multi-gene signature scoring, and Seurat-based label transfer confirmed ganglioglioma-associated microglial cell identity and refined their cell states.

Ganglioglioma-associated myeloid cellular hierarchy was evaluated with CellRank/CytoTRACE, SCENT signaling entropy rate, and scVelo RNA velocity[9, 31, 42, 77]. These generally identified cluster 11 and 17 members of greater potency/stemness and therefore candidates for tumor-associated myeloid precursor states (e.g., **Figure 6B**).

Myeloid cell interactions were explored. Interestingly, myeloid cell-expressed concentrations of transcripts encoding lymphocyte co-stimulatory and co-inhibitory ligands appeared positively correlated throughout the supercluster (**Figure 6C**). Among the top ranked pathways in unbiased ligand-receptor analysis by CellChat and CellPhoneDB[24, 35], microglia participated in significant antigen presentation, including as senders and receivers of the MHC class II pathway, senders for the MHC class I pathway (clusters 5, 11, 17), and receivers for the APP-CD74 signaling pathway (MHC class II antigen processing, recall **sFigure 3A-B** and **sTable 3-5**). On the other hand, microglia also participated heavily in TGFB signaling, as senders and receivers, and they were receivers for CSF1-CSF1R (from neoplastic clusters 3, 14, and 8), consistent with a classically M2-promoting milieu and activities (**Figure 6D,** recall **sFigure 3A-B** and **sTable 3-5**)[5]. Hence, ganglioglioma-associated microglial cell interactions appeared along a spectrum of aberrant activation, consistent with what was ascertained about myeloid cell states earlier.

To ascertain important cellular signaling pathways in ganglioglioma-associated myeloid cells, microglia were subject to gene set enrichment analysis (GSEA) using clusterProfiler[85]. Among Wikipathways, the TYROBP causal network in microglia was among the top 2 activated pathways for each microglial cluster (**Figure 6E**). The microglia pathogen phagocytosis pathway was also among the top 2 activated pathways for clusters 5 and 17. Top activated GO pathways for clusters 11 and 17 were pathways related to the ribosome and translation whereas those for clusters 1 and 5 included B cell activation, regulation of immune response, leukocyte differentiation, lymphocyte activation, and innate immune response among other immune cell activation/differentiation pathways. These findings overall support the assignment of these cells as microglia and show a spectrum of pathway utilization among the microglia particularly as it pertains to modulation of immune response.

### Lymphocytes

Understanding tumor-infiltrating lymphocytes has become of great basic, translational, and clinical interest of late[54]. To explore ganglioglioma tumor-infiltrating lymphocytes, we started by interrogating for lymphocyte markers. Ganglioglioma cluster 7 was largely *PTPRC*-, *CD2*-, and *CD3*-expressing (**Figure 7A**), consistent with T lymphocyte identity[63, 92]. Cytotoxic T lymphocyte (CTL) activity is central to anti-tumor immune response[54]. Cells represented in the upper and right side of cluster 7 UMAP expressed the classic *CD2+CD3+CD4-CD8+* cytotoxic T lymphocyte markers including granzyme- and perforin-encoding RNA (**Figure 7A**), consistent with these cells representing cytotoxic T lymphocytes[63, 92]. On the other hand, the remaining cluster 7 cells expressed *CD4* but not *CD8* or other cytotoxic T lymphocyte-specific markers (**Figure 7A**), consistent with these representing helper T lymphocytes[63, 92]. Cluster 28 was *PTPRC*-expressing but negative for *CD2* and *CD3* (**Figure 7A**). On the other hand, this cluster had expression of B cell markers *CD19*, *CD20* (*MS4A1*), *CD22*, and immunoglobulin components (**Figure 7A**), consistent with B lymphocyte identity[63, 92]. Hence, interrogation for lymphocyte markers revealed two ganglioglioma-infiltrating lymphocyte clusters, T lymphocytes in cluster 7 and B lymphocytes in cluster 28. This was further supported by top cluster-specific differentially expressed transcripts (**sFigure 7A** and recall **sTable 2**) and Seurat-based label transfer (**sFigure 7B**).

**Figure 7:**
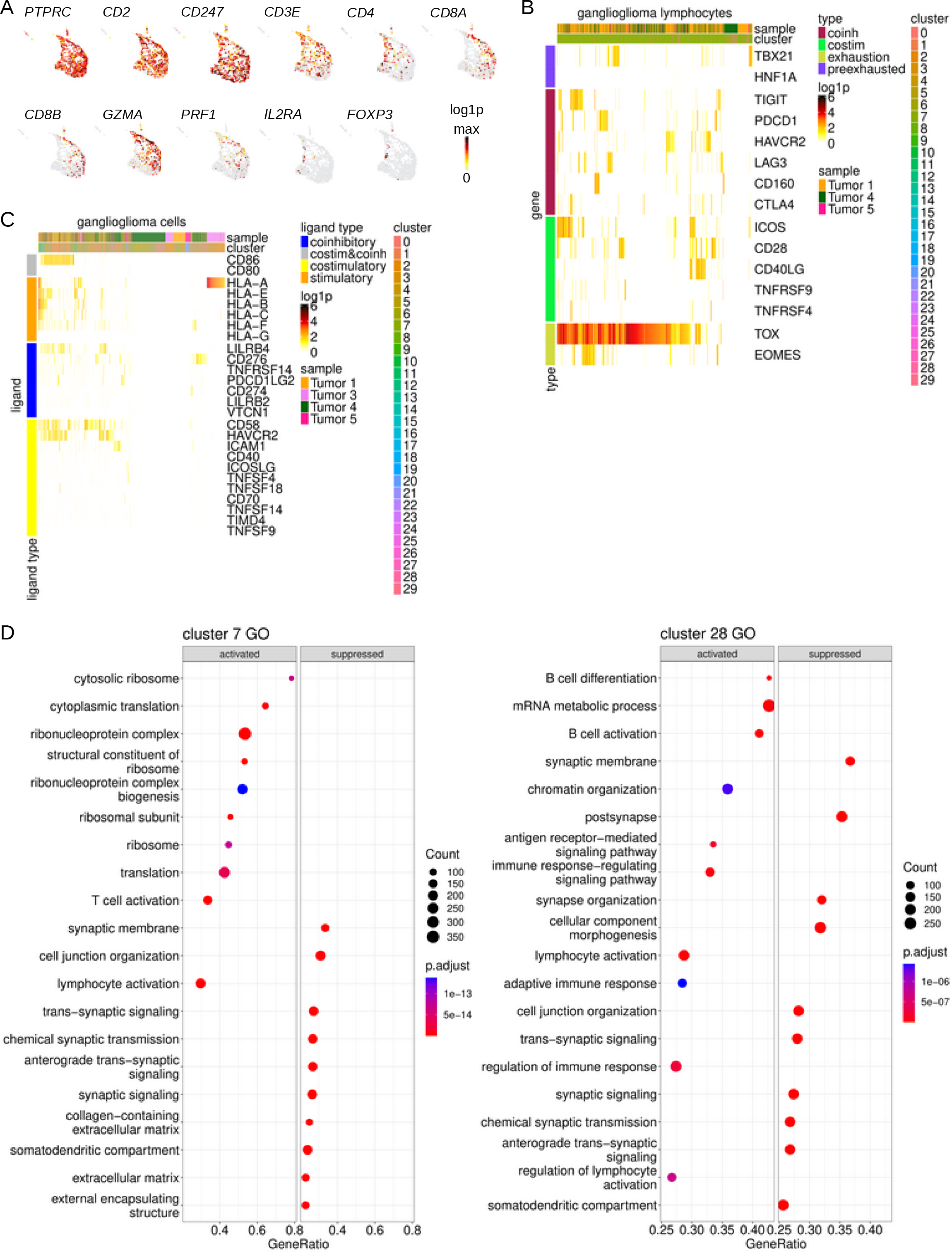
Ganglioglioma-infiltrating lymphocytes. (A) Lymphocyte feature plots of RNA concentrations for key general T lymphocyte (*PTPRC*, *CD2*, *CD3* (*CD3E*, *CD247*)), T helper (*CD4*), cytotoxic T lymphocyte (*CD8* (*CD8A* and *CD8B*), *GZMA*, *PRF1*), and regulatory T (*IL2RA*, *FOXP3*) markers. (B) Heatmap of lymphocyte RNA concentrations (log1p, base 2) for important markers of cytotoxic T lymphocyte state. (C) Heatmap of ganglioglioma cell expression of *a priori* important T cell modulating ligands. (D) Most (top 10 or fewer if Bonferroni-adjusted-p-value>0.05) activated or suppressed GO pathways for lymphocyte clusters.

To fine tune our understanding of tumor-infiltrating lymphocyte cell states, we explored cell markers further. The cluster 7 CTL subpopulation expressed a mixture of co-inhibitory receptor-encoding transcripts *CTLA4*, *PD1*, *TIGIT*, *LAG3*, *TIM3*, and *CD160* and co-stimulatory receptor-encoding RNAs *CD28*, *ICOS*, and *CD40LG* (**Figure 7B**)[5, 63, 92]. They had higher expression of exhaustion-related transcription factor-encoding RNAs *EOMES* and *TOX* than the precursor or pre-exhaustion factors *TBX21* and *HNF1A* (**Figure 7B**)[5, 63, 92]. These observations altogether suggest some degree of CTL dysfunction and exhaustion.

In addition to interrogation of individual markers, cell states were determined by signature score (rank-based, by UCell) based on subtype signatures previously evaluated in a single cell context[5]. Similar to what was observed for myeloid cell M1 versus M2 state, ganglioglioma-associated T lymphocyte M1 polarizing signatures and M2 polarizing signatures[5] were actually positively correlated (**sFigure 7C-D**). Signature scoring also identified a G2/M subpopulation of lymphocytes in cluster 7 (**sFigure 7E**)[5]. UCell scores were significantly higher for the Azizi *et al*. CD8 T cell activation, pro-inflammatory, and cytolytic effector pathway signatures[5] for those cells previously identified as enriched for CTL markers (**sFigure 7E**), confirming this assessment. In sum, tumor-infiltrating lymphocyte signature-scoring confirmed and refined preliminary cell state assignments based on individual markers.

We next turned to lymphocyte intercellular interactions because they constitute important determinants of T lymphocyte anti-tumor and anti-inflammatory roles. First, we analyzed expression of ligands (from the perspective of T cells) of interest *a priori*. MHC class I molecules allow presentation of a cell’s peptides to a CTL. Such presentation allows for CTL-mediated killing of tumor cells expressing neoantigens. Across many tumor types, neoplastic cell MHC class I expression is frequently downregulated as a means of escape from CTL-mediated tumor cell killing[22]. Tumor MHC class I expression has, in turn, been associated with prognosis and prediction of response to immunotherapy[30]. Tumor-associated myeloid cell MHC class I expression also has the potential to modulate the balance between immunologically “hot” and “cold” tumor microenvironment[13]. For our ganglioglioma cells, MHC class I component expression was particularly robust among microglia (**Figure 7C**). Additionally, neoplastic cell expression of MHC class I machinery-encoding transcripts was highly variable, with only subpopulations (highly-)expressing (**Figure 7C**). These results are consistent with those reported to date on ganglioglioma; others have found MHC class I upregulation among ganglioglioma-associated microglia and neoplastic neuron-like cells[64]. Altogether, these results suggest that (at least) the foundation for anti-tumor CTL activity exists in ganglioglioma.

To evaluate further, we interrogated for expression of ligands for CTL-bound receptors that modulate CTL activity in the context of successful MHC class I binding by T cell receptor. Some transcripts encoding co-stimulatory or co-inhibitory ligands, *CD86*, *LILRB4*, *CD58*, and *HAVCR2* were robustly expressed across the microglial supercluster, but other such ligands were generally scarce (**Figure 7C**)[5, 63, 92]. These results suggest opportunity, with some degree of priming capacity in the ganglioglioma-associated CTL milieu as well as potentially targetable immune checkpoints.

Beyond these core CTL-modulating ligand-receptor interactions, ganglioglioma-associated lymphocytes appeared to have relatively little in the way of ligand-receptor interaction. In unbiased ligand-receptor interaction nomination by CellChat[35], the lymphocyte clusters were among the least interactive clusters in terms of both incoming and outgoing signals (**sFigure 7F** and recall **sFigure 3A-B**). These findings may reflect a relatively immunologically “cold” tumor microenvironment and/or the relatively intensive nature of ganglioglioma neoplastic cell intercellular signaling.

In terms of cellular signaling, cluster-based GSEA[85] revealed cluster 7 top enriched pathways to include T cell activation (from GO); T cell receptor signaling (from KEGG); and TCR signaling, modulators of TCR signaling and T cell activation (WP; **Figure 7D and sFigure 7G-H**). Cluster 28 top enriched pathways included B cell receptor signaling, B cell differentiation, B cell activation, antigen-mediated signaling, immune response-regulating, lymphocyte activation, regulation of immune response, and regulation of lymphocyte activation (GO); BCR signaling and NF-κB signaling (KEGG); and BCR signaling (WP; **Figure 7D and sFigure 7G, I**). Unbiased transcription factor nomination by SCENIC[69] concurred with the spectrum of T cell activation among cluster 7 cells and B cell activation among cluster 28 cells (**sFigure 7J-K**). Altogether, these findings further confirmed preliminary lymphocyte typing and state.

### Protein-level validation with CITE-seq

Integrated proteogenomic analyses have suggested that transcript concentrations generally correlate with protein (i.e. functional end product in which we are typically most interested) concentrations. However, complex cellular feedback loops may at least theoretically allow for incongruous or even anticorrelated transcript and protein levels.

To address this issue, we used CITE-seq. CITE-seq allows simultaneous probing for epitopes of interest with transcriptomic profiling. We used CITE-seq to interrogate for correlation of cellular protein epitope and epitope-encoding transcript levels for transcripts/epitopes of particular interest in the context of ganglioglioma, including CD34, endothelial marker CD31, and the immune cell markers CD45RA, CD14, CD3, CD8A, CD4, and CD19. This also allowed correlation of transcriptomic profiles with the presence of these particular epitopes of interest.

We analyzed n=3725 cells from three tumors with CITE-seq (**sTable 1**). These findings reiterated much of the results from snRNA-seq, but due to the smaller number of cells used for CITE-seq, the latter assays appeared insensitive to some of the more rare phenomena observed in the snRNA-seq data (please see **Supplementary Information**, **sFigure 8,** and **sTable 6** for more information). CITE-seq analysis confirmed a strong correlation between *CD34* expression and expression of the CD34 epitope commonly used for diagnostic purposes (R=0.37, p<2.2e-16 by Pearson correlation test, **sFigure 8S).** These results confirm the snRNA-seq results using an orthogonal method which includes protein-level data (CITE-seq).

### Spatial transcriptomics identified neuroectoderm neural precursor-like cells deep in neoplastic niches

To determine spatial context of tumor cells, four gangliogliomas analyzed by snRNA-seq above were subjected to spatially-resolved transcriptomic profiling (stRNA-seq). H&E slides were manually neuropathologist (GYL)-annotated with exclusion of spots with significant artifact, folded tissue, tissue degradation, and/or blood products. StRNA-seq yielded a mean of n=1791 high-quality RNA-seq profiles per tumor in 55 µm spots (**sFigure 9A-D**; note that with this spot size, there are multiple cells per spot). Two of the four slides show appreciable histological variation. Tumor 1 had distinct (prominently or minimally/mildly) myxoid, hypercellular, hypocellular, and vascular areas making up 10 distinct annotations (**sFigure 9C**). Tumor 4 had (prominently) myxoid, non-myxoid, generic tumor, and vascular areas making up 4 distinct annotations (**sFigure 9D**). K-nearest neighbor clustering correlated with these distinctions (**sFigure 9E-F**). Final clusters were determined based on the combination of histopathological annotation and k-nearest neighbor clustering (**Figure 8A-B**).

**Figure 8:**
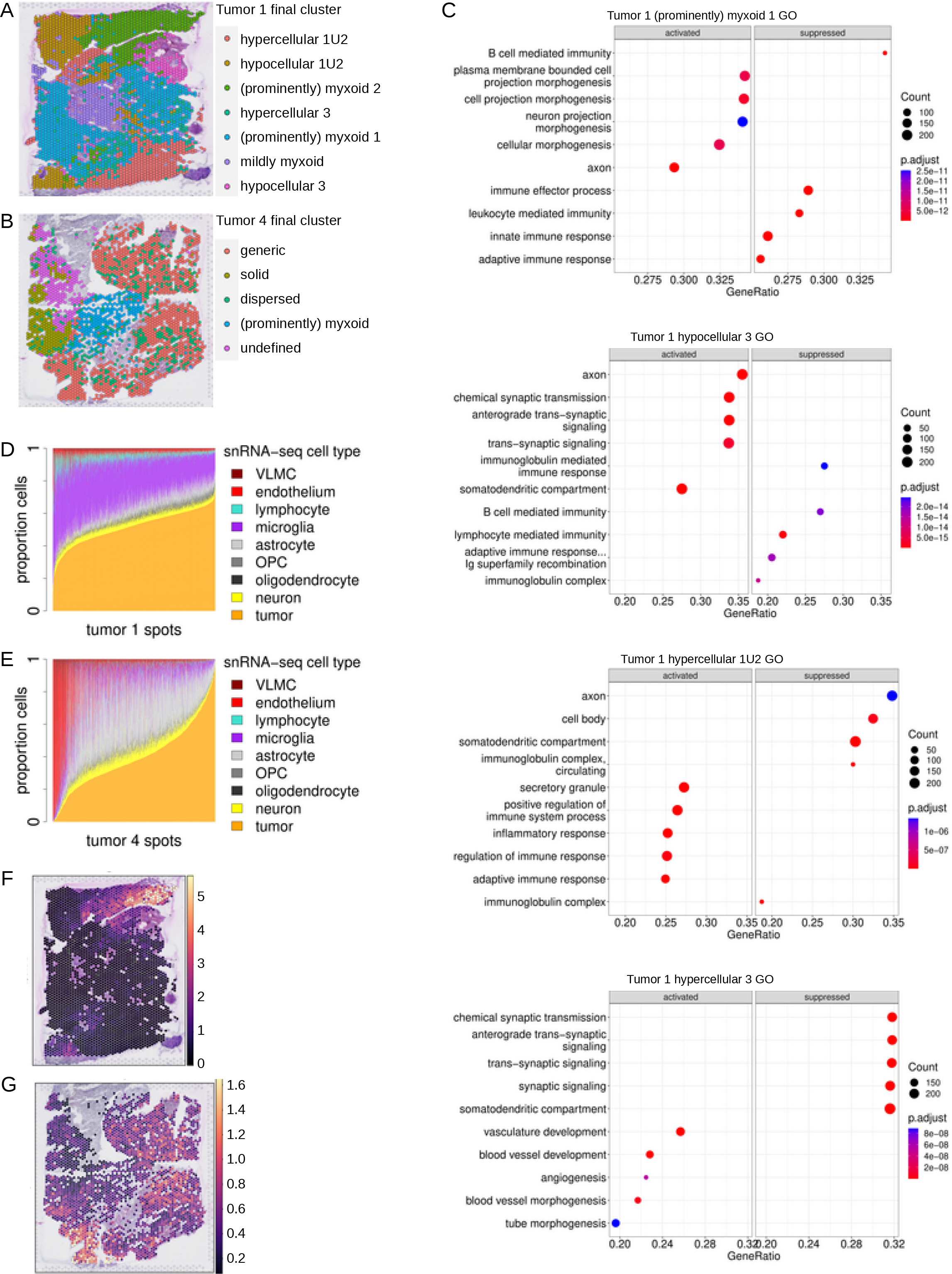
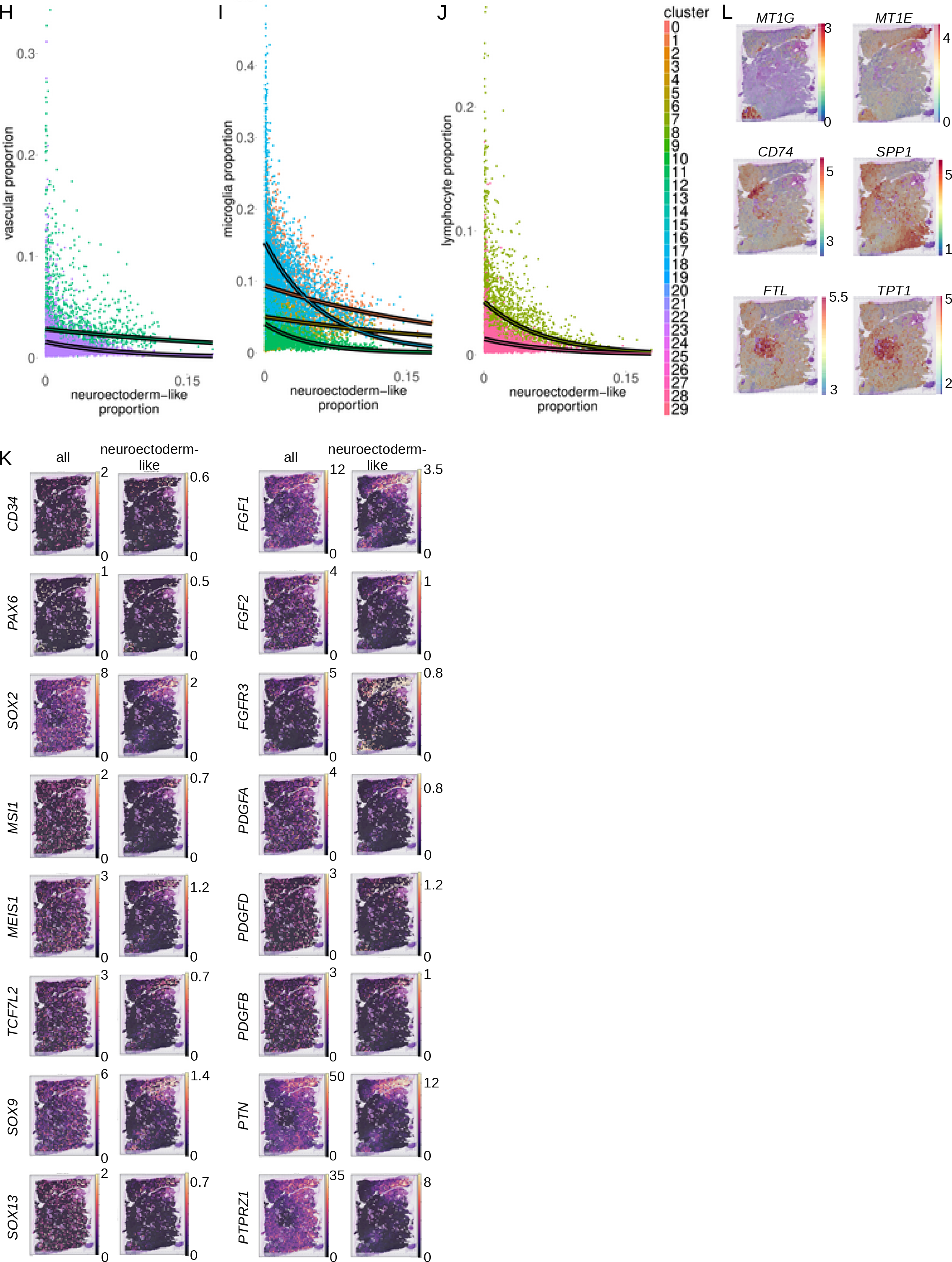
Ganglioglioma spatial transcriptomics shown for two samples from tumor 1 and tumor 4. (A-B) Spatial feature plots of final clusters for tumor 1 (A) and tumor 4 (B). Note ‘U’ in the final cluster name is used to denote union of multiple Space Ranger-generated graph-based (kNN) clusters. (C) GO term enrichment for select tumor 1 clusters. (D-E) Cell type proportions per spot as determined by cell2location with snRNA-seq as reference for tumor 1 (D) and tumor 4 (E). OPC=oligodendrocyte precursor cell. VLMC=vascular leptomeningeal cell. (F) Tumor 1 cell2location-calculated neuroectoderm neural precursor cell-like cell abundance per spot. (G) Analogous to (F) but for tumor 4. (H-J) Inverse correlation of neuroectoderm neural precursor-like cell proportion and that of vascular (H), microglial (I), and lymphoid (J) cells, shown for tumor 1 spot cell proportions calculated by cell2location with snRNA-seq clusters as reference. Points and curves are colored according to the cell type whose proportion is being compared to the proportion of neuroectoderm neural precursor-like cells. Curves (outlined in black to enhance visibility) represent non-linear (exponential) least-squares fit for each reference cell type. (K) Overall (left) and cell type-specific (right) expression of neuroectoderm neural precursor cell-like cell markers as well as important transcription factors and ligand-receptor pairs based on snRNA/CITE-seq analysis. (L) Feature plots of log1p (base 2) [RNA] for examples of top spatially-variable features from tumor 1, as determined by mark-variogram method in Seurat.

To characterize these distinct tumor regions, top cluster-specific differentially expressed genes were determined, and cluster-variable pathway activation was probed by clusterProfiler[85]. Several cancer-relevant patterns emerged (**Figure 8C, sFigure 9G,** and **sTable 7**). There was a clearly inverse correlation between enrichment of immune cell response/effector function pathways and enrichment of neuronal- (or macroglial-) function related pathways. For instance, in tumor 1, clusters (prominently) myxoid 1, (prominently) myxoid 2, and hypocellular 3 were each characterized by depletion of various immune response pathways but with enrichment of neuronal morphogenesis pathways ((prominently) myxoid 1), GO astrocyte projection ((prominently) myxoid 2), or neuron-related pathways (hypocellular 3, **Figure 8C**). In contrast, in tumor 1, clusters mildly myxoid, hypocellular 1U2, and hypercellular 1U2 each showed enrichment of numerous immune response pathways with associated depletion of cell morphogenesis (mildly myxoid), synaptic pathways (hypocellular 1U2), other neuron-related pathways (hypercellular 1U2), or neural crest differentiation (hypercellular 1U2, **Figure 8C**). Hence, tumor cellular composition appeared quite heterogenous on the molecular level with pro-inflammatory - and presumably anti-tumor - environments and neoplastic glioneuronal cell areas that appear to be relatively privileged from immune regulation. There also appeared to be an inverse correlation between enrichment of neuron-related pathways and vascular genesis-related pathways including the Wikipathways VEGFA-VEGFR2 pathway. For each of tumor 1 and tumor 4, there were dispersed spots that co-clustered (into tumor 1 hypercellular 3 and tumor 4 dispersed) wherein there was significant enrichment in vascular genesis-related pathways and Wikipathways VEGFA-VEGFR2 pathway with associated enrichment of inflammatory pathways, PI3K-AKT(-mTOR) pathways, and depletion of neuron-related pathways (**Figure 8C** and not shown). In contrast, tumor 1 hypocellular 3 showed depletion of Wikipathways VEGFA-VEGFR2 pathway in the context of significant enrichment of multiple neuron-related pathways (**Figure 8C** and not shown). Other cluster-specific differences also emerged. For example, tumor 1 (prominently) myxoid 1 enrichment of Wikipathways glioblastoma signaling and Wikipathways WNT signaling pathway and pluripotency; tumor 1 (prominently) myxoid 2 enrichment of GO zinc homeostasis and depletion of Wikipathways oxidative phosphorylation; tumor 1 hypocellular 1U2 depletion of various (m)RNA metabolic pathways (**Figure 8C** and not shown).

Given the multiple cells per spot, stRNA-seq spot cell composition was determined by decomposition/deconvolution by cell2location[39] and UCell signature scoring[3] using ganglioglioma snRNA-seq and CITE-seq data as parallel references. Cell composition appeared to vary considerably, yet appropriately, within slides and also between samples with spots estimated to be composed of typically 20-80% neoplastic cells (**Figure 8D-E**). Substantial proportions of each of microglia and macroglia were found as were smaller proportions of neurons, lymphocytes, and vascular cells, as expected. Cell2location tissue regions derived from cell abundance kNN generally correlated with histopathological annotations and up front kNN clustering (**sFigure 9H-I**), further supporting the model-calculated cell abundances. For slides from two of the tumors, methods concurred on the presence of clusters of significant areas of neuroectoderm neural precursor-like cell signature (**Figure 8F-G** and **sFigure 9J-K**).

In order to better understand important intercellular interactions, particularly with the neuroectodermal neural precursor-like cells, we tested for co-localization of different cell types to any given spot. When comparing all four tumor samples, themes emerged. Interestingly, neuroectoderm neural precursor cell-like cell abundance was highly inversely correlated with the presence of vascular cells, microglia, or lymphocytes (**Figure 8H-J**). This suggests these cells occupy a hypovascular, perhaps relatively hypoxic environment. Also, they appear to be relatively spared of immune targeting. Other than relative exclusion from these precursor-like cell areas, microglia resembling those from each of snRNA-seq cluster 1, 5, and 11 appeared randomly dispersed throughout each of the four tumor samples. Microglia resembling those from snRNA-seq cluster 17 appeared somewhat more concentrated in certain regions of some tumors. For instance, these cells were especially abundant in the Tumor 1 hypercellular 1U2 and hypocellular 1U2 clusters and in the Tumor 4 solid cluster. In contrast, neuroectoderm neural precursor cell-like cell abundance appeared strongly correlated with cell abundance of other neoplastic clusters (not shown).

To further investigate co-localization, stRNA-seq transcriptomic profiles were subjected to non-negative matrix factorization (NMF)[39]. Interestingly, tumor 4 neuroectoderm neural precursor -like cells tended to occupy discrete locations, with these cells separating out at a low number of factors compared to the number of reference clusters (by k=13 for snRNA-seq n=30 clusters; not shown). Of note, tumor 4 neuroectoderm neural precursor-like cells separated from endothelial cells very readily (the vast majority by k=5 with n=30 reference snRNA-seq clusters; not shown). Similar to what was seen for tumor 4 neuroectoderm neural precursor-like cells, tumor 1 neuroectoderm-like cells separated by k=5 from endothelial cells (**sFigure 9L-N**). These findings in tumor 1 and tumor 4 suggest endothelial cells are readily distinguished from and spatially distinct from neuroectoderm neural precursor-like cells. In contrast, at least a subset of tumor 1 neuroectoderm neural precursor-like cells appeared to co-localize with particular other neoplastic cell types (**sFigure 9L-N**). These observations appear consistent with these neuroectoderm neural precursor cell-like tumor cells residing deeply within neoplastic niches.

We next sought to determine the signaling and transcriptional program context of neuroectodermal neural precursor cell-like tumor cell niches. Interestingly, there was visually apparent co-localization of tumor neuroectoderm-like cells with high *PTPRZ1*, *PTN*, and *FGFR3* (**Figure 8K**), identified as important for neuroectoderm-like neoplastic cell signaling as above. Cell type-specific expression was also estimated by cell2location. This identified tumor neuroectoderm-like cell-specific expression of important neuroectoderm neural precursor like-cell markers including *CD34*, *PAX6*, *SOX2*, *MSI1*, *MEIS1*, and *TCF7L2*, further validating the nature of the snRNA-seq neuroectoderm-like cells and the identification of these cells *in situ* despite their being a small minority of tissue cells analyzed spatially (**Figure 8K**). Interestingly, there was significant neuroectoderm-like cell-specific *PTN*-*PTPRZ1*, *FGF1/2-FGFR3*, and *PDGFA/D-PDGFRB* expression (**Figure 8K**). This suggests a substantial autocrine mechanism for maintenance of the ganglioglioma stem/progenitor cell compartment.

To further explore the spatial context of ganglioglioma cells, top spatially variable genes, agnostic to cell type assignment, were uncovered using the mark-variogram method within Seurat[32]. Interesting patterns emerged. For example, for tumor 1, metallothioneins dominated top spatially-variable features, including *M1E* (#4), *MT1G* (#6), *MT3* (#7), *MT1X* (#10), *MT1M* (#13), and *MT2A* (#24) among the top 25 (**Figure 8L** and not shown). Interestingly, metallothionein expression appeared correlated with neuroectoderm neural precursor-like cell abundance, with high *MT1E*, *MT1G*, *MT1X*, and *MT1M* each exclusive to areas of neuroectoderm-like cells (**Figure 8L** and not shown). In contrast, *CD74* (antigen-presenting cell marker) and *SPP1* (oligodendrocyte and microglia marker[63, 92]) were top-25 spatially-variable transcripts that were largely excluded from neuroectoderm neural precursor-like cell areas (**Figure 8L**), once again consistent with the findings above suggesting neuroectoderm neural precursor-like cells to occupy niches with relatively little immune cell occupancy. *FTL* and *TPT1* had somewhat different distributions but were similarly inversely associated with neuroectoderm neural precursor-like cell abundance (**Figure 8L**).

### Neuroectoderm neural precursor-like programs associate with ganglioglioma bulk transcriptomic profiles

We hypothesized that particular ganglioglioma cell states could be detected in bulk transcriptomic data to varying degrees among different low grade glioneuronal tumors and gliomas. This, in turn, could assist with disease classification. To test this hypothesis, we obtained clinically-annotated bulk pediatric low grade glioma/glioneuronal tumor transcriptomic data sets (n=151 patients for Bergthold *et al*. of whom n=122 had clinical outcomes annotations and n=81 patients for the US National Cancer Institute’s Clinical Proteomic Tumor Analysis Consortium (CPTAC) with the diagnosis of ganglioglioma or pediatric low grade glioma)[10, 59].

To determine the degree of representation of the single nucleus/CITE-seq ganglioglioma cluster cells in the bulk data, deconvolution of the bulk data was performed using deconvolution/decomposition algorithms[60] and UCell signature scoring[3]. Despite differences in the nature of the bulk datasets (e.g. 6k gene array vs. bulk RNA-seq), the methods tested, and the cells present in reference snRNA-seq and CITE-seq datasets, common themes emerged. Bulk-sequenced tumors were identified as composed of ∼40-70% tumor cells with microglial, macroglial, and vascular minority populations identified (**Figure 9A-B**). In comparing cell composition by diagnosis, other low grade gliomas - pilocytic astrocytoma in particular - actually appeared to have greater components of the OPC-like cell states (**Figure 9C-D**). Conversely, gangliogliomas appeared to have a greater neuron component than other low grade gliomas and pilocytic astrocytoma in particular (**Figure 9C-D**). SnRNA-seq neuroectoderm neural precursor-like cell abundance appeared lowest among pilocytic astrocytomas and at least trended somewhat higher for other histologies, including ganglioglioma (where it was still rare, with median 2% of ganglioglioma cells from the Bergthold *et al*. dataset and 1% of ganglioglioma cells from the CPTAC dataset[10, 59], similar to expected, **Figure 9C-D**).

**Figure 9:**
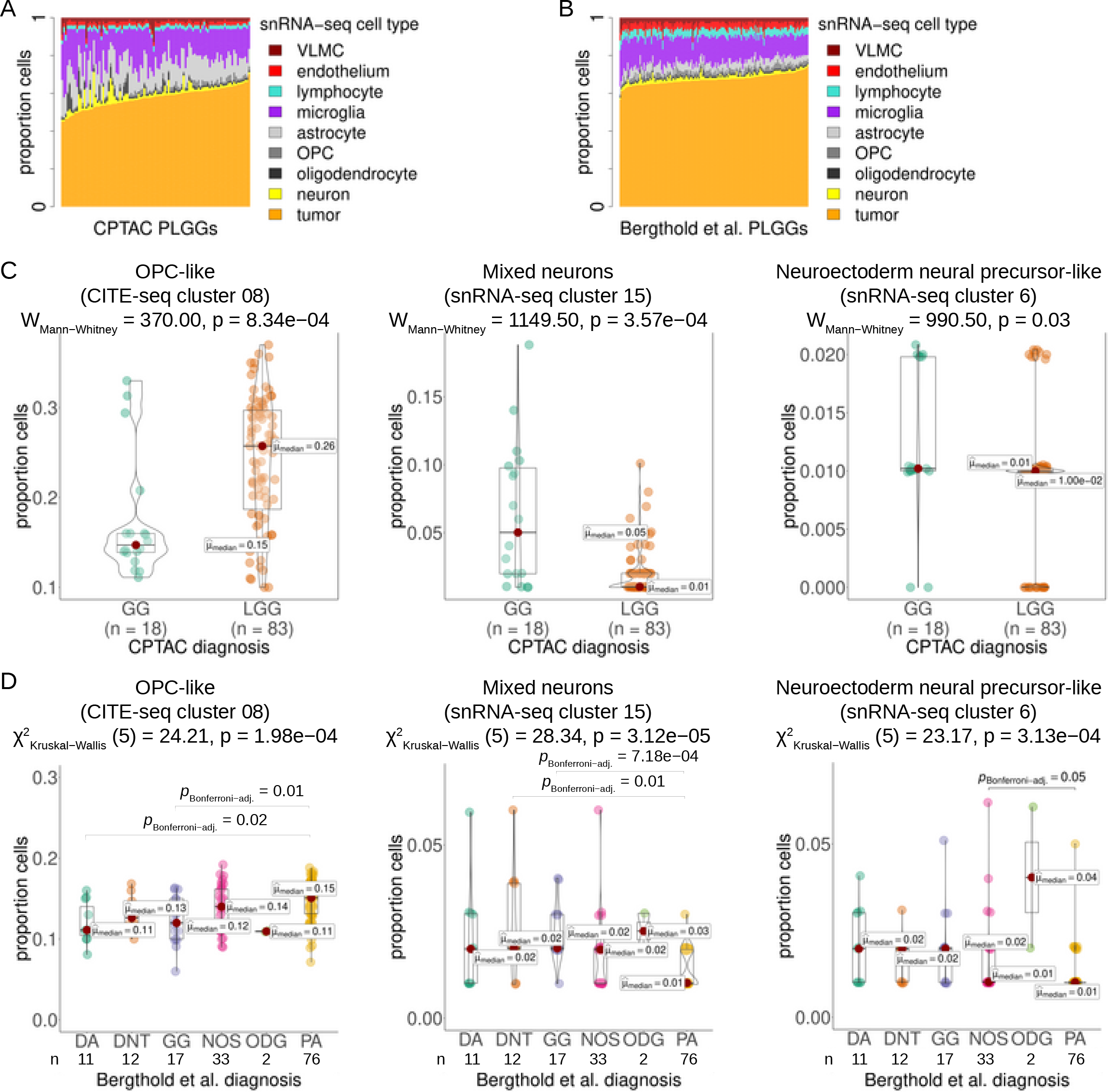
Pediatric low grade glioma bulk transcriptomic data deconvolution in light of snRNA/CITE-seq results. (A) CPTAC pediatric low grade glioma and ganglioglioma bulk RNA-seq deconvolution example using dtangle method with snRNA-seq data cell type reference. (B) Bergthold *et al*. low grade glioma bulk RNA-seq deconvolution analogous to (A). (C) Violin plots with CPTAC tumor OPC-like, mixed neuron, and neuroectoderm-like cell composition, by diagnosis. Medians marked with red dots and labeled. Statistical comparisons by Mann Whitney with p-values shown. (D) Analogous to (C) for Bergthold *et al*. tumors except for statistical testing, Kruskal−Wallis was used initially, followed by Dunn test with Bonferroni adjustment as needed. OPC=oligodendrocyte precursor cell, DA=diffuse astrocytoma, GG=ganglioglioma, ODG=oligodendroglioma, DNT=dysplastic neuroepithelial tumor, NOS or LGG=low-grade glioma, not otherwise specified. PA=pilocytic astrocytoma.

Hence, bulk transcriptomic deconvolution confirmed that ganglioglioma neuroectodermal neural precursor-like cell, neuron(-like) cell, and OPC(-like) cell abundance are potentially useful for disease classification.

### Prognostic genetic signatures

We hypothesized that neoplastic precursor cell markers and/or inflammatory markers derived from our deep transcriptomic approaches could inform development of prognostic signatures for ganglioglioma and similar tumors. We used the insights from the above deep transcriptomic analyses to nominate prognostic features for low grade glioma/glioneuronal tumor patients. For Bergthold *et al*. patients, clinical annotations included diagnosis, extent of resection, self-organizing matrix cluster, age, receipt of chemotherapy, BRAF status, event free survival (EFS), and death[10]. Resection status was found to be important with gross total resection associated with significantly better EFS whereas EFS was numerically similar to one another after biopsy, subtotal resection, and near total resection (**Figure 10A-B** and **sFigure 10A-B**). For CPTAC, clinical annotations included diagnosis (we excluded diagnoses other than ganglioglioma and other low grade glioma), extent of resection, BRAF status, tumor grade, EFS, and death[59]. Resection status was once again strongly associated with EFS (**sFigure 10C-H**).

**Figure 10:**
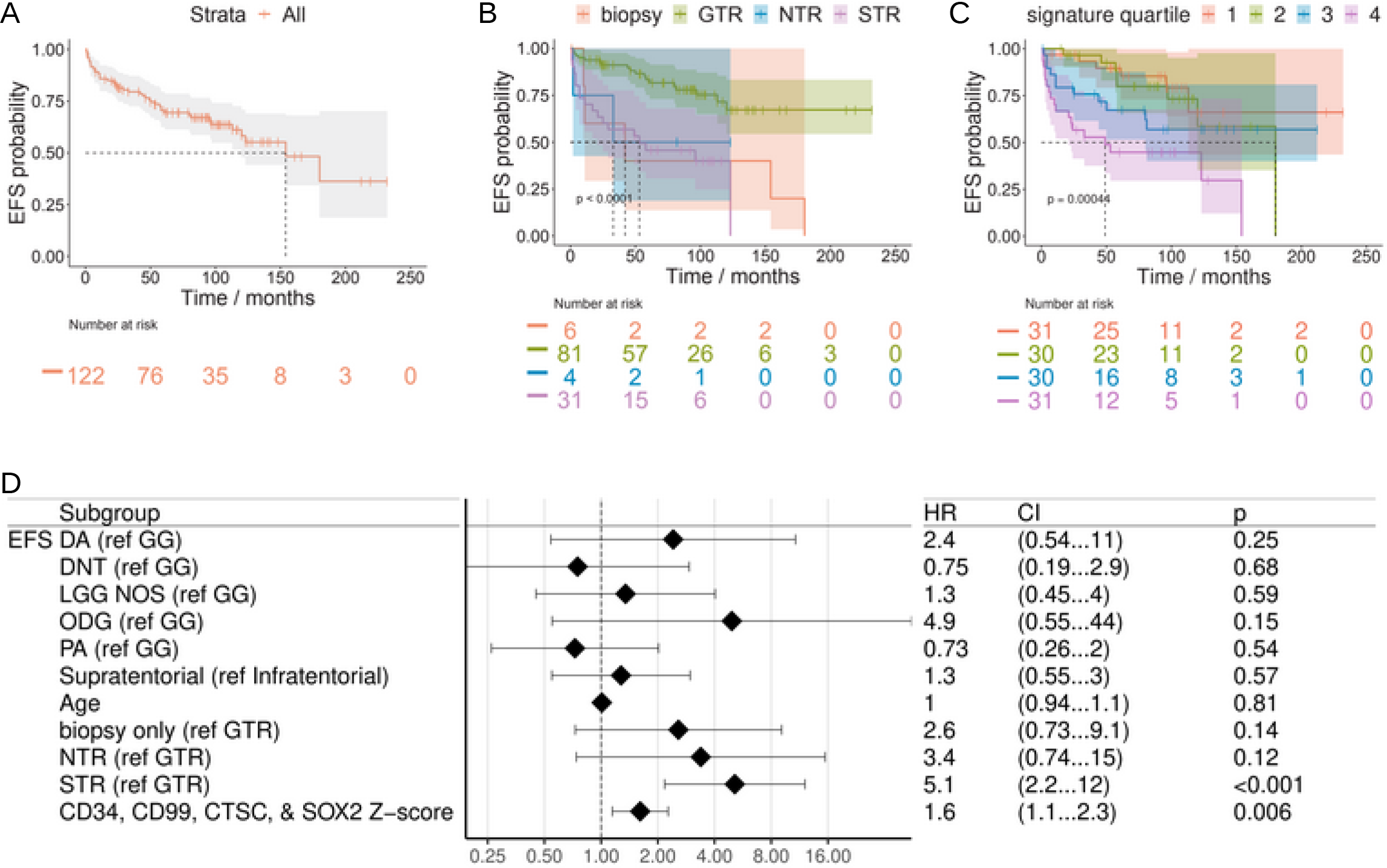
Low grade glioma outcomes analysis. (A-B) Kaplan-Meier EFS (95% CI shaded, dotted line for median) for Bergthold *et al*. patients overall (A) and by resection status (B). (C) Kaplan-Meier EFS (95% CI shaded, dotted line for median) for Bergthold *et al*. patients by UCell score quartile for the gene signature of CD34, SOX2, CD99, and CTSC. (D) Forest plot of COXPH EFS HR for Bergthold *et al*. patients with continuous variables kept continuous. Includes HR for Z-score for the gene signature of CD34, SOX2, CD99, and CTSC. EFS=event-free survival, NR=none reported, GTR=gross total resection, NTR=near total resection, G/NTR=gross or near total resection, STR=subtotal resection, WT=wild-type. DA=diffuse astrocytoma, GG=ganglioglioma, ODG=oligodendroglioma, DNT=dysplastic neuroepithelial tumor, NOS or LGG=low-grade glioma, not otherwise specified. PA=pilocytic astrocytoma.

We next turned our attention to prognostic genetic markers. *CDKN2A* (and adjacent *CDKN2B*) loss has been previously identified in a small minority of gangliogliomas as well as in other pediatric CNS tumors, such as pleomorphic xanthroastrocytoma (where *CDKN2A/B* loss is typical)[23, 58, 68]. The prognostic significance of *CDKN2A/B* loss in ganglioglioma is unclear though there may be some association with more adverse histopathological features and poorer prognosis[23, 58, 68]. In contrast, we found *CDKN2A* expression was among the top event-associated transcripts in both data sets (Bergthold *et al*. #16/6100 and CPTAC #23/14827 by average log2-fold-change). When limiting the diagnosis to ganglioglioma, *CDKN2A* was not prognostic (Bergthold *et al*. rank #3201 and CPTAC rank #2840). Instead the association of *CDKN2A* expression with EFS events appeared to be associated with other diagnoses, mostly pilocytic astrocytomas.

The remaining top low grade glioma event-associated transcripts in common between Bergthold *et al*. and CPTAC were overwhelmingly associated with inflammation and/or the extracellular matrix. Particularly interesting was a possible clinical event-associated enrichment for components of the cytosolic DNA sensing pathways (represented in KEGG and Wikipathways, **sFigure 10I-J**), including the cGAS/STING pathway, which has apparent and/or potential roles in tumorigenesis, prognosis, and therapeutic approach[33].

To interrogate for prognostic multi-gene-signatures, we started with gene signatures of curated stemness-associated factors from the snRNA-seq data and unsupervised machine learning. From this approach, multiple 3-5 gene, curated signatures were of borderline significance, though none were robustly significant by both Kaplan-Meier and Cox proportional hazards modeling using both gene signature z- and Mann Whitney U-test-scoring (arbitrarily using score quarterization for the Kaplan-Meier method). For example, the four-gene signature *CD34*, *SOX2*, *CD99*, and *CTSC* scores appeared at least trending towards association with worse EFS in this regard (**Figure 10C-D** and **sFigure 10K-L**).

To develop more sophisticated prognostic genetic signatures and models, we employed supervised machine learning. A model was trained on Bergthold *et al*. (in mlr3/mlr3proba: top 2.5% clinical or genetic features selected by surv.rfsrc (fast unified random forests for survival, regression, and classification) embedded method, surv.aorsf (accelerated oblique random survival forests) learner selected (**sFigure 10M**) with subsequent sequential feature reduction (**sFigure 10N**) and learner hyperparameterization yielding tuned model with Harrell’s concordance-index of 0.87) and validated on CPTAC, yielding a 14-feature signature (resection status combined with expression of 13 transcripts: *UBE2M*, *ELAVL1*, *FRY*, *IGFBP4*, *STX4*, *PSMC6*, *MTX2*, *NFE2L3*, *C1QB*, *ELANE*, *SMARCD2*, *ZNF117*, and *ABL1*). This model predicted CPTAC outcomes with Gonen and Heller’s concordance-index 0.89. Hence, we were able to nominate two promising multi-gene prognostic signatures: a curated 4-gene signature and a 13-gene signature (in combination with resection status) from supervised machine learning.

## Discussion

Deep transcriptomics has recently led to advances in our understanding of many cancers, including brain tumors[27, 55, 67]. However, such techniques have not yet been published for glioneuronal tumors, and application of spatial transcriptomics approaches to brain tumors is in its infancy. Here, we applied deep transcriptomic approaches including spatial transcriptomics to gain insight into important questions regarding glioneuronal tumor biology. Our data supports our hypothesis that glioneuronal tumor CD34+ cells represent neuroectoderm neural precursor-like tumor precursor cells. Moreover, we found evidence of perturbations to mTOR/AKT signaling via PDGF/PDGFR and/or FGF/FGFR agonism in combination with PTN antagonism of the otherwise antagonistic and highly expressed PTPRZ1. The combination of this AKT perturbation with MAPK activation (which was present via BRAF V600E in all tumors studied presently) has been found to be sufficient for glioneuronal tumorigenesis in animal models[17]. Our data suggests such a mechanism for ganglioglioma tumorigenesis with CD34+ glioneuronal tumor cells as the tumor stem cells. Additionally, we identified important transcriptional regulators for primitive tumor cells including TCF7L2/MEIS1-PAX6 and SOX transcriptional cascades. Interestingly, though the neuroectoderm neural precursor-like cells appeared to occupy a hypovascular niche, they (at least among ganglioglioma neoplastic cells) appeared to be especially high users of oxidative phosphorylation pathways. It is possible this is related to an adaptation, e.g. via upregulation of mitochondrial activity given that the master regulator of mitochondrial biogenesis PPARGC1A was nominated as a neuroectoderm neural precursor-like cell associated transcription factor in an unbiased fashion by SCENIC.

This study reiterated neoplastic and stromal cell states common to myriad brain tumors. These include neoplastic OPC-like cells identified in primary brain tumors as diverse as (adult) glioblastoma[20], diffuse midline glioma[27], pilocytic astrocytoma[67], and now in glioneuronal tumors. Interestingly, these cells have in the past been hypothesized to be tumor precursor cells for other primary brain tumor types. However, subsequent study has not always been consistent with this hypothesis[79]. Within our study, we saw this effect. By the CITE-seq data alone with 1.8k neoplastic cells, the most primitive neoplastic cluster was an OPC-like cluster. However, with increase in cell sampling by an order of magnitude (i.e. to what was obtained for snRNA-seq), we were able to identify a rarer, much more primitive neuroectoderm neural precursor-like population that also fits with the hypothesized tumor stem cell population. This data points to the limitations of undersampling in this context or - conversely - the significant insights possible with sufficient sampling. Given the alignment of our results with *a priori* data and hypotheses, our sampling may very well be adequate to confidently identify the most important characteristics of ganglioglioma stem cells.

Our major results appear generalizable. For instance, (the rare) neoplastic *CD34*+ cells were found in tumor 1, tumor 3, tumor 4, and tumor 5 by snRNA-seq and/or CITE-seq. Moreover, we were able to uncover nests of the neuroectoderm neural precursor-like cells in tumors 1 and 4 by stRNA-seq and transcriptionally-similar cell states more broadly among gangliogliomas by deconvolution of bulk transcriptomic data.

In addition to neuroectoderm neural precursor-like and OPC-like neoplastic cells, we were able to identify myriad stromal cell types in the context of ganglioglioma, including oligodendrocytes, OPCs, astrocytes, various inhibitory and excitatory neurons, endothelial cells, VLMCs, microglia, and lymphocytes and their associated cell states including significant interactions with the neoplastic cells. We characterized the immune cells extensively. We weighed in on the controversy regarding tumor-associated myeloid cell M1-M2 polarization[5, 43]. Our data supports the model of aberrant co-activation of these programs rather than polarization between them. Furthermore, we found evidence that this aberrancy may be directly related to T lymphocyte dysfunction and T lymphocyte stimulation of M1 and M2 characteristics simultaneously. Our data also appears to further support recent observations that tumor-associated immune cells appear to fall along a spectrum rather than as discrete states[5]. Additionally, we found that even low grade tumors with tumor-infiltrating lymphocytes may include potentially suppressible regulatory components (e.g. T regs) and potentially anti-tumor cytotoxic T lymphocytes. Altogether, this data suggests a possible role for immune therapies such as immune check point inhibitors. However, we did uncover CTL exhaustion and dysfunction as potential limiting factors to be overcome.

Another theme of general interest that we uncovered was neoplastic PTPRZ1 antagonism. We found this among diverse primary brain tumors tested including glioblastoma, pilocytic astrocytoma, and ganglioglioma. Interestingly, PTPRZ1 appeared in our SCENIC SOX regulons, and SOX transcription factors are often hyperactive in primary brain tumors[73]. PTPRZ1 overexpression on its own would be expected to abrogate AKT/mTOR signaling and cancer progression[86]. These tumors appear to evolve to overcome this antagonism by overexpression of PTN. This suggests antagonism of PTPRZ1 antagonism by PTN as a possible therapeutic approach for many primary brain tumor types. FGF/FGFR and/or PDGF/PDGFR antagonism may also be important in the context of ganglioglioma, either in combination with PTN-PTPRZ1 antagonism or on their own.

Finally, we translated these findings into existing clinically-annotated bulk transcriptomic data sets to identify different cell state compositions therein as well as expound upon potentially clinically useful genetic markers and multi-gene signatures.

## Conclusions

We applied deep transcriptomic approaches including spatial transcriptomics towards study of open questions regarding glioneuronal tumor biology. Our data supports our hypothesis that GNT CD34+ cells represent neuroectoderm neural precursor-like tumor precursor cells. We found evidence of neoplastic cell dual perturbations to BRAF/MEK and PI3K/AKT/mTOR pathways and identified ganglioglioma gene regulatory networks (resembling those present during neuroectoderm neural development) and associated immune cell states. We translated these findings into low grade glioneuronal and glial tumor bulk transcriptomic deconvolution which suggested insights into tumor classification. Translation to prognostic signature nomination yielded insights into prognostication.

## Supporting information

sT1 Patient and tumor characteristics.xlsx

sT2 Top DEG by snRNA-seq cluster by Wilcoxon.xlsx

sT3 SnRNA-seq CellChat interaction probabilities by pathway.xlsx

sT4 SnRNA-seq CellChat interaction prob by sig L-R.xlsx

sT5 SnRNA-seq CellChat interaction p-val by sig L-R.xlsx

sT6 Top DEG by CITE-seq cluster by Wilcoxon.xlsx

sT7 Top DEG by stRNA-seq cluster by Wilcoxon.xlsx

sT8 SnRNA-seq cluster MSigDB cell type signature FGSEA.xlsx

sT9 SnRNA-seq cell MSigDB cell type neural signature UCell.xlsx

sT10 CITE-seq cluster MSigDB cell type signature FGSEA.xlsx

sT11 CITE-seq cell MSigDB cell type neural signature UCell.xlsx

Supplementary information 1 Supplementary text.docx

## Declarations

### Ethics approval and consent to participate

Ganglioglioma tumor samples for deep transcriptomics were obtained per Duke University Health System Institutional Review Board (IRB) protocol Pro00072150. The methods were carried out in accordance with the approved guidelines, with written informed consent obtained from all subjects or their guardians where appropriate.

### Consent for publication

Not applicable

### Availability of data and material

Seurat objects of CITE-seq, snRNA-seq, and stRNA-seq count matrices and associated annotations are uploaded on Zenodo (https://zenodo.org/record/7677962; DOI: 10.5281/zenodo. 7677962). Additional original data is available upon request.

### Authors’ contributions

JAR performed snRNA-seq, CITE-seq, stRNA-seq, bulk transcriptomic, and prognostic genetic signature in silico work, analysis, and interpretation and drafted and revised the manuscript. MEG performed pre-CITE-seq optimization and participated in manuscript drafting. VJ ran CellRanger or SpaceRanger for the snRNA-seq, CITE-seq, or stRNA-seq data and participated in data analysis and manuscript drafting. VC participated in data interpretation and manuscript drafting. DMA participation in project conception, experimental design, data interpretation, and manuscript drafting. SGG participated in experimental design, data interpretation, and manuscript drafting. EMT participated in project conception, tissue procurement, experimental design, and manuscript drafting. GYL participated in project conception, experimental design, neurohistopathological expert analysis of H&E slides, manuscript drafting, and manuscript revision. ZJR oversaw project conception, experimental design, data analysis and interpretation, manuscript drafting, and manuscript revision.

## Acknowledgments

The authors would like to thank members of the Duke Brain Tumor Center Biorepository and Database (Diane Satterfield, Merrie Burnett, and Elizabeth Thomas) and the Duke Molecular Physiology Institute (Karen Abramson and Emily Hocke) for assistance with this project.

## Methods

### Reference data sets

Reference single-cell data was obtained via UCSC cell browser (Nowakowski *et al*. developing brain[52], Eze *et al*. developing brain[25], Darmanis *et al*. glioblastoma[20], Wang/Muller *et al*. glioblastoma[82], Allen human cortex[93], Allen human M1 cortex[93]), Broad Institute single cell portal (Tirosh *et al*. oligodendroglioma[78], Filbin *et al*. diffuse midline glioma[27], Reitman *et al*. pilocytic astrocytoma[67]), and GEO GSE104276 (Zhong *et al*. developing prefrontal cortex[91]), GSE168408 (Herring *et al*. developing prefrontal cortex[34]), GSE144136 (Nagy *et al*. adult prefrontal cortex[49]), GSE156728 (Zheng *et al*. T cells[90]), GSE154763 (Cheng *et al*. myeloid cells[18]), and GSE114724 (Azizi *et al*. immune cells[5]). Bulk transcriptomic data and associated annotations were obtained from CPTAC[59] via pedcbioportal.kidsfirstdrc.org and, for Bergthold *et al*., from GEO GSE60898[10].

### Patients

Ganglioglioma tumor samples for deep transcriptomics were obtained per Duke University Health System Institutional Review Board (IRB) protocol Pro00072150. The methods were carried out in accordance with the approved guidelines, with written informed consent obtained from all subjects or their guardians where appropriate. Patient characteristics are summarized in **supplementary table 1**.

### SnRNA-seq

Frozen banked tumor tissue was obtained from the Duke Brain Tumor Center Biorepository and Database. For quality assurance, RNA integrity number was checked with goal RIN>7. Nuclei were isolated. Briefly, 50 mg tissue was minced to ∼0.5 mm cubes, transferred to lysis solution (10 mM Tris-HCl pH 7.4 (Sigma), 5 mM NaCl (Sigma), 3 mM MgCl_2_ (VWR), 0.1% NP-40 substitute (Sigma), and 0.5% RNasin Plus (Promega, aq)), incubated on ice 5 minutes, and incubated with tituration 10-15 times every 30 seconds for 10 minutes on ice. Residual debris was removed by 70 mcm filter (VWR). Nuclei were centrifuged 300g x5 minutes at 4°C, rinsed x2 with resuspension buffer (1% BSA (ThermoFisher), 0.5% RNasin Plus, and 1X PBS pH 7.4 (Corning, aq)), resuspended in OptiPrep solution (25 mM KCl (ThermoFisher), 5 mM MgCl_2_ (ThermoFisher), 20mM Tris-HCl pH 7.8 (ThermoFisher), 50% OptiPrep Density Gradient Medium (Sigma), and 100 mM sucrose (Sigma, aq)), pelleted 10000g x10 minutes at 4°C, resuspended in resuspension buffer, and assessed for intact nuclei. Otherwise, snRNA-seq was performed using 3′v3 Single Cell technology according to the manufacturer’s protocol (10X Genomics, San Diego, CA).

### CITE-seq

Fresh tumor tissue was obtained for CITE-seq. Tumor cells were dissociated, washed, and resuspended. Antibodies for CITE-seq were anti-CD34 (clone 581), anti-CD31 (clone WM59), anti-CD45RA (clone HI100), anti-CD3 (clone UCHT1), anti-CD8A (clone RPAT8), anti-CD4 (clone RPAT4), anti-CD14 (clone 63D3), and anti-CD19 (clone HIB19) TotalSeq-A antibodies (BioLegend, San Diego, CA). Cells from one tumor were used to optimize preparation by adding variable antibody concentrations to 1 million cells in 50 mcl, washing with 200 mcl FACS buffer, resuspending in 200 mcl FACS buffer, and analyzing by flow cytometry. Otherwise, cells were used for CITE-seq using 3′v3 Single Cell Immune Profiling technology according to the manufacturer’s protocol (10X Genomics, San Diego, CA).

### Spatial transcriptomics

Frozen banked tumor tissue was obtained from the Duke Brain Tumor Center Biorepository and Database. StRNA-seq was performed using Spatial 3’ v1 technology according to the manufacturer’s protocol (10X Genomics, San Diego, CA).

### SnRNA-seq and CITE-seq data preprocessing and initial processing

Raw sequencing data was processed into unique molecular identifier count (UMI) matrices using CellRanger (v6.1.2 for snRNA-seq and v3.1.0 for CITE-seq, human genome build GRCh38, cellranger mkfastq -> cellranger count, 10X Genomics). Processing beyond this point was carried out in Ubuntu 20.04 using R 4.2.1 or python version 3.x (depending upon python packages in use). Within Seurat v4[32], snRNA-seq log-normalization, scaling, SCTransformation, PCA, and UMAP (based on PCA dims 1-50) was performed. For the snRNA-seq projection shown, SCTransform was run based on all cells, 10k variable features, regressing out based on percent mitochondrial RNA, and otherwise defaults. For CITE-seq, gene symbols were updated with HGNChelper v0.8.1[81], and this was followed by log-normalization, scaling, PCA, and UMAP (based on PCA dims 1-50). Neighbors and clusters were also found within Seurat. Gene symbols were updated by HGNChelper v0.8.1 for analyses sensitive to antiquated gene symbols. For each data set, data integration (batch=sample) was attempted using several algorithms with varied feature sets and settings (see **Supplementary Information 1** for more details). The benefits of integration (in terms of reducing potential technical artifacts, which appeared minimal to begin with) were consistently outweighed by the loss of meaningful biological variation (as assessed qualitatively and quantitatively by NMI, ARI, ASW (cell type), and isolated label scores). Consequently, unmanipulated data was used for downstream analyses.

### StRNA-seq data preprocessing and initial processing

Raw sequencing data was processed into unique molecular identifier count (UMI) matrices using SpaceRanger v1.3.0 (human genome build GRCh38, spaceranger mkfastq -> spaceranger count, 10X Genomics). The graph-based clusters shown were also calculated with spaceranger count. Slides were manually annotated (GYL). Empty spots were excluded as were those with excessive artifact (folded tissue and excessive blood product in particular). Graph-based clusters were then regrouped based on manual annotation for final clusters. Processing beyond this point was carried out in Ubuntu 20.04 using R 4.2.1 or python version 3.x (depending upon python packages in use). Gene symbols were updated by HGNChelper v0.8.1.

### Initial cell typing

SnRNA-seq and CITE-seq cells were initially typed based expression of *a priori* markers of interest, favoring unbiased atlases where possible, such as the Human Protein Atlas[63, 92] and HuBMAP[19, 94]. Markers are outlined in the results. Additionally, cells were typed using Seurat v4-base label transfer to published reference atlases listed under data above[32]. Cells were also typed by UCell v2.0.1 signature scoring[3].

### Inference of copy number alterations

Neoplastic-appearing cells from merged snRNA-seq or merged CITE-seq data were analyzed by inferCNV v1.12.0 in parallel[55]. Oligodendroglioma and associated oligodendrocytes were used as positive and negative controls, respectively[78]. Tumor 3 snRNA-seq data was excluded due to low complexity. SnRNA-seq and CITE-seq stromal cells were also used as controls. For run, cutoff=0.1 was used. Data was subjected to apply_median_filtering with window_size=7. Runs were otherwise by defaults. Data was displayed using ComplexHeatmap v2.13.2. Results were verified with CopyKAT v1.1.0[28].

### Cellular hierarchy

From CellRanger outputs, counts were obtained by velocyto v0.17.17 run10x and otherwise using the defaults[46]. Data from different tumors was then merged and then subset by cell lineage (e.g. neoplastic, microglia, lymphoid, vascular) based on cell typing for RNA velocity analysis by scVelo v0.2.4[9]. The results shown were obtained using the package defaults, with modes described in the text. SCENT v1.0.3 signaling entropy rate results shown were calculated using the original counts and defaults with the net17Jan16.m used as the protein-protein interaction network adjacency matrix[77]. CellRank v1.5.1 pseudotimes were calculated using the CytoTRACE kernel[31, 42] based on the original counts after merging samples and subsetting by lineage (neoplastic, myeloid, and lymphoid). Tumor 3 snRNA-seq data was excluded due to low complexity resulting in aberrantly high calculated pseudotime. Defaults used unless specified otherwise.

### Ligand-receptor analysis

Significant snRNA-seq and CITE-seq ligand-receptor interactions were nominated in an unbiased manner among merged snRNA-seq or merged CITE-seq data using CellChat v1.5.0 and CellPhoneDB v3.1.0[24, 35]. Defaults were used for the results shown.

### Gene set enrichment analysis (GSEA)

Average log2-fold-change was calculated within Seurat v4 based on the comparisons described in the text[32]. Gene lists were sorted by descending log2-fold-change and clusterProfiler v4.4.4 gseGO run with p-value cutoff=0.05 with Bonferroni correction[85]. GseKEGG and gseWP were performed analogously. Defaults were otherwise used. KEGG pathways of interest were visualized by Pathview v1.36.1[45].

### Gene regulatory network inference

Transcription factors were nominated among snRNA-seq and CITE-seq using pySCENIC v0.12.0 defaults[69]. Transcription factors/regulons of interest were further curated by Wilcoxon rank sum test comparison using AUC log2-fold-change≥0.1. For individual clusters this was performed relative to all other clusters. For each neoplastic cluster, it was also performed relative to the union of stromal neural clusters. For neoplastic *CD34*+ cells, it was performed relative to neoplastic *CD34*-cells.

### StRNA-seq data deconvolution

StRNA-seq spots were deconvoluted using cell2location v0.1[39]. For deconvolution, mitochondrial, ribosomal, X, and Y transcripts were excluded. Each of snRNA-seq and CITE-seq data sets was used as reference in turn. N cells per spot was 12 and detection_alpha=20. Training max epochs=1000 for references and =30000 for spatial data. Deconvolution, NMF, and cell type-specific gene expression were otherwise calculated using recommended/default settings.

### Top spatially variable genes

Top spatially variable genes were calculated for each stRNA-seq sample in Seurat v4 using the mark-variogram method and defaults otherwise[32].

### Bulk transcriptomic data deconvolution

Bulk data deconvolution was performed with granulator v1.1.0 using defaults[60]. Dtangle results shown are the results from using defaults within granulator, subsequently scaled to total composition of one for each sample. Statistical analysis comparing subpopulations was performed using the ggstatsplot v0.9.4.900 package and otherwise as described in the text[56].

### Clinical data analysis

Bergthold *et al*.[10] and CPTAC[59] bulk transcriptomic data was filtered for those with annotation of outcomes (event free survival in particular). Kaplan-Meier curves were calculated survfit from survival package v3.4.0[76] and plotted with log-rank p-values using ggsurvplot from the survminer library v0.4.9[2]. Correlograms of available clinical annotations were created by calculation of bias-corrected Cramer’s V for nominal vs. nominal variables, Spearman for numeric vs. numeric variables, and ANOVA for nominal vs. numeric variables. Multi-gene signature z-scores were calculated by multiplying n gene z-scores and taking the nth root. Multi-gene Mann Whitney U-test scores were calculated using UCell[3]. Prognostic models were developed by supervised machine learning in mlr3 using mlr3proba[41, 71]. Bergthold *et al*. relative-count expression combined with clinical annotations was subjected to importance scoring using surv.rfsrc learner, the top 2.5% features were selected, and surv.aorsf learner selected with subsequent sequential feature reduction and learner algorithm hyperparameterization yielding tuned model using a 14-feature signature (resection status combined with 13 transcripts: *UBE2M*, *ELAVL1*, *FRY*, *IGFBP4*, *STX4*, *PSMC6*, *MTX2*, *NFE2L3*, *C1QB*, *ELANE*, *SMARCD2*, *ZNF117*, and *ABL1*). Surv.aorsf final parameters were n_tree=10000, n_split=5, n_retry=24, mtry_ratio=0.8111, control_type=cph, and split_min_stat=11.2. The model was validated on CPTAC FPKM expression combined with clinical annotations.

