## supplemental figures for "Ganglioglioma deep transcriptomics reveals primitive neuroectoderm neural precursor-like population"

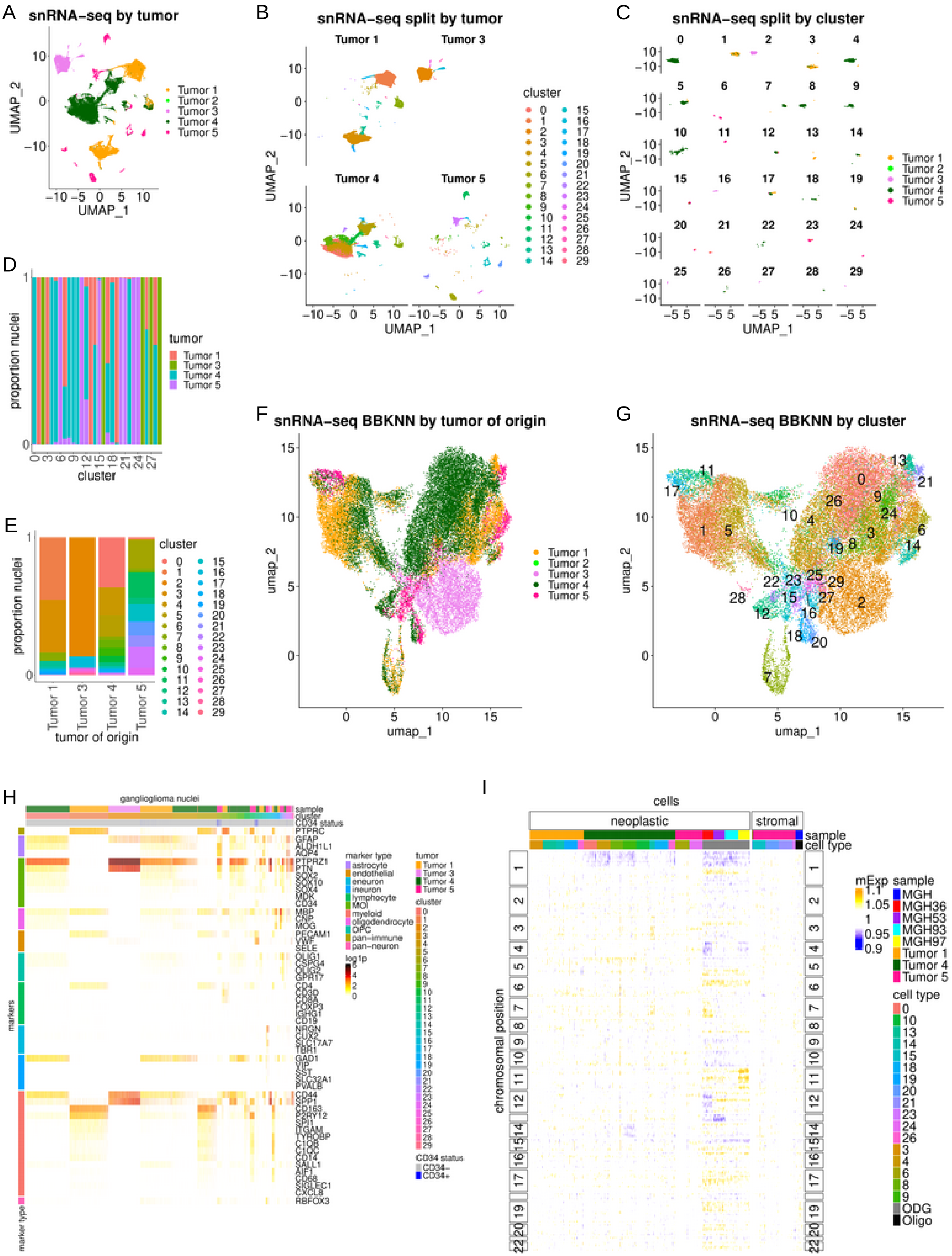
A
snRNA-seq by tumor
UMAP_1
UMAP_2
Tumor 1
Tumor 2
Tumor 3
Tumor 4
Tumor 5
B
snRNA-seq split by tumor
Tumor 1
Tumor 3
Tumor 4
Tumor 5
cluster
C
snRNA-seq split by cluster
D
proportion nuclei
cluster
tumor
E
tumor of origin
F
snRNA-seq BBKNN by tumor of origin
umap_1
umap_2
G
snRNA-seq BBKNN by cluster
H
ganglioglioma nuclei
sample
cluster
CD34 status
markers
marker type
astrocyte
endothelial
eneuron
ineuron
lymphocyte
MOI
myeloid
oligodendrocyte
OPC
pan−immune
pan−neuron
log1p
CD34−
CD34+
PTPRC
GFAP
ALDH1L1
AQP4
PTPRZ1
PTN
SOX2
SOX10
SOX4
MDK
CD34
MBP
CNP
MOG
PECAM1
VWF
SELE
OLIG1
CSPG4
OLIG2
GPR17
CD4
CD3D
CD8A
FOXP3
IGHG1
CD19
NRGN
CUX2
SLC17A7
TBR1
GAD1
VIP
SST
SLC32A1
PVALB
CD44
SPP1
CD163
P2RY12
SPI1
ITGAM
TYROBP
C1QB
C1QC
CD14
SALL1
AIF1
CD68
SIGLEC1
CXCL8
RBFOX3
I
cells
neoplastic
stromal
sample
cell type
chromosomal position
mExp
MGH
MGH36
MGH53
MGH93
MGH97
ODG
Oligo

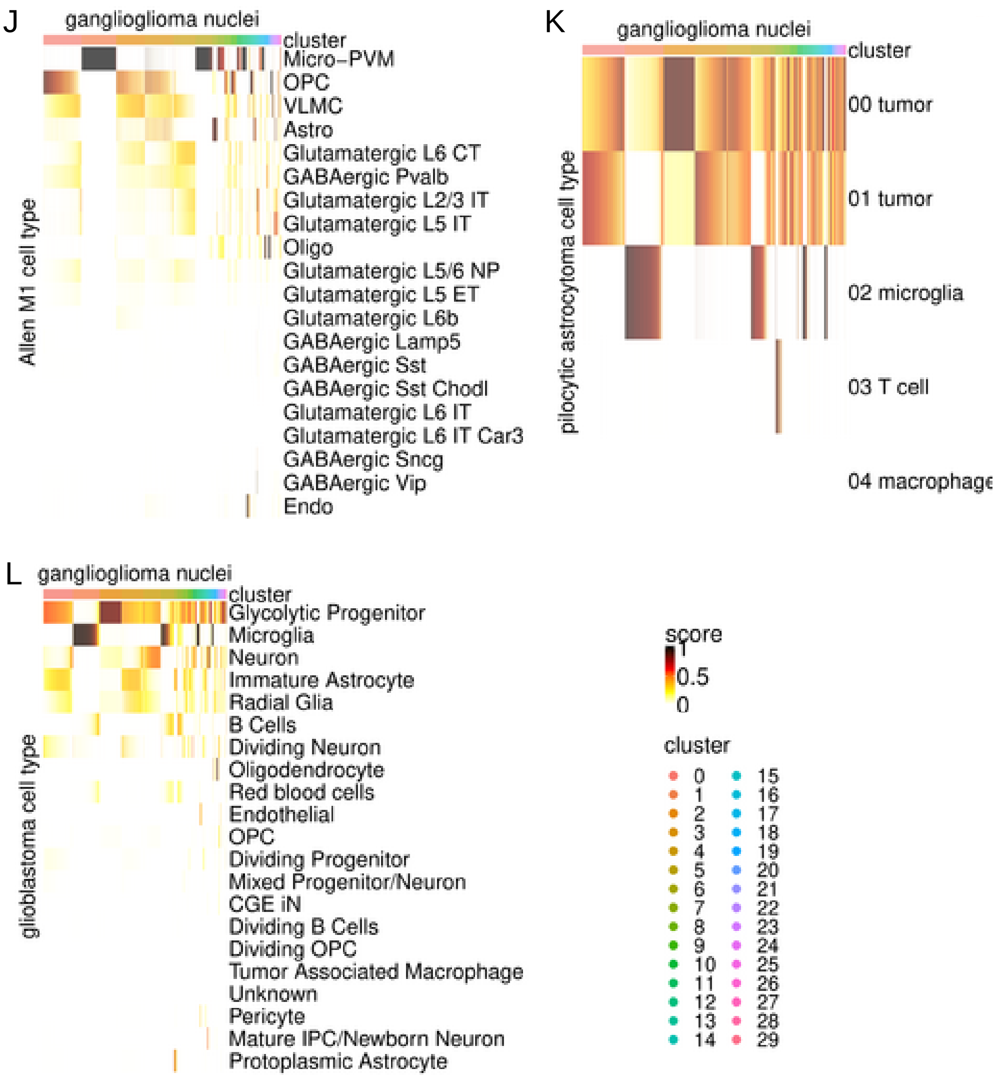

Supplementary Figure 1: Ganglioglioma single nucleus RNA-seq.
(A) UMAP plot of SCTransformed data from 34,907 nuclei, colored by tumor.
(B-C) UMAP plots split by tumor and colored by cluster (B) or split by cluster and colored by tumor-of-origin (C).
(D-E) Proportions of clusters by tumor (D) or tumors by cluster (E).
(F-G) Example of integrative assay UMAP output from BBKNN using highly variable genes and colored by tumor (F) or cluster (G).
(H) Heatmap of log1p (base 2) expression for cell typing markers of interest.
(I) Ganglioglioma inferred copy number variation based on snRNA-seq using inferCNV. Controls include oligodendroglioma samples (defined by 1p,19q codeletion; MGH36, MGH53, MGH93, and MGH97) Other known alterations include MGH36 del4,amp7,del12; MGH53 del13; and MGH97 del4,amp11. Findings were consistent with tumor 4 neoplastic-appearing cells with del1p with a neoplastic subpopulation with del14.
(J-L) Heatmap of Seurat label transfer results for ganglioglioma nuclei queried against adult brain (Allen M1, J), pilocytic astrocytoma (Reitman *et al*., K), and glioblastoma (Wang *et al*., L) reference atlases with query nuclei as columns and reference annotations as rows.

A

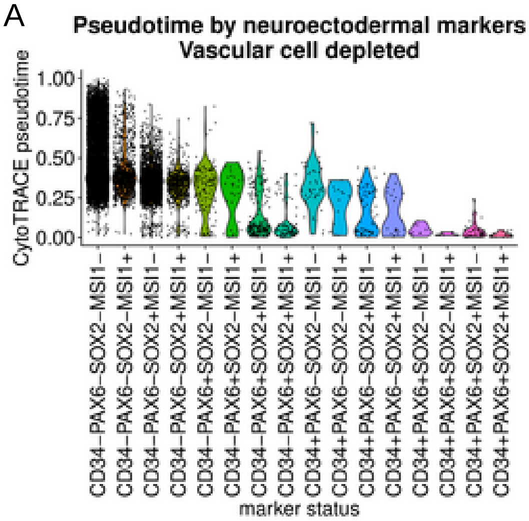

Supplementary Figure 2: Ganglioglioma potent neuroectoderm neural precursor-like cells.
(A) Violin plot of ganglioglioma snRNA-seq neoplastic cell CytoTRACE pseudotime by neuroectodermal neural precursor-like cell marker status.

A

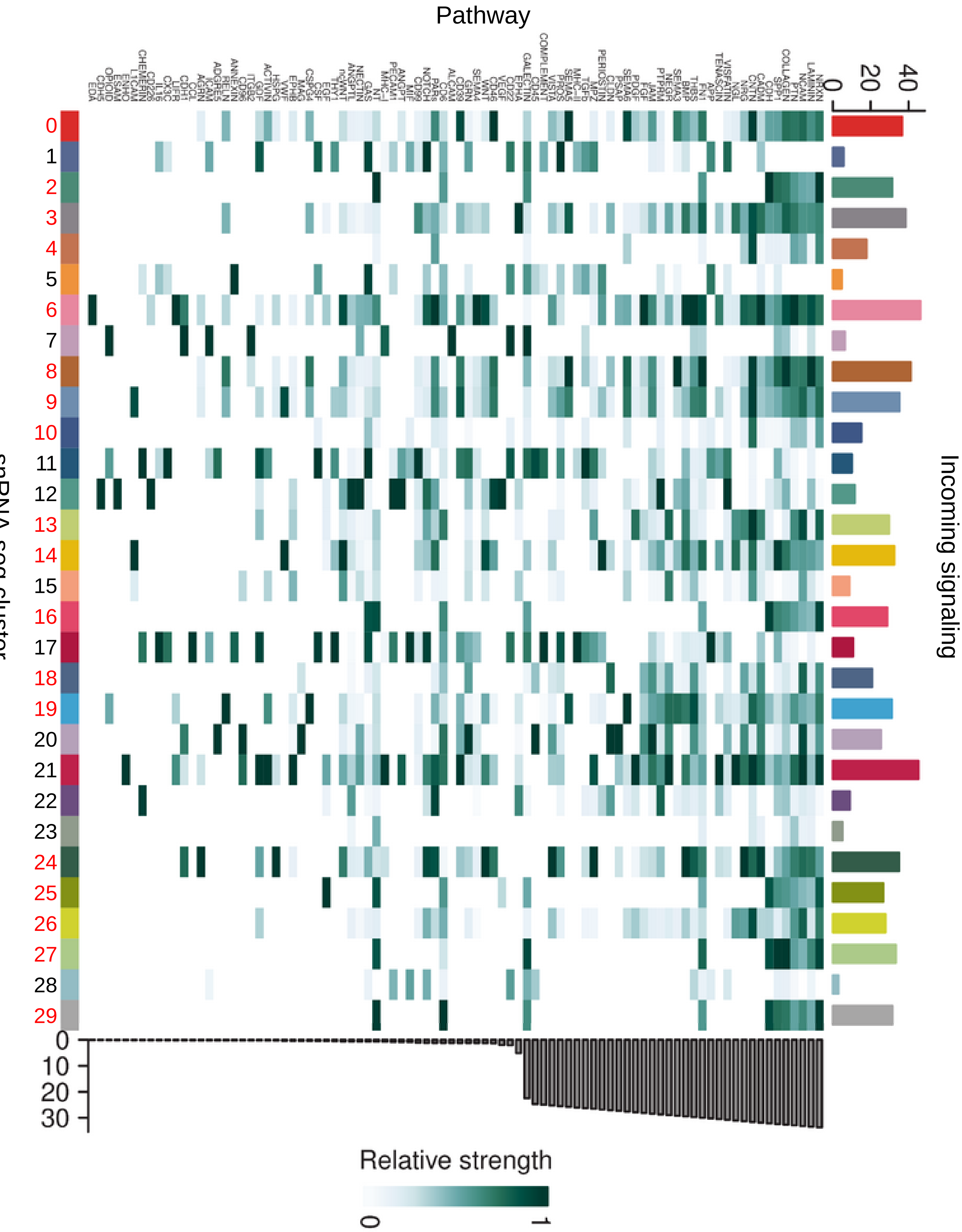
Pathway
snRNA-seq cluster
Incoming signaling
0
20
40
0
10
20
30
Relative strength
0
1

B

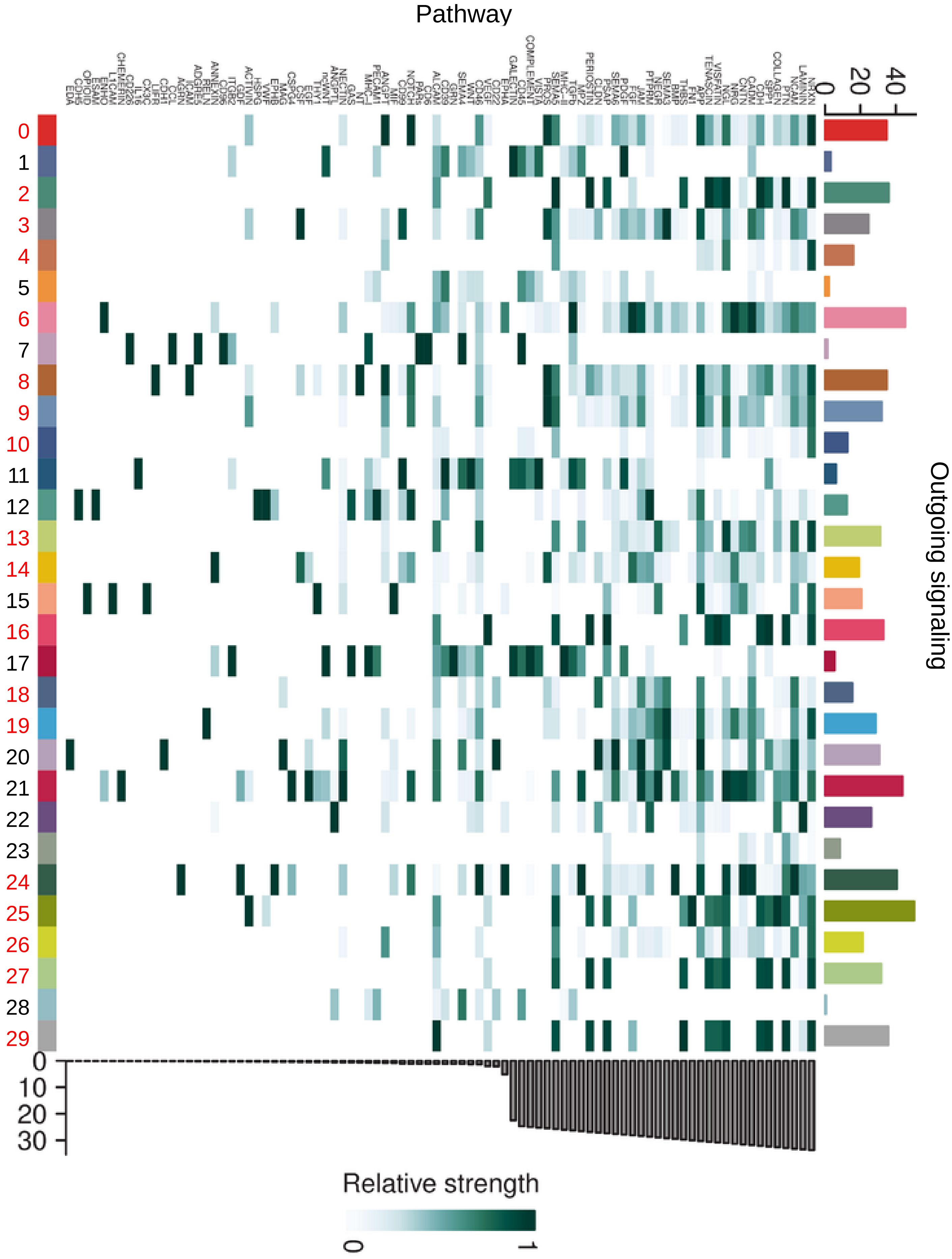
Pathway
snRNA-seq cluster
Outgoing signaling
0 20 40
0
10
20
30
Relative strength
0
1
0 1 2 3 4 5 6 7 8 9 10 11 12 13 14 15 16 17 18 19 20 21 22 23 24 25 26 27 28 29

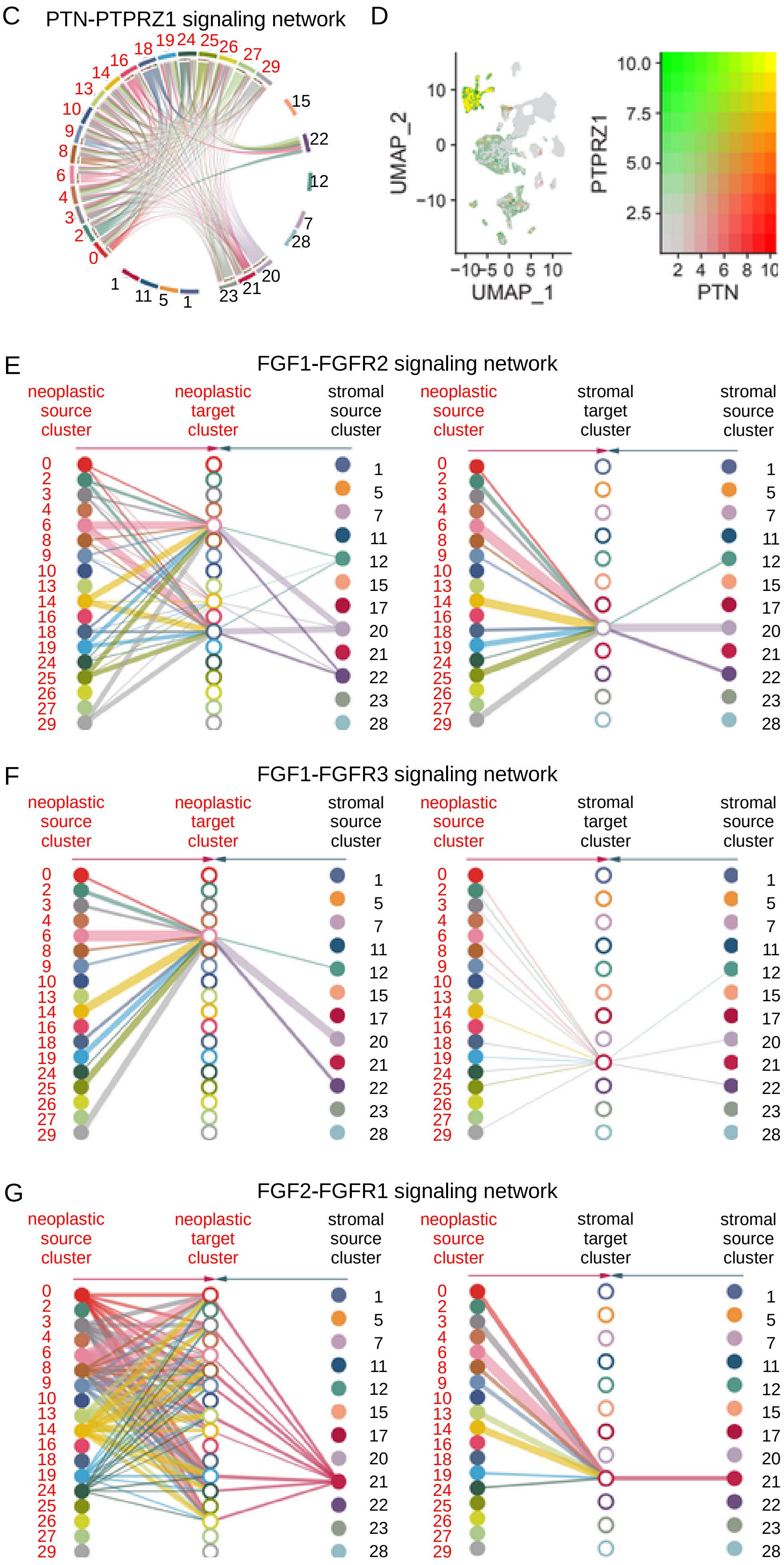
C
PTN-PTPRZ1 signaling network
0 2 3 4 6 8 9 10 13 14 16 18 19 24 25 26 27 29
15 22 12 7 28 20 21 23 1 5 11 1
D
UMAP_2
UMAP_1
10 0 −10
−10 −5 0 5 10
PTPRZ1
PTN
10.0 7.5 5.0 2.5
2 4 6 8 10
E
FGF1-FGFR2 signaling network
neoplastic source cluster
neoplastic target cluster
stromal source cluster
neoplastic source cluster
stromal target cluster
stromal source cluster
0 2 3 4 6 8 9 10 13 14 16 18 19 24 25 26 27 29
1 5 7 11 12 15 17 20 21 22 23 28
F
FGF1-FGFR3 signaling network
neoplastic source cluster
neoplastic target cluster
stromal source cluster
neoplastic source cluster
stromal target cluster
stromal source cluster
0 2 3 4 6 8 9 10 13 14 16 18 19 24 25 26 27 29
1 5 7 11 12 15 17 20 21 22 23 28
G
FGF2-FGFR1 signaling network
neoplastic source cluster
neoplastic target cluster
stromal source cluster
neoplastic source cluster
stromal target cluster
stromal source cluster
0 2 3 4 6 8 9 10 13 14 16 18 19 24 25 26 27 29
1 5 7 11 12 15 17 20 21 22 23 28

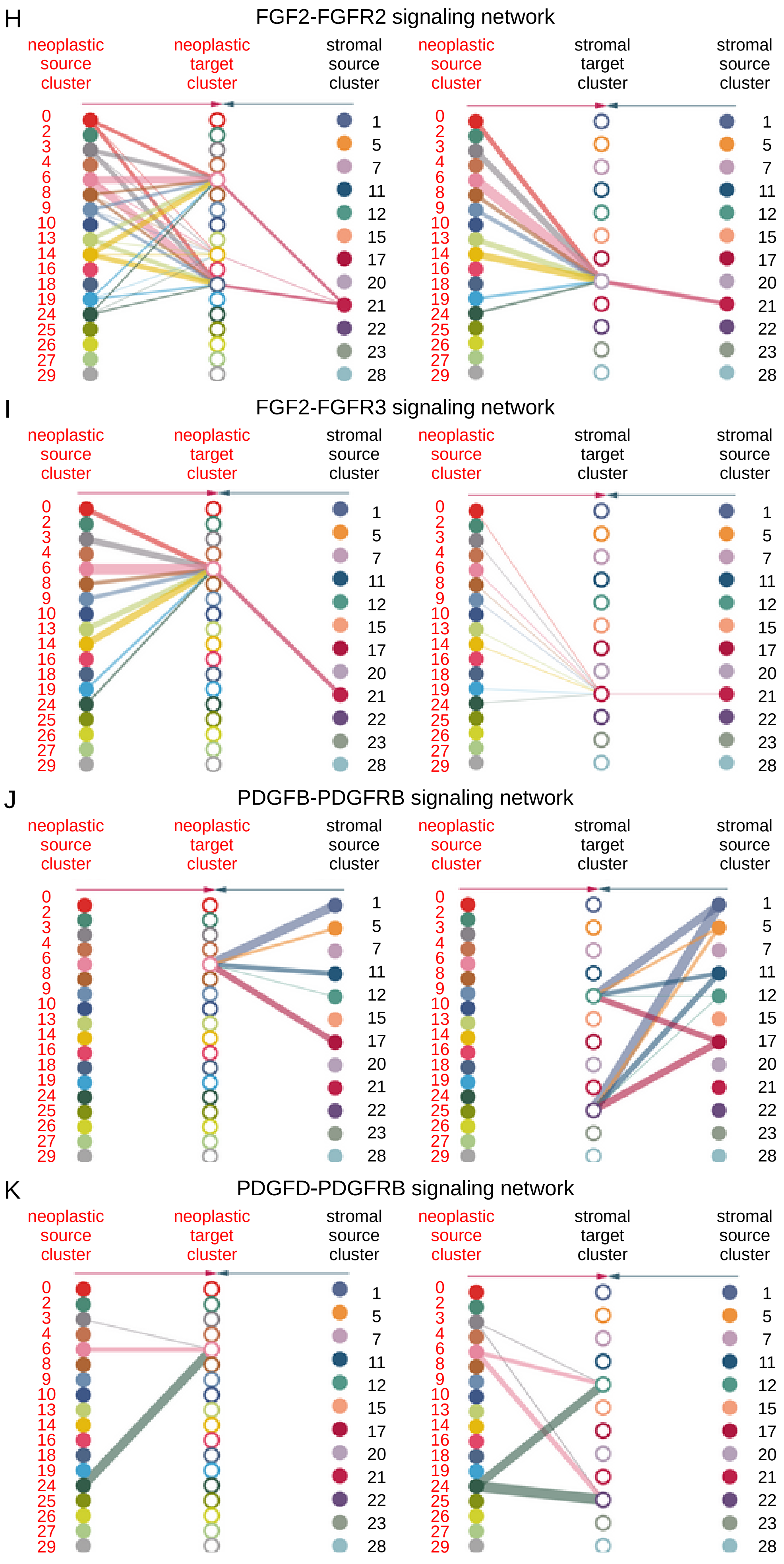
H
FGF2-FGFR2 signaling network
neoplastic source cluster
neoplastic target cluster
stromal source cluster
neoplastic source cluster
stromal target cluster
stromal source cluster
I
FGF2-FGFR3 signaling network
neoplastic source cluster
neoplastic target cluster
stromal source cluster
neoplastic source cluster
stromal target cluster
stromal source cluster
J
PDGFB-PDGFRB signaling network
neoplastic source cluster
neoplastic target cluster
stromal source cluster
neoplastic source cluster
stromal target cluster
stromal source cluster
K
PDGFD-PDGFRB signaling network
neoplastic source cluster
neoplastic target cluster
stromal source cluster
neoplastic source cluster
stromal target cluster
stromal source cluster
0 2 3 4 6 8 9 10 13 14 16 18 19 24 25 26 27 29
1 5 7 11 12 15 17 20 21 22 23 28

L

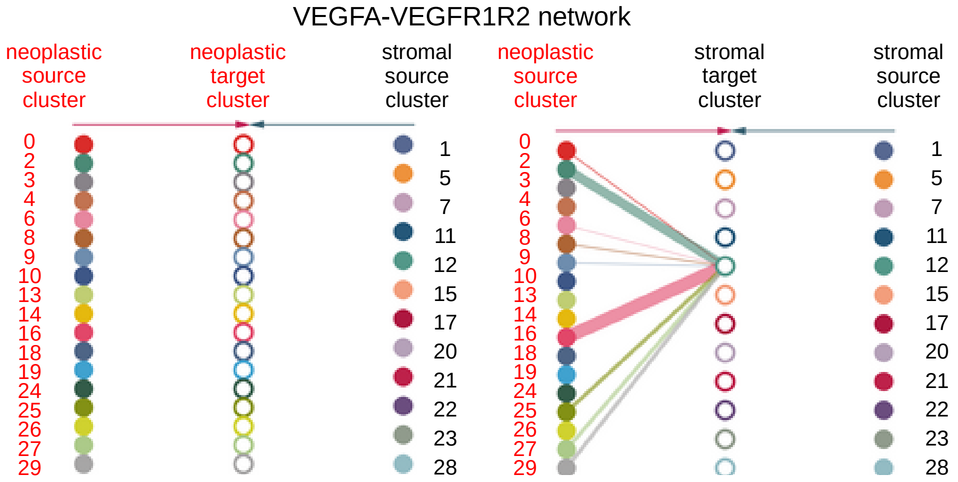

Supplementary Figure 3: Ganglioglioma neoplastic cell communications of interest by snRNA-seq evaluated with CellChat.
(A-B) Heatmap of significant incoming (A) and outgoing (B) signal ligand-receptor pathways by cluster. Neoplastic clusters are labeled in red. Stromal clusters are labeled in black.
(C) Significant PTN-PTPRZ1 interaction as nominated by CellChat with results shown as hierarchy plot cord diagram.
(D) Feature plot of ganglioglioma *PTN* (red), *PTPRZ1* (green), or both (yellow) expression.
(E) Significant FGF1-FGFR2 interaction as nominated by CellChat with results shown as hierarchy plots. Line thickness reflects strength of observed interaction. Interactions targeting neoplastic cells on the left plot and interactions targeting stromal cells on the right plot. Note that within each plot, neoplastic cluster sources are on the left and stromal cluster sources are on the right. Neoplastic targets are in the same order as neoplastic sources in the left plot, and stromal targets are in the same order as stromal sources in the right plot.
(F-L) Analogous to (E) for other top (statistically significant) ligand-receptor interactions.

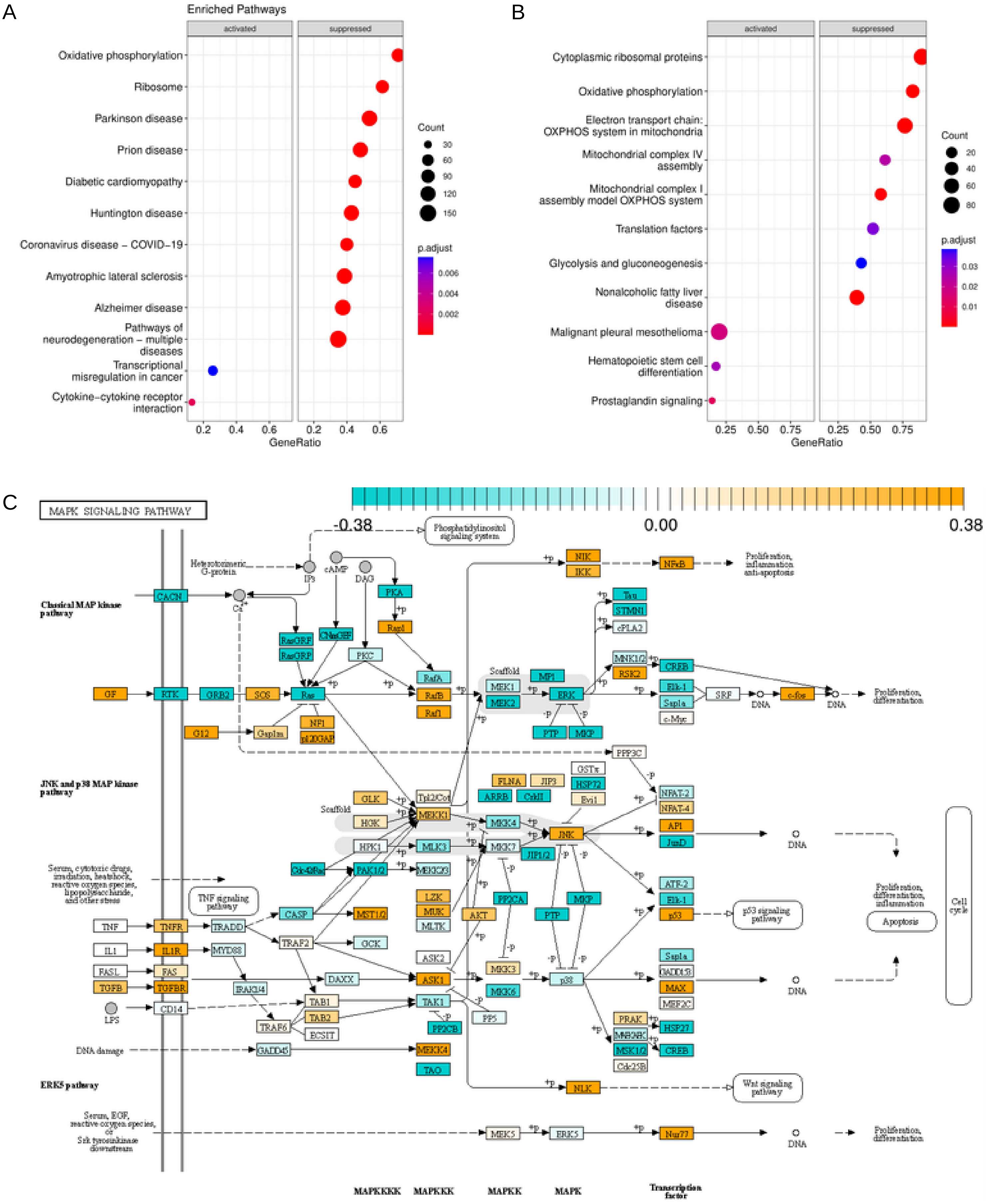
A
Enriched Pathways
activated
suppressed
Oxidative phosphorylation
Ribosome
Parkinson disease
Prion disease
Diabetic cardiomyopathy
Huntington disease
Coronavirus disease − COVID−19
Amyotrophic lateral sclerosis
Alzheimer disease
Pathways of neurodegeneration − multiple diseases
Transcriptional misregulation in cancer
Cytokine−cytokine receptor interaction
Count
30
60
90
120
150
p.adjust
0.006
0.004
0.002
0.2 0.4 0.6
GeneRatio
B
activated
suppressed
Cytoplasmic ribosomal proteins
Oxidative phosphorylation
Electron transport chain: OXPHOS system in mitochondria
Mitochondrial complex IV assembly
Mitochondrial complex I assembly model OXPHOS system
Translation factors
Glycolysis and gluconeogenesis
Nonalcoholic fatty liver disease
Malignant pleural mesothelioma
Hematopoietic stem cell differentiation
Prostaglandin signaling
Count
20
40
60
80
p.adjust
0.03
0.02
0.01
0.25 0.50 0.75
GeneRatio
C
MAPK SIGNALING PATHWAY
-0.38
0.00
0.38
Classical MAP kinase pathway
JNK and p38 MAP kinase pathway
ERK5 pathway
Phosphatidylinositol signaling system
TNF signaling pathway
p53 signaling pathway
Wnt signaling pathway
Proliferation, inflammation, anti-apoptosis
Proliferation, differentiation
Proliferation, differentiation, inflammation
Apoptosis
Cell cycle
DNA damage
DNA
MAPKKKK
MAPKKK
MAPKK
MAPK
Transcription factor

D

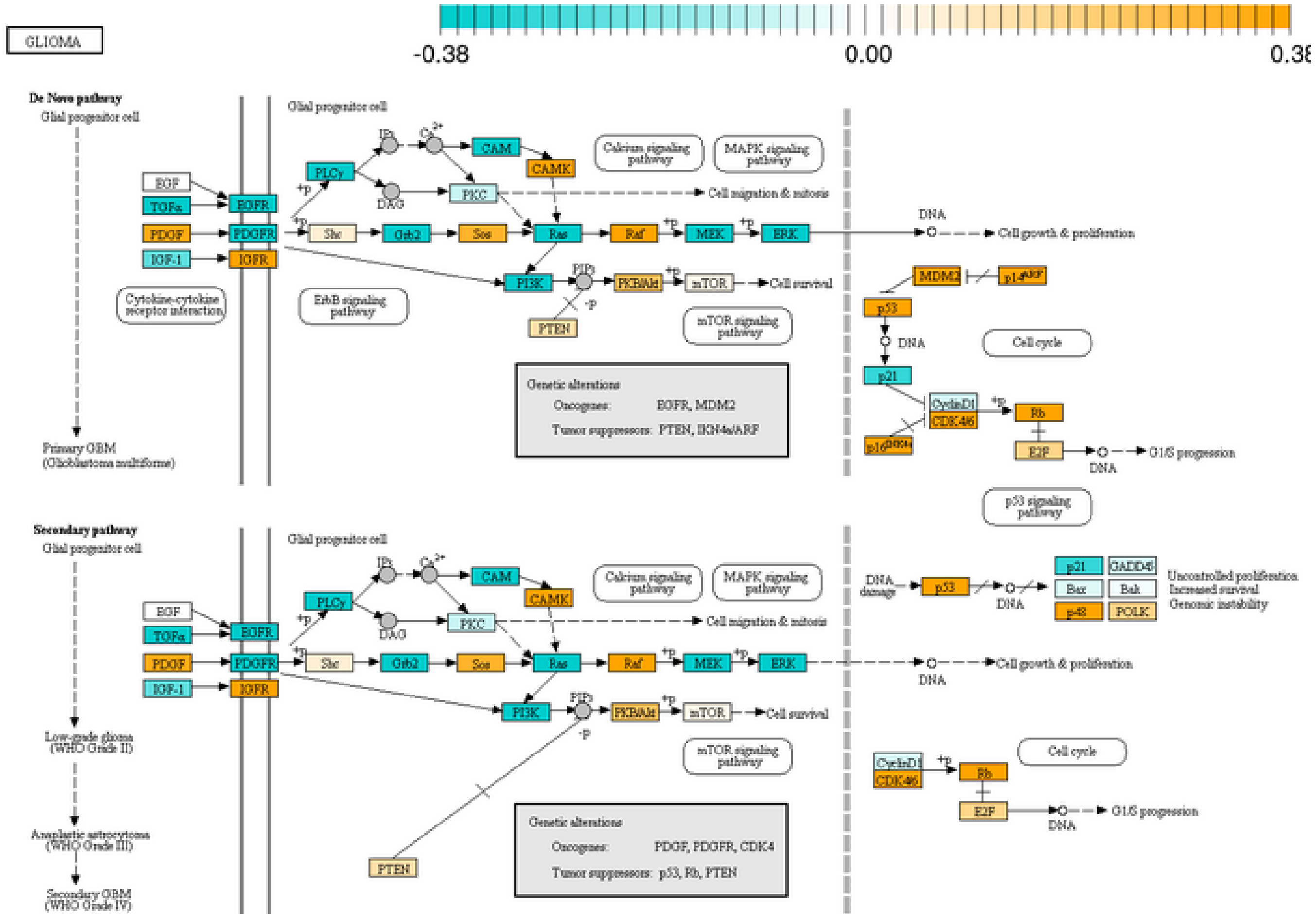
GLIOMA
-0.38
0.00
0.38
De Novo pathway
Glial progenitor cell
EGF
TGFα
PDGF
IGF-1
EGFR
PDGFR
IGFR
Cytokine-cytokine receptor interaction
Primary GBM (Glioblastoma multiforme)
Glial progenitor cell
PLCγ
IP3
Ca2+
CAM
CAMK
DAG
PKC
Calcium signaling pathway
MAPK signaling pathway
Cell migration & mitosis
Shc
Grb2
Sos
Ras
Raf
MEK
ERK
DNA
Cell growth & proliferation
PI3K
PIP3
PKB/Akt
mTOR
Cell survival
PTEN
ErbB signaling pathway
mTOR signaling pathway
Genetic alterations
Oncogenes: EGFR, MDM2
Tumor suppressors: PTEN, INK4a/ARF
MDM2
p14ARF
p53
p21
Cell cycle
CyclinD1
CDK4/6
Rb
p16INK4a
E2F
G1/S progression
p53 signaling pathway
Secondary pathway
Glial progenitor cell
Low-grade glioma (WHO Grade II)
Anaplastic astrocytoma (WHO Grade III)
Secondary GBM (WHO Grade IV)
DNA damage
Bax
Bak
GADD45
p48
POLK
Uncontrolled proliferation
Increased survival
Genomic instability
Oncogenes: PDGF, PDGFR, CDK4
Tumor suppressors: p53, Rb, PTEN

E

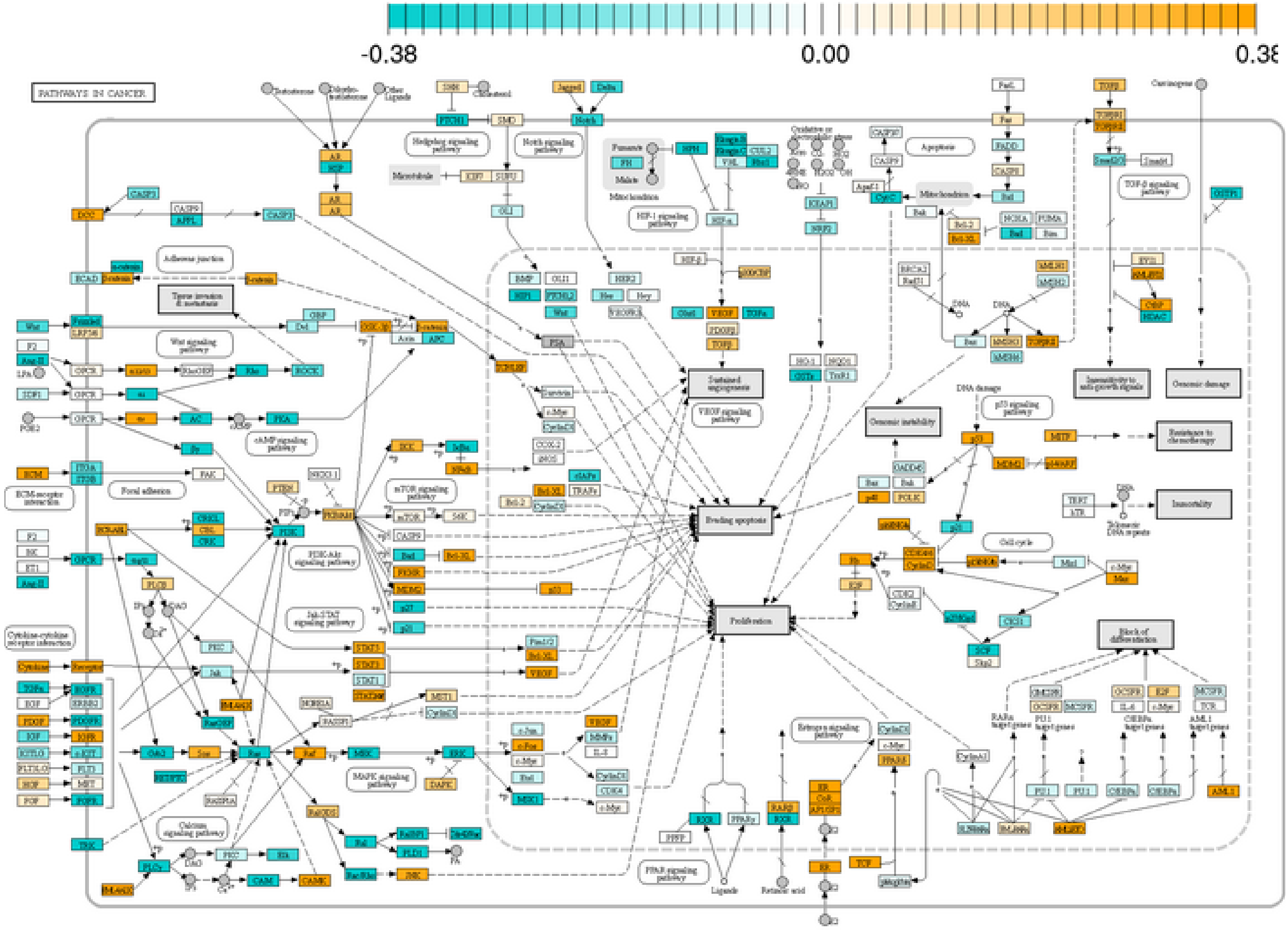
-0.38
0.00
0.38
PATHWAYS IN CANCER
Evading apoptosis
Proliferation
Sustained angiogenesis
Genomic instability
Immortality
Block of differentiation
Resistance to chemotherapy

F

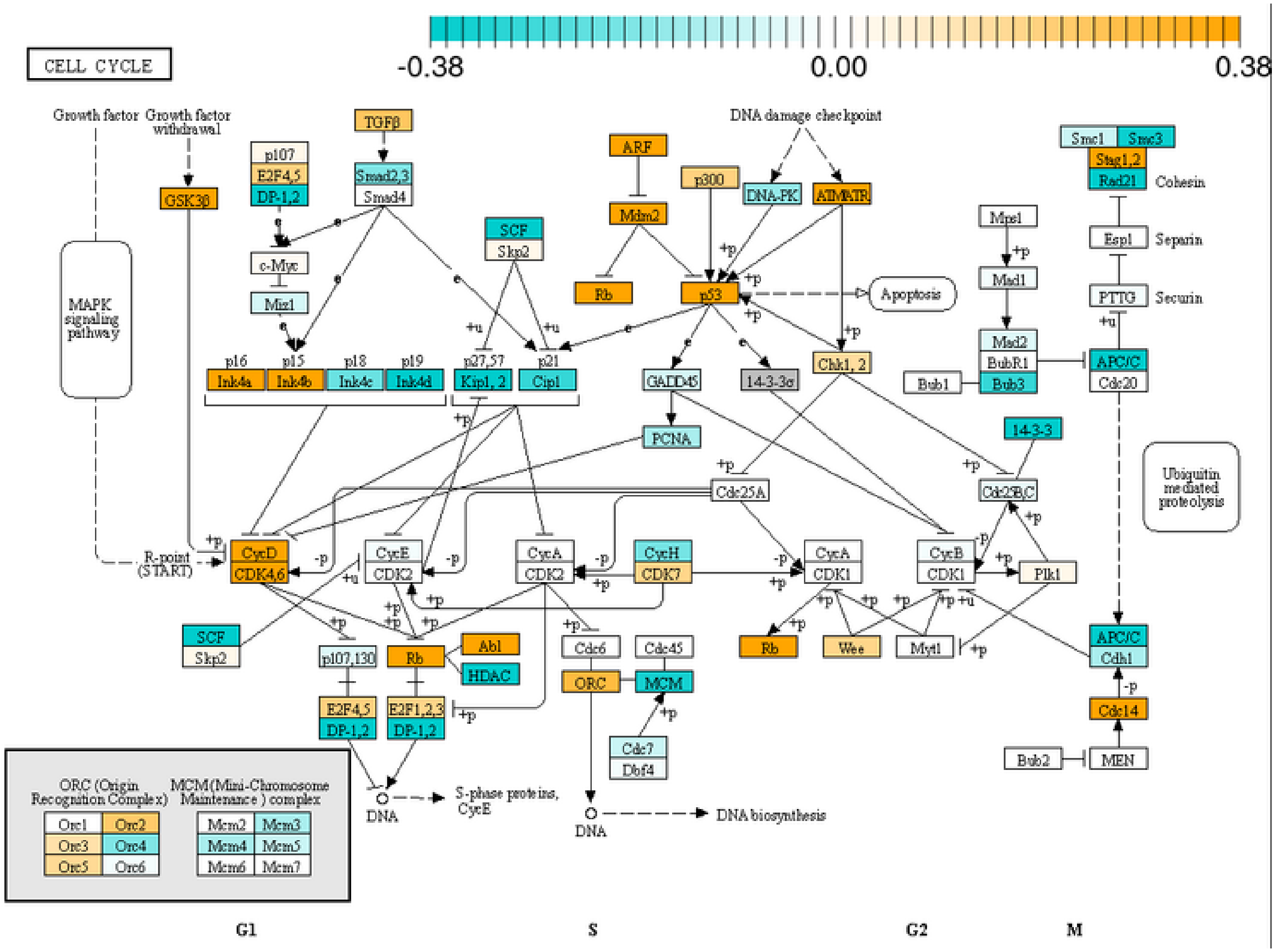
-0.38
0.00
0.38
CELL CYCLE
Growth factor
Growth factor withdrawal
TGFβ
DNA damage checkpoint
p107
E2F4,5
DP-1,2
Smad2,3
Smad4
GSK3β
ARF
Mdm2
p300
DNA-PK
ATM/ATR
Smc1
Smc3
Stag1,2
Rad21
Cohesin
c-Myc
Miz1
SCF
Skp2
Rb
p53
Apoptosis
Mps1
Mad1
Espl
Separin
PTTG
Securin
MAPK signaling pathway
p16
p15
p18
p19
Ink4a
Ink4b
Ink4c
Ink4d
p27,57
p21
Kip1, 2
Cip1
GADD45
14-3-3σ
Chk1, 2
Mad2
BubR1
Bub3
Bub1
APC/C
Cdc20
PCNA
14-3-3
Cdc25A
Cdc25B,C
Ubiquitin mediated proteolysis
R-point (START)
CycD
CDK4,6
CycE
CDK2
CycA
CDK2
CycH
CDK7
CycA
CDK1
CycB
CDK1
Plk1
SCF
Skp2
p107,130
Rb
Abl
HDAC
Cdc6
Cdc45
ORC
MCM
Rb
Wee
Myt1
APC/C
Cdh1
E2F4,5
DP-1,2
E2F1,2,3
DP-1,2
Cdc7
Dbf4
Cdc14
Bub2
MEN
ORC (Origin Recognition Complex)
MCM (Mini-Chromosome Maintenance) complex
Orc1
Orc2
Orc3
Orc4
Orc5
Orc6
Mcm2
Mcm3
Mcm4
Mcm5
Mcm6
Mcm7
DNA
S-phase proteins, CycE
DNA
DNA biosynthesis
G1
S
G2
M

G

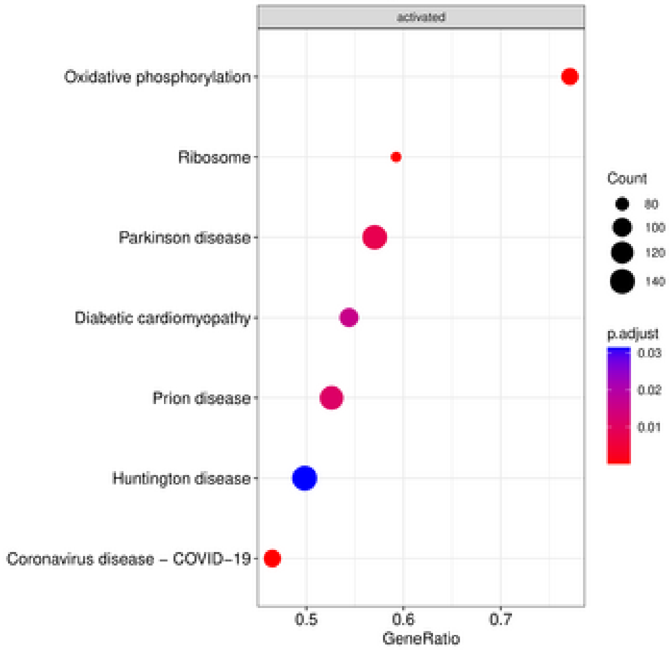
activated
Oxidative phosphorylation
Ribosome
Parkinson disease
Diabetic cardiomyopathy
Prion disease
Huntington disease
Coronavirus disease – COVID–19
0.5
0.6
0.7
GeneRatio
Count
80
100
120
140
p.adjust
0.03
0.02
0.01

H

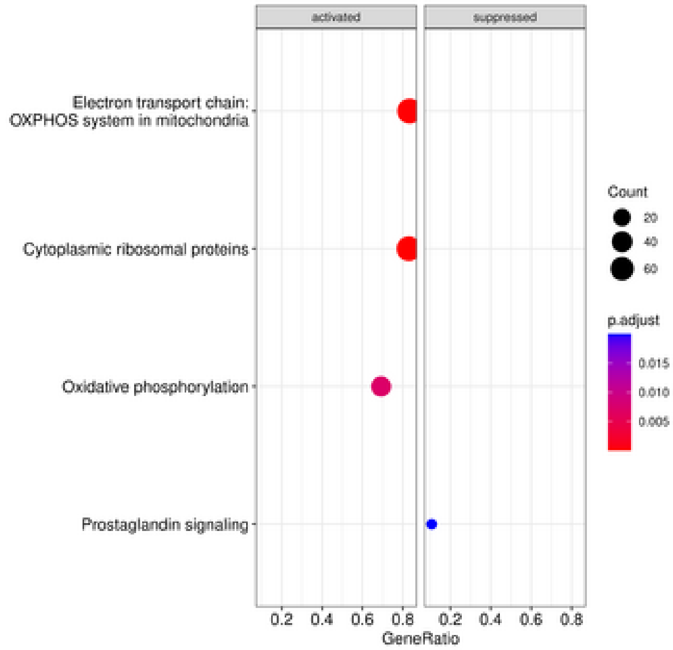
activated
suppressed
Electron transport chain: OXPHOS system in mitochondria
Cytoplasmic ribosomal proteins
Oxidative phosphorylation
Prostaglandin signaling
0.2
0.4
0.6
0.8
0.2
0.4
0.6
0.8
GeneRatio
Count
20
40
60
p.adjust
0.015
0.010
0.005

I

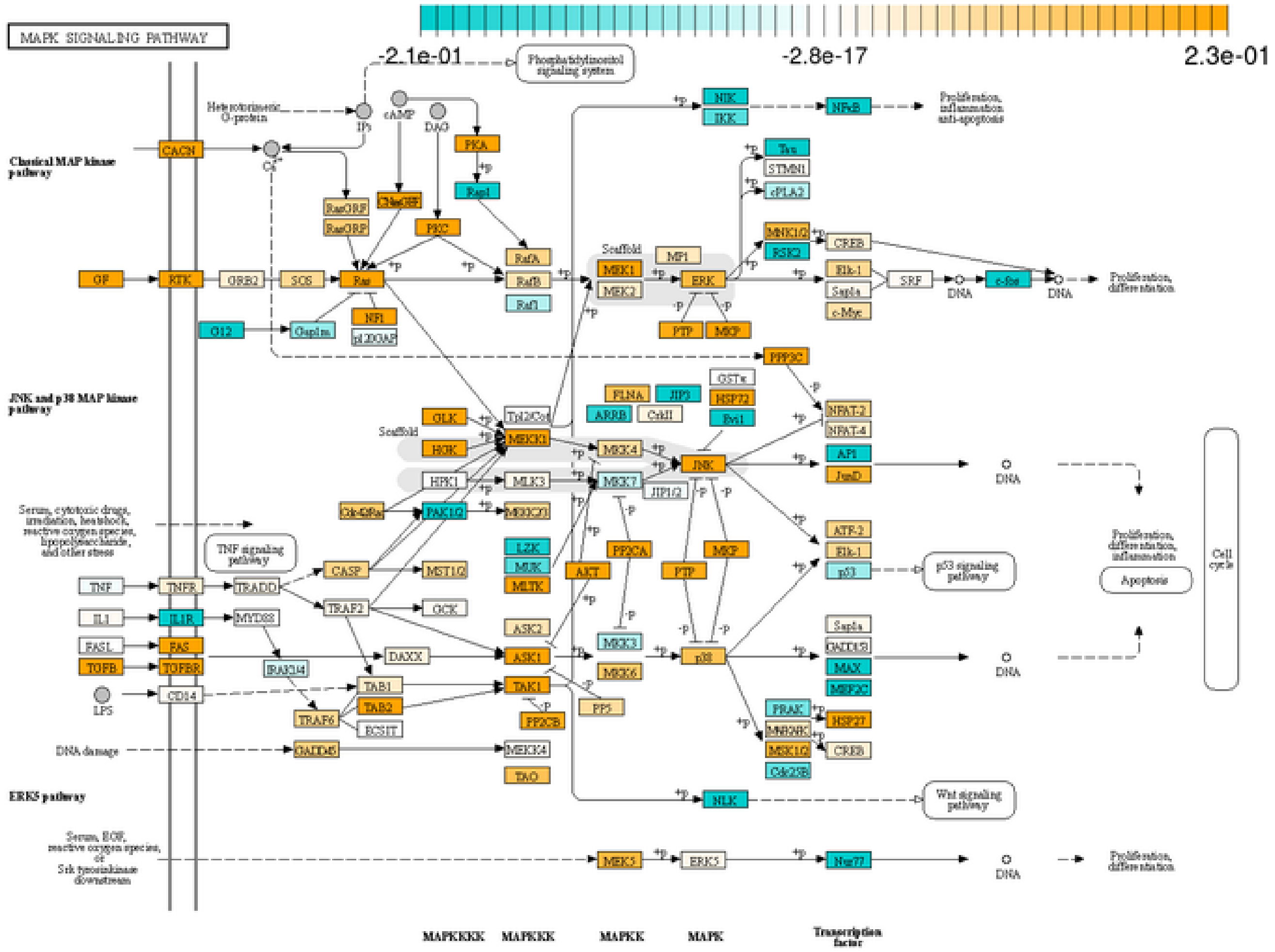
MAPK SIGNALING PATHWAY
-2.1e-01
-2.8e-17
2.3e-01
Classical MAP kinase pathway
JNK and p38 MAP kinase pathway
ERK5 pathway
Phosphatidylinositol signaling system
Proliferation, inflammation, anti-apoptosis
Proliferation, differentiation
Proliferation, differentiation, inflammation
Apoptosis
Cell cycle
TNF signaling pathway
p53 signaling pathway
Wnt signaling pathway
DNA damage
DNA
MAPKKK
MAPKKK
MAPKK
MAPK
Transcription factor

J

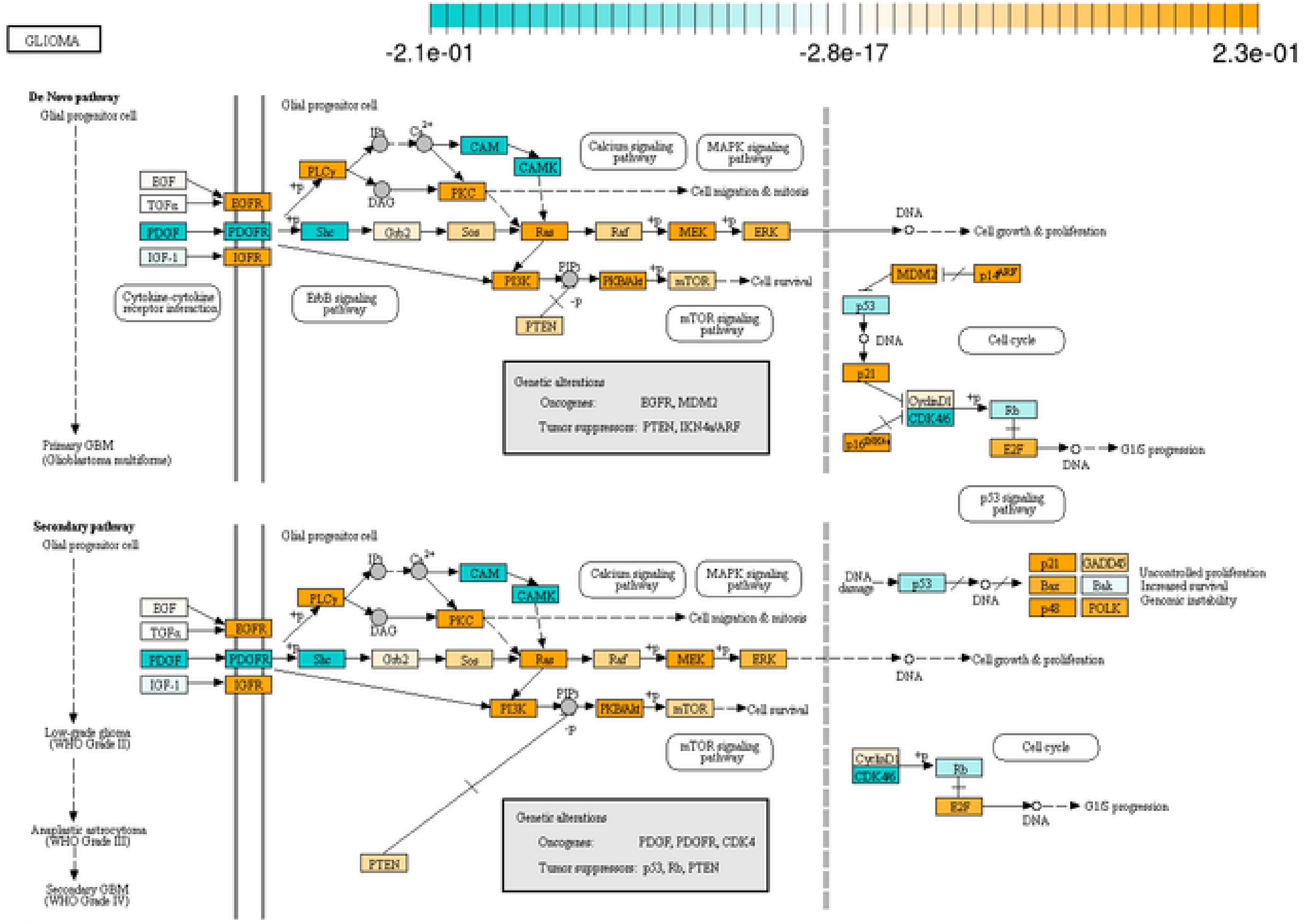
GLIOMA
-2.1e-01
-2.8e-17
2.3e-01
De Novo pathway
Glial progenitor cell
Primary GBM (Glioblastoma multiforme)
Calcium signaling pathway
MAPK signaling pathway
Cell migration & mitosis
Cell growth & proliferation
Cell survival
Cytokine-cytokine receptor interaction
ErbB signaling pathway
mTOR signaling pathway
Cell cycle
G1/S progression
p53 signaling pathway
Genetic alterations
Oncogenes: EGFR, MDM2
Tumor suppressors: PTEN, INK4a/ARF
Secondary pathway
Low-grade glioma (WHO Grade II)
Anaplastic astrocytoma (WHO Grade III)
Secondary GBM (WHO Grade IV)
Uncontrolled proliferation
Increased survival
Genomic instability
Oncogenes: PDGF, PDGFR, CDK4
Tumor suppressors: p53, Rb, PTEN

K

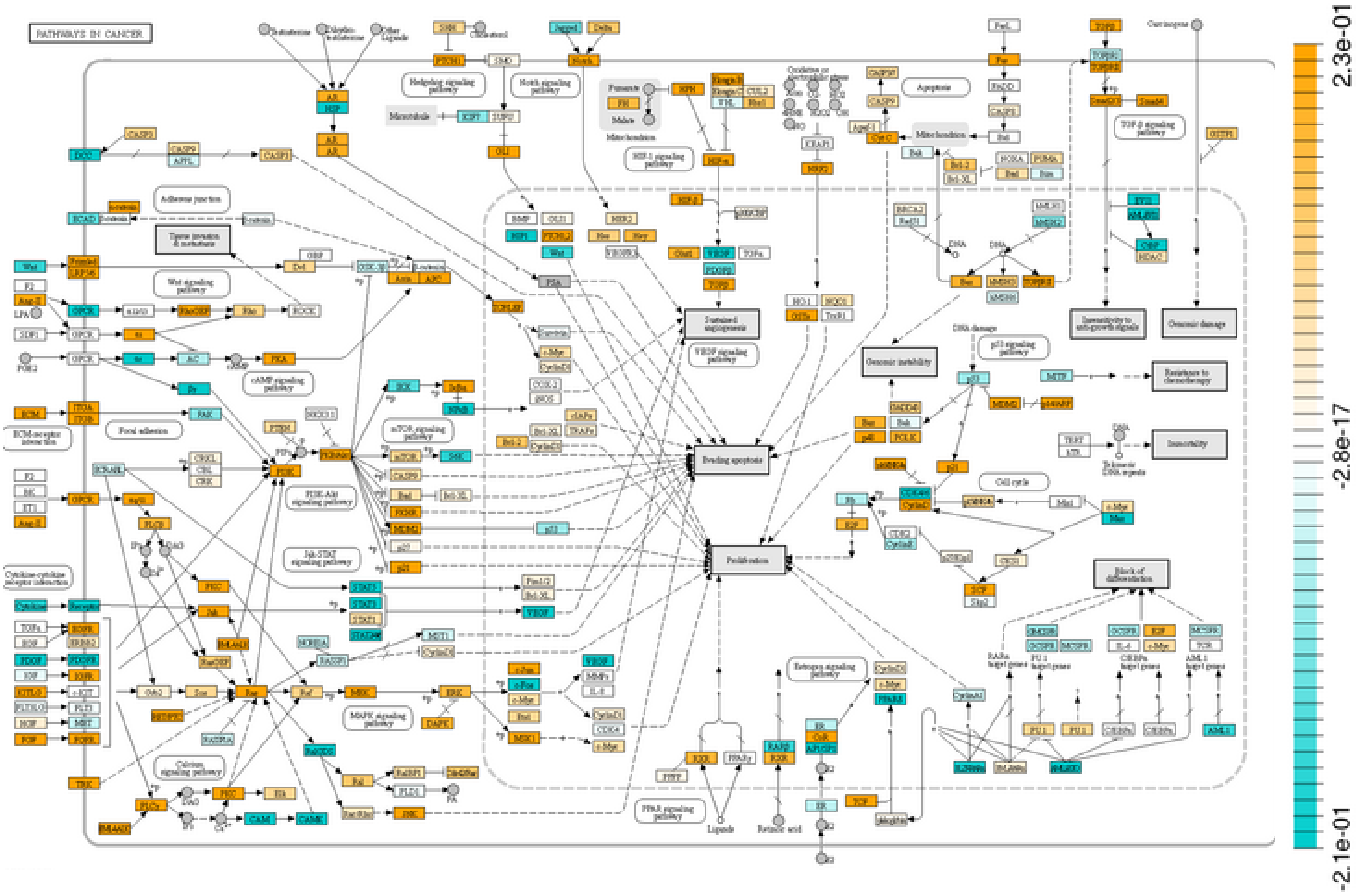
PATHWAYS IN CANCER
2.3e-01
-2.8e-17
-2.1e-01

L

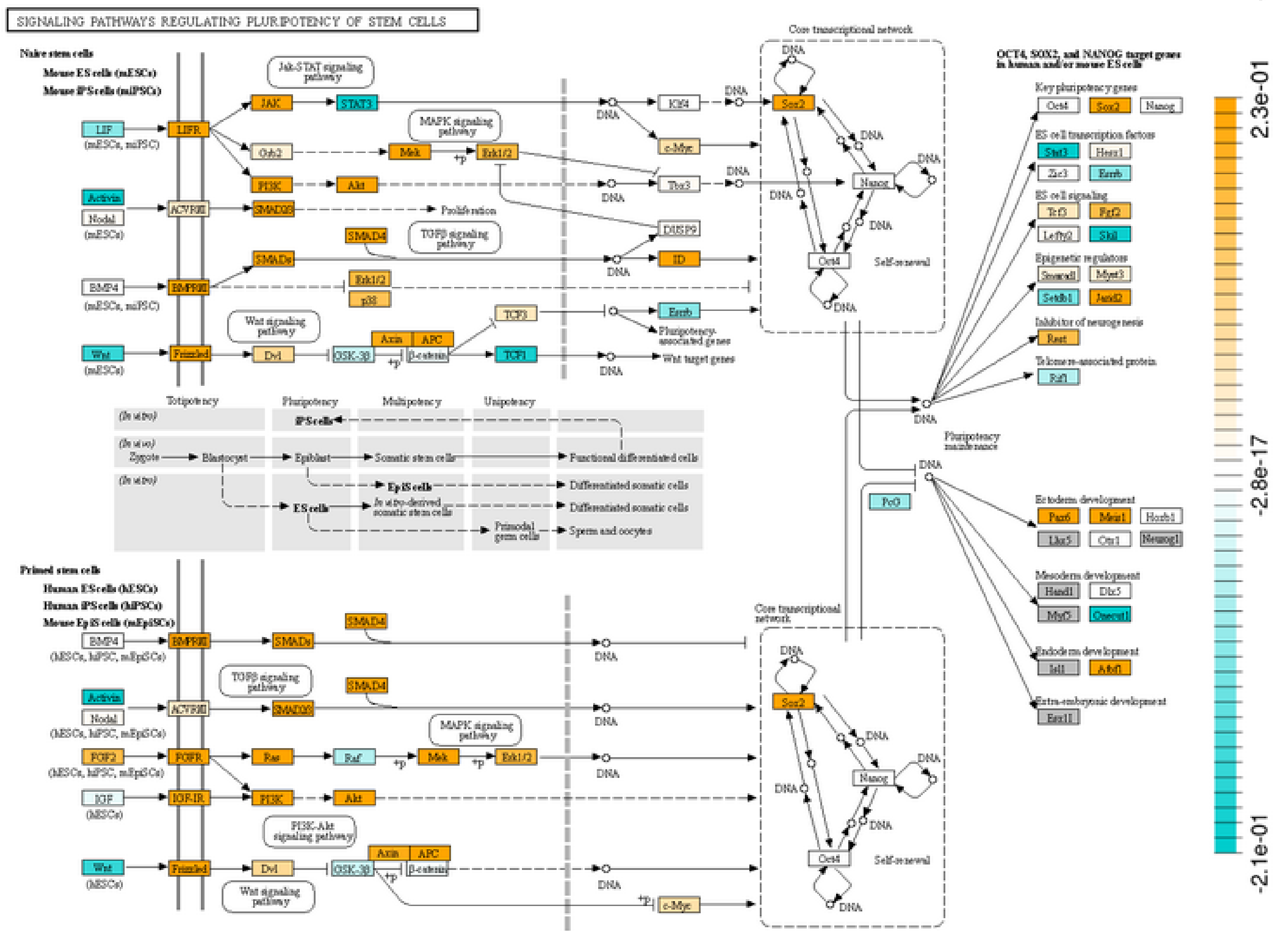
SIGNALING PATHWAYS REGULATING PLURIPOTENCY OF STEM CELLS
Core transcriptional network
Self-renewal
2.3e-01
-2.8e-17
-2.1e-01

Supplementary Figure 4: Ganglioglioma snRNA-seq gene set enrichment analysis.
(A-B) Dot plots of top 10 (or fewer if Bonferroni-adjusted p-value>0.05) most activated or suppressed KEGG (A), or Wiki (B) pathways when comparing neoplastic-appearing cells as a whole to normal-appearing neural cells. Bonferroni-adjusted-p-value shown as well as gene count in each pathway.
(C-F) Pathview representations of RNA average log2-fold-change from comparing neoplastic-appearing cells as a whole to normal-appearing neural cells for select KEGG pathways: MAPK signaling (C), glioma (D), cancer signaling (E), and cell cycle (F).
(G-H) Dotplots of top most activated or suppressed KEGG (G) or Wiki (H) pathways when comparing *CD34+* neoplastic-appearing cells as a whole to *CD34-* neoplastic-appearing cells. Bonferroni-adjusted-p-value shown as well as gene count in each pathway.
(I-L) Pathview representations of RNA average log2-fold-change from comparing *CD34+* neoplastic-appearing cells as a whole to *CD34-* neoplastic-appearing cells for select KEGG pathways: MAPK signaling (I), glioma (J), cancer signaling (K), and pluripotency (L).

A

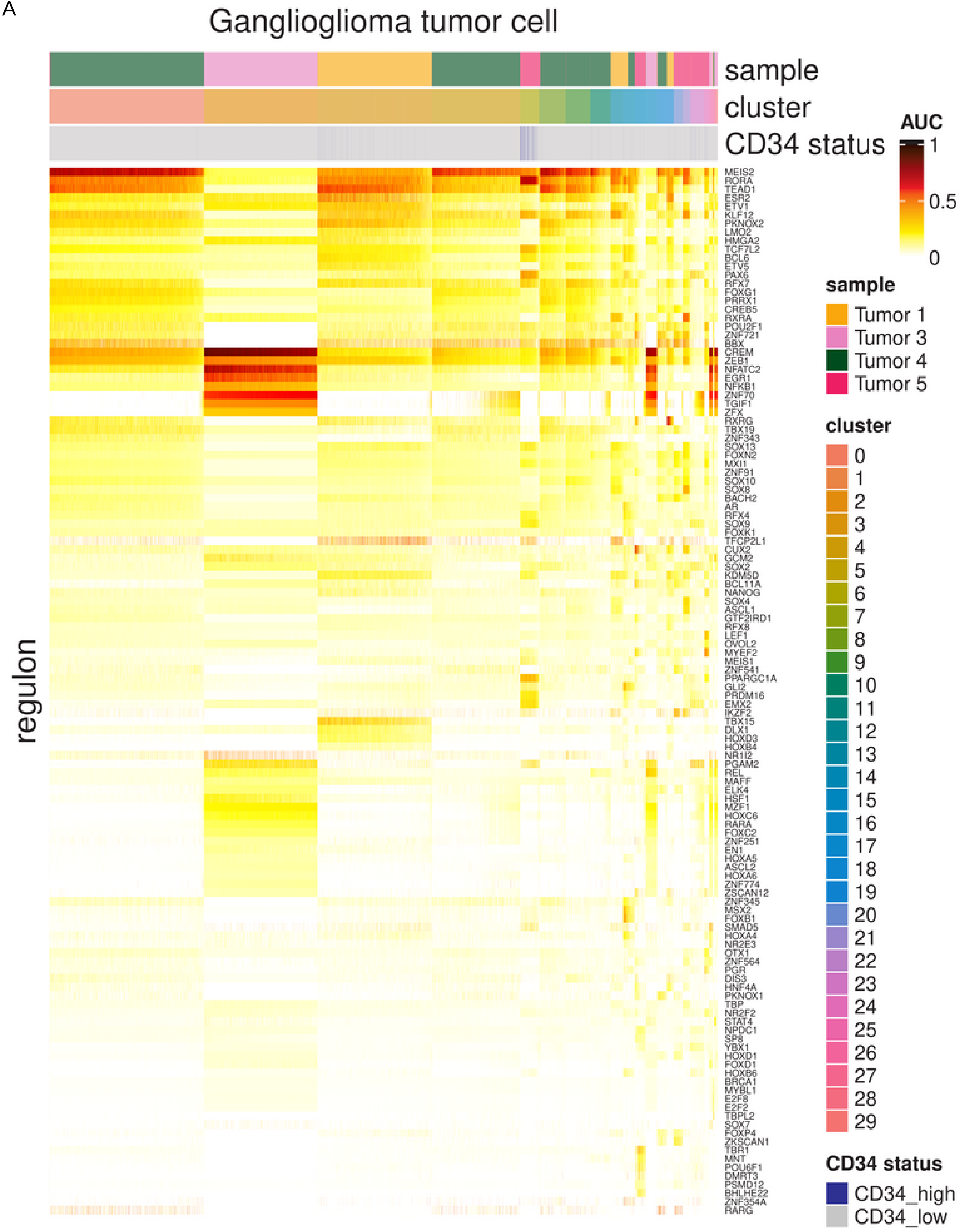
Ganglioglioma tumor cell
sample
cluster
CD34 status
AUC
1
0.5
0
regulon
MEIS2
RORA
TEAD1
ESR2
ETV1
KLF12
PKNOX2
LMO2
HMGA2
TCF7L2
BCL6
ETV5
PAX6
RFX7
FOXG1
PRRX1
CREB5
RXRA
POU2F1
ZNF721
BBX
CREM
ZEB1
NFATC2
EGR1
NFKB1
ZNF70
TGIF1
ZFX
RXRG
TBX19
ZNF343
SOX13
FOXN2
MXI1
ZNF91
SOX10
SOX8
BACH2
AR
RFX4
SOX9
FOXK1
TFCP2L1
CUX2
GCM2
SOX2
KDM5D
BCL11A
NANOG
SOX4
ASCL1
GTF2IRD1
RFX8
LEF1
OVOL2
MYEF2
MEIS1
ZNF541
PPARGC1A
GLI2
PRDM16
EMX2
IKZF2
TBX15
DLX1
HOXD3
HOXB4
NR1I2
PGAM2
REL
MAFF
ELK4
HSF1
MZF1
HOXC6
RARA
FOXC2
ZNF251
EN1
HOXA5
ASCL2
HOXA6
ZNF774
ZSCAN12
ZNF345
MSX2
FOXB1
SMAD5
HOXA4
NR2E3
OTX1
ZNF564
PGR
DIS3
HNF4A
PKNOX1
TBP
NR2F2
STAT4
NPDC1
SP8
YBX1
HOXD1
FOXD1
HOXB6
BRCA1
MYBL1
E2F8
E2F2
TBPL2
SOX7
FOXP4
ZKSCAN1
TBR1
MNT
POU6F1
DMRT3
PSMD12
BHLHE22
ZNF354A
RARG
sample
Tumor 1
Tumor 3
Tumor 4
Tumor 5
cluster
CD34 status
CD34_high
CD34_low

B

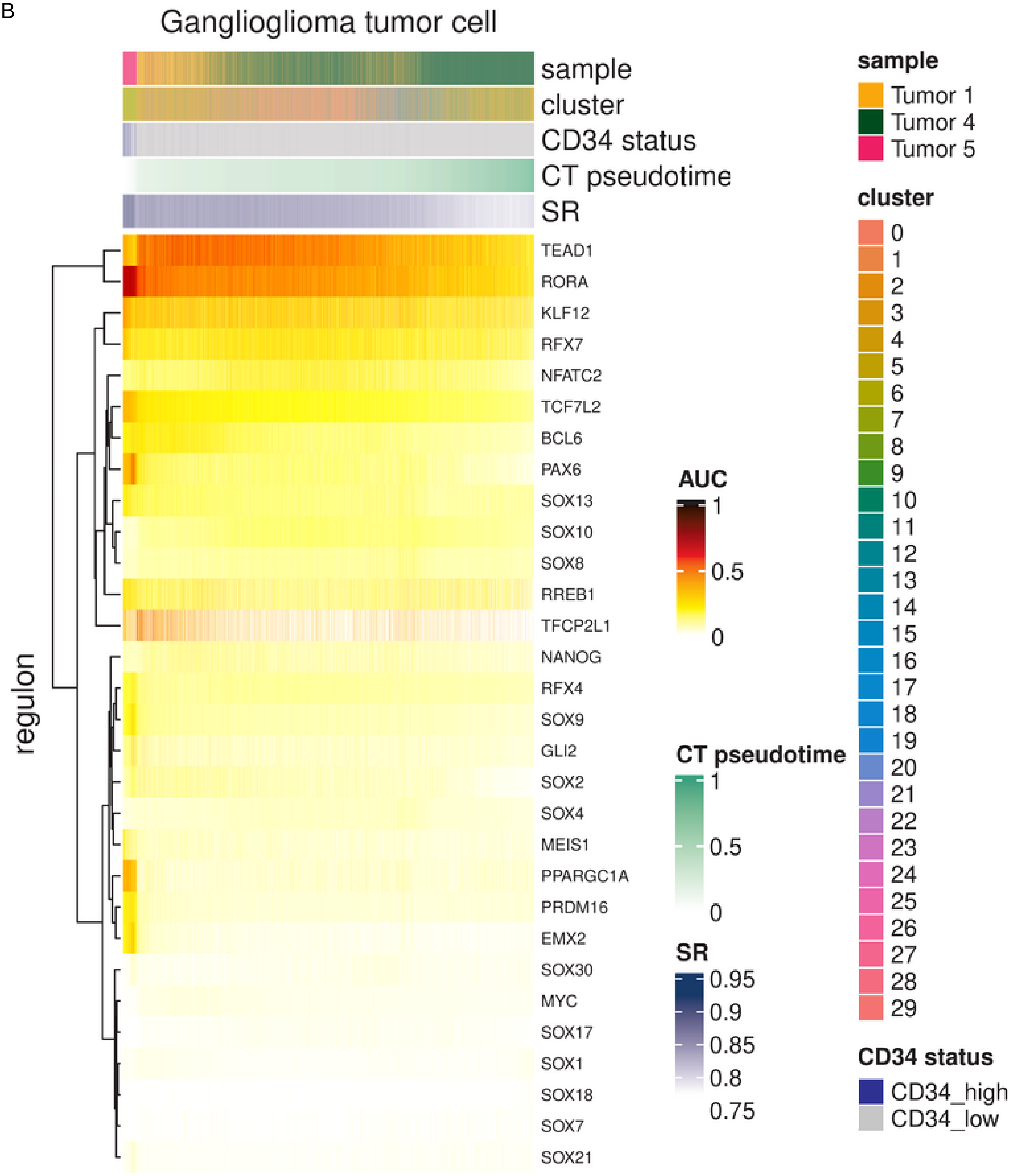

Ganglioglioma tumor cell
sample
cluster
CD34 status
CT pseudotime
SR
regulon
TEAD1
RORA
KLF12
RFX7
NFATC2
TCF7L2
BCL6
PAX6
SOX13
SOX10
SOX8
RREB1
TFCP2L1
NANOG
RFX4
SOX9
GLI2
SOX2
SOX4
MEIS1
PPARGC1A
PRDM16
EMX2
SOX30
MYC
SOX17
SOX1
SOX18
SOX7
SOX21
AUC
1
0.5
0
CT pseudotime
1
0.5
0
SR
0.95
0.9
0.85
0.8
0.75
sample
Tumor 1
Tumor 4
Tumor 5
cluster
CD34 status
CD34_high
CD34_low

C

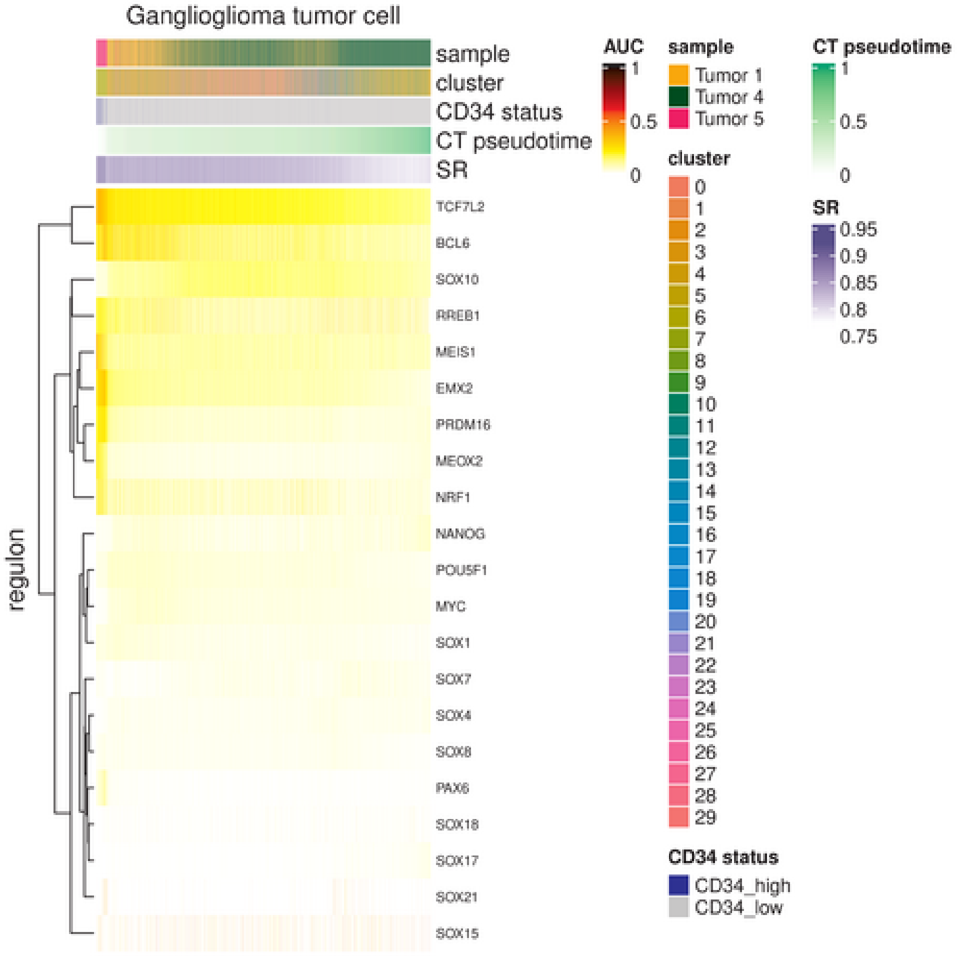
Ganglioglioma tumor cell
sample
cluster
CD34 status
CT pseudotime
SR
regulon
TCF7L2
BCL6
SOX10
RREB1
MEIS1
EMX2
PRDM16
MEOX2
NRF1
NANOG
POU5F1
MYC
SOX1
SOX7
SOX4
SOX8
PAX6
SOX18
SOX17
SOX21
SOX15
AUC
1
0.5
0
sample
Tumor 1
Tumor 4
Tumor 5
cluster
CD34 status
CD34_high
CD34_low
CT pseudotime
1
0.5
0
SR
0.95
0.9
0.85
0.8
0.75

D

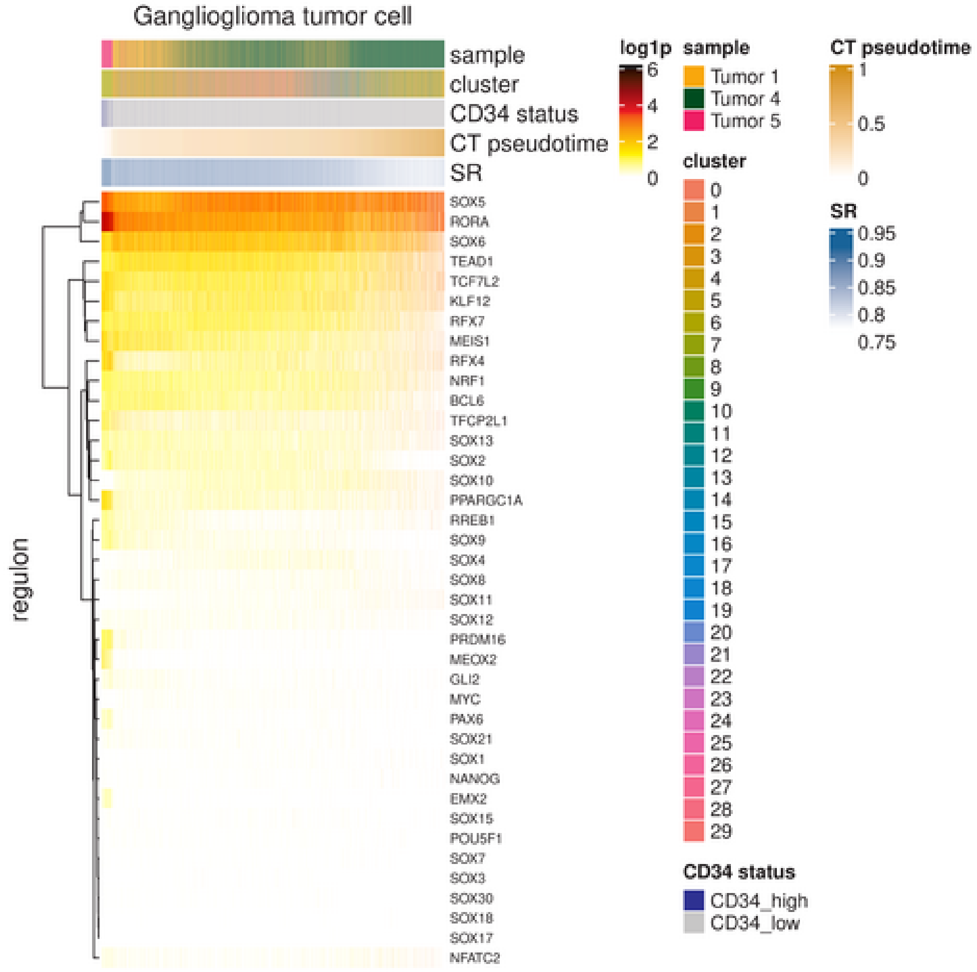
Ganglioglioma tumor cell
sample
cluster
CD34 status
CT pseudotime
SR
regulon
SOX5
RORA
SOX6
TEAD1
TCF7L2
KLF12
RFX7
MEIS1
RFX4
NRF1
BCL6
TFCP2L1
SOX13
SOX2
SOX10
PPARGC1A
RREB1
SOX9
SOX4
SOX8
SOX11
SOX12
PRDM16
MEOX2
GLI2
MYC
PAX6
SOX21
SOX1
NANOG
EMX2
SOX15
POU5F1
SOX7
SOX3
SOX30
SOX18
SOX17
NFATC2
log1p
6
4
2
0
sample
Tumor 1
Tumor 4
Tumor 5
cluster
CD34 status
CD34_high
CD34_low
CT pseudotime
1
0.5
0
SR
0.95
0.9
0.85
0.8
0.75

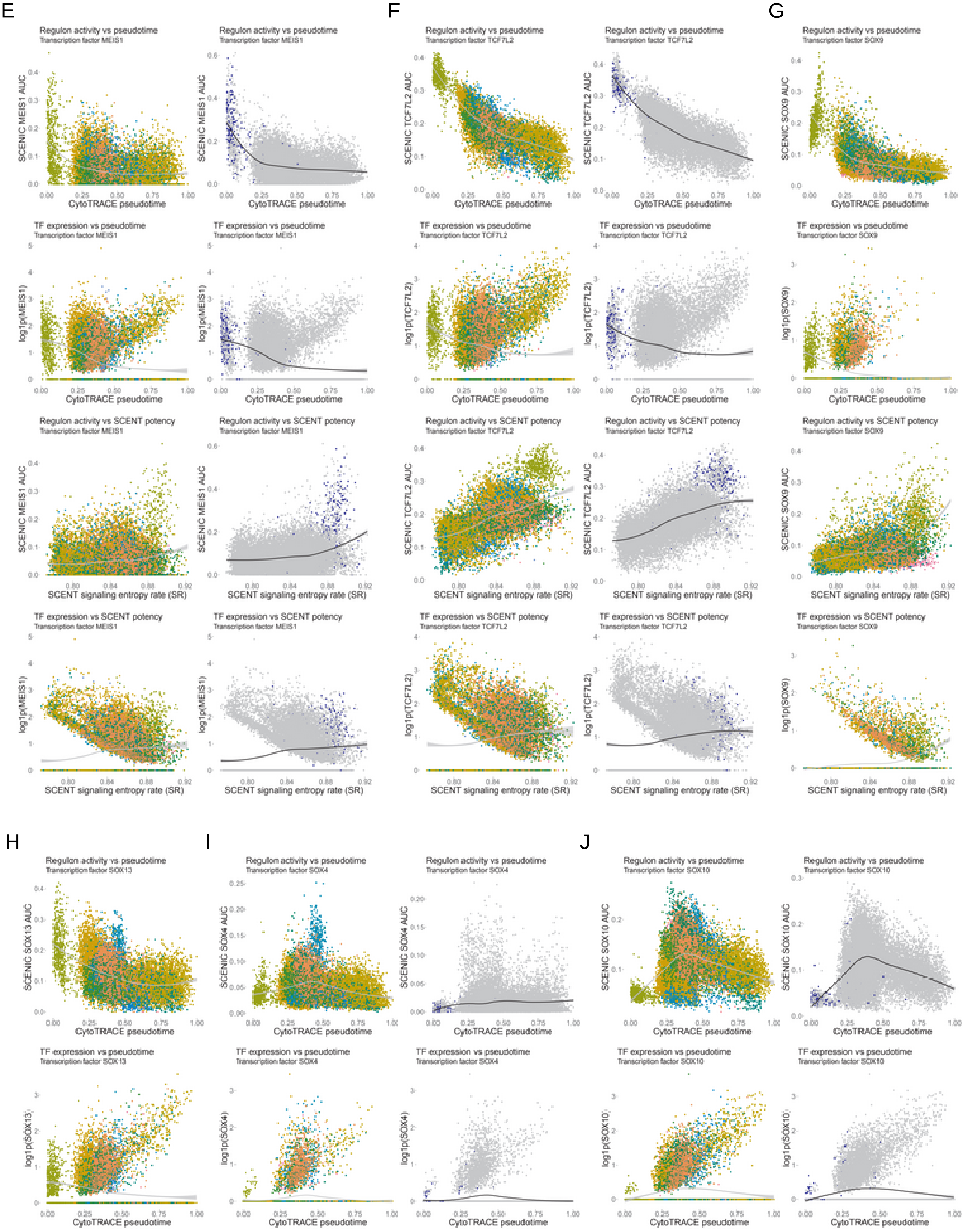
E
Regulon activity vs pseudotime
Transcription factor MEIS1
SCENIC MEIS1 AUC
CytoTRACE pseudotime
TF expression vs pseudotime
log1p(MEIS1)
Regulon activity vs SCENT potency
SCENT signaling entropy rate (SR)
TF expression vs SCENT potency
F
Regulon activity vs pseudotime
Transcription factor TCF7L2
SCENIC TCF7L2 AUC
CytoTRACE pseudotime
TF expression vs pseudotime
log1p(TCF7L2)
Regulon activity vs SCENT potency
SCENT signaling entropy rate (SR)
TF expression vs SCENT potency
G
Regulon activity vs pseudotime
Transcription factor SOX9
SCENIC SOX9 AUC
CytoTRACE pseudotime
TF expression vs pseudotime
log1p(SOX9)
Regulon activity vs SCENT potency
SCENT signaling entropy rate (SR)
TF expression vs SCENT potency
H
Regulon activity vs pseudotime
Transcription factor SOX13
SCENIC SOX13 AUC
CytoTRACE pseudotime
TF expression vs pseudotime
log1p(SOX13)
I
Regulon activity vs pseudotime
Transcription factor SOX4
SCENIC SOX4 AUC
CytoTRACE pseudotime
TF expression vs pseudotime
log1p(SOX4)
J
Regulon activity vs pseudotime
Transcription factor SOX10
SCENIC SOX10 AUC
CytoTRACE pseudotime
TF expression vs pseudotime
log1p(SOX10)

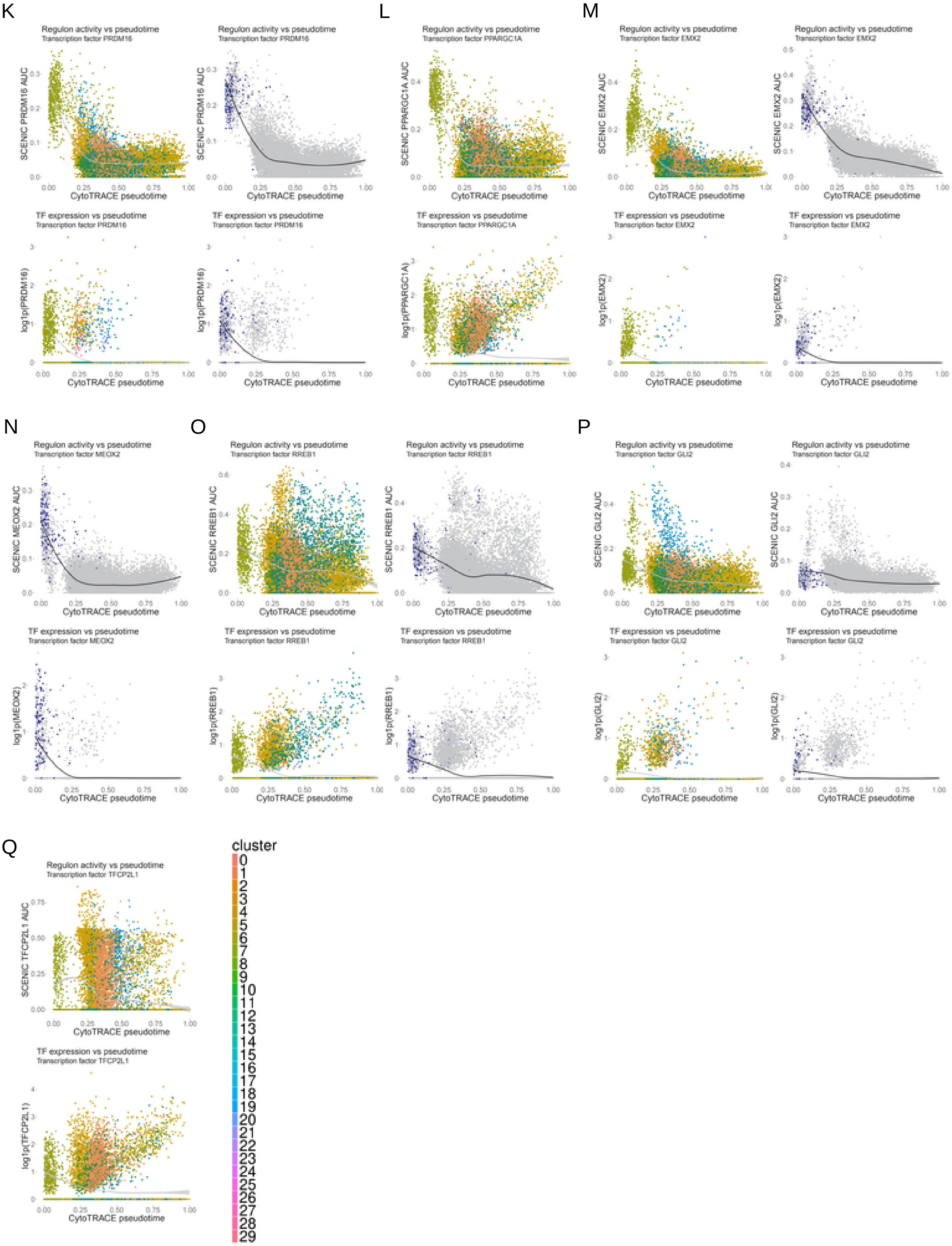
K
Regulon activity vs pseudotime
Transcription factor PRDM16
SCENIC PRDM16 AUC
CytoTRACE pseudotime
TF expression vs pseudotime
Transcription factor PRDM16
log1p(PRDM16)
L
Regulon activity vs pseudotime
Transcription factor PPARGC1A
SCENIC PPARGC1A AUC
TF expression vs pseudotime
Transcription factor PPARGC1A
log1p(PPARGC1A)
M
Regulon activity vs pseudotime
Transcription factor EMX2
SCENIC EMX2 AUC
TF expression vs pseudotime
Transcription factor EMX2
log1p(EMX2)
N
Regulon activity vs pseudotime
Transcription factor MEOX2
SCENIC MEOX2 AUC
TF expression vs pseudotime
Transcription factor MEOX2
log1p(MEOX2)
O
Regulon activity vs pseudotime
Transcription factor RREB1
SCENIC RREB1 AUC
TF expression vs pseudotime
Transcription factor RREB1
log1p(RREB1)
P
Regulon activity vs pseudotime
Transcription factor GLI2
SCENIC GLI2 AUC
TF expression vs pseudotime
Transcription factor GLI2
log1p(GLI2)
Q
Regulon activity vs pseudotime
Transcription factor TFCP2L1
SCENIC TFCP2L1 AUC
TF expression vs pseudotime
Transcription factor TFCP2L1
log1p(TFCP2L1)
CytoTRACE pseudotime
cluster

Supplementary Figure 5: Ganglioglioma significant transcription factors.
(A) Heatmap of SCENIC AUC for top differentially acting transcription factors from the neural(-like) clusters for each ganglioglioma neural(-like) cell.
(B) Heatmap of neoplastic cell cluster-based SCENIC AUC for the union of top cluster 6-associated transcription factors with SOX family and neuroectodermal (NANOG, POU5F1, PAX6, and MYC) transcription factors of interest *a priori*, ordered by CytoTRACE pseudotime. Note tumor 3 data was omitted due to low complexity confounding pseudotime calculation.
(C) Analogous to panel B but for *CD34* status-based SCENIC AUC and *CD34*+ associated factors.
(D) Transcription factor expression for the union of factors from panels B-C.
(E) Scatterplots of MEIS1 cluster-based SCENIC AUC and TF expression vs CytoTRACE pseudotime or SCENT signaling entropy rate (SR) with data colored by cluster (left) or (right) *CD34* status-based SCENIC AUC and TF expression vs CytoTRACE pseudotime or SCENT signaling entropy rate (SR) with data colored by *CD34* status. Loess best fit shown as a gray or black curve; gray shading for the 95% confidence interval.
(F-Q) Analogous results to (E) are shown for (F) TCF7L2, (G) SOX9 (cluster-based results only due to *CD34*-based regulon not meeting filters); continued using only CytoTRACE pseudotime for space: (H) SOX13 (only cluster-based), (I) SOX4, (J) SOX10, (K) PRDM16, (L) PPARGC1A (only cluster-based), (M) EMX2, (N) MEOX2 (only CD34-based), (O) RREB1, (P) GLI2, (Q) TFCP2L1 (only cluster-based).

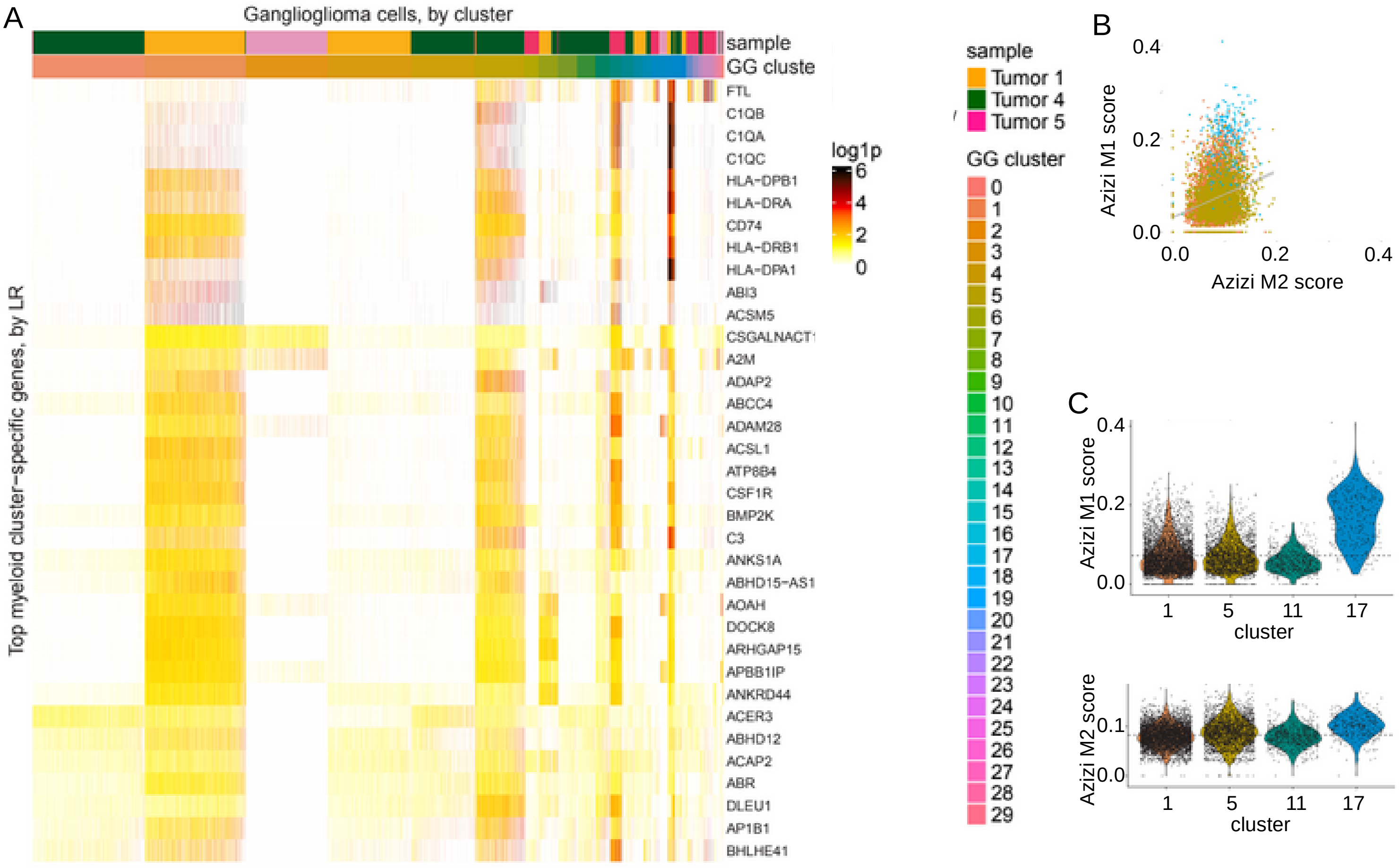
A
Ganglioglioma cells, by cluster
sample
GG cluste
Top myeloid cluster-specific genes, by LR
FTL
C1QB
C1QA
C1QC
HLA-DPB1
HLA-DRA
CD74
HLA-DRB1
HLA-DPA1
ABI3
ACSM5
CSGALNACT1
A2M
ADAP2
ABCC4
ADAM28
ACSL1
ATP8B4
CSF1R
BMP2K
C3
ANKS1A
ABHD15-AS1
AOAH
DOCK8
ARHGAP15
APBB1IP
ANKRD44
ACER3
ABHD12
ACAP2
ABR
DLEU1
AP1B1
BHLHE41
log1p
6
4
2
0
sample
Tumor 1
Tumor 4
Tumor 5
GG cluster
0 1 2 3 4 5 6 7 8 9 10 11 12 13 14 15 16 17 18 19 20 21 22 23 24 25 26 27 28 29
B
Azizi M1 score
Azizi M2 score
0.0
0.2
0.4
C
Azizi M1 score
Azizi M2 score
cluster
1
5
11
17
0.0
0.1
0.2
0.4

D

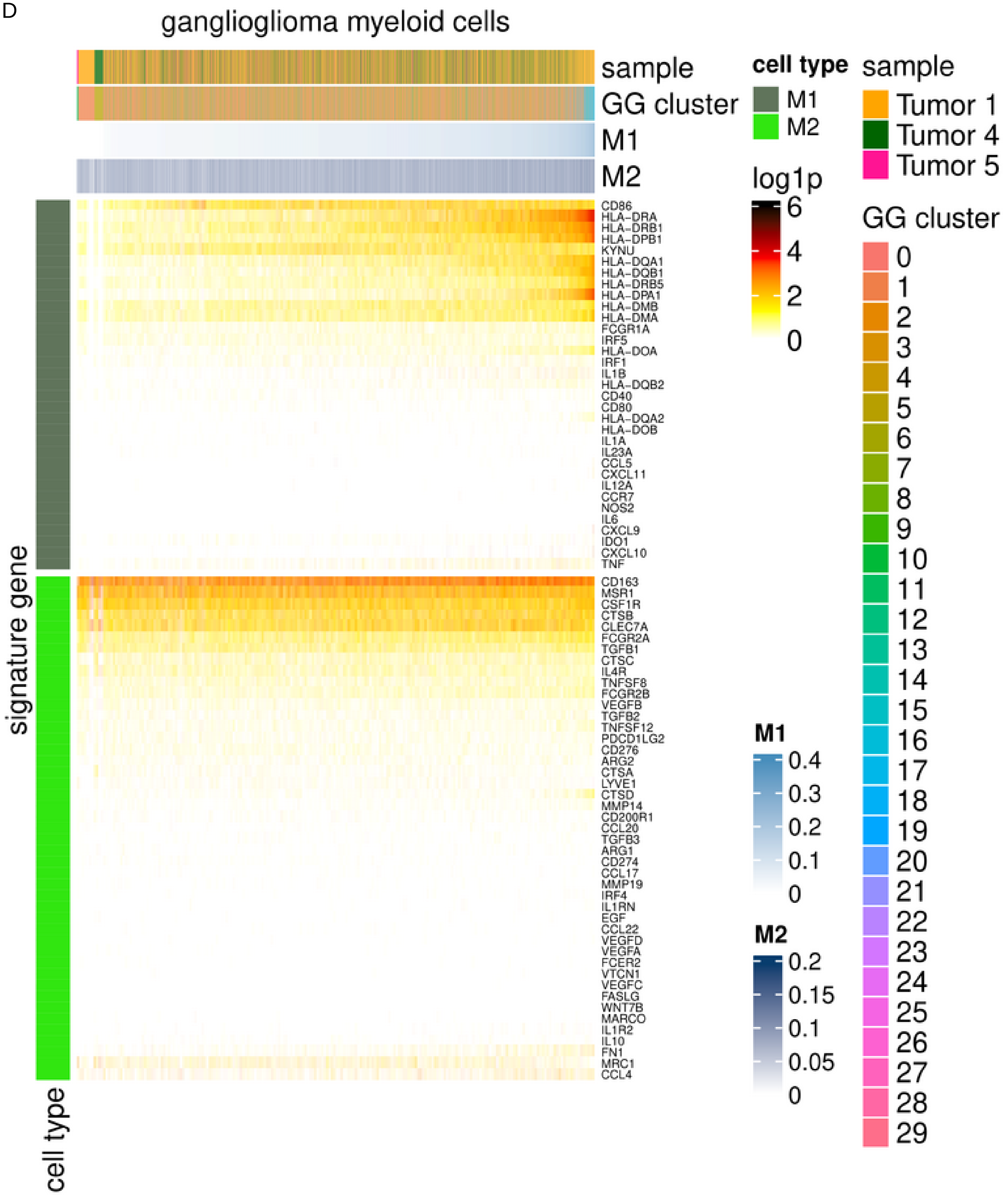
ganglioglioma myeloid cells
sample
GG cluster
M1
M2
cell type
M1
M2
sample
Tumor 1
Tumor 4
Tumor 5
log1p
6
4
2
0
GG cluster
CD86
HLA-DRA
HLA-DRB1
HLA-DPB1
KYNU
HLA-DQA1
HLA-DQB1
HLA-DRB5
HLA-DPA1
HLA-DMB
HLA-DMA
FCGR1A
IRF5
HLA-DOA
IRF1
IL1B
HLA-DQB2
CD40
CD80
HLA-DQA2
HLA-DOB
IL1A
IL23A
CCL5
CXCL11
IL12A
CCR7
NOS2
IL6
CXCL9
IDO1
CXCL10
TNF
CD163
MSR1
CSF1R
CTSB
CLEC7A
FCGR2A
TGFB1
CTSC
IL4R
TNFSF8
FCGR2B
VEGFB
TGFB2
TNFSF12
PDCD1LG2
CD276
ARG2
CTSA
LYVE1
CTSD
MMP14
CD200R1
CCL20
TGFB3
ARG1
CD274
CCL17
MMP19
IRF4
IL1RN
EGF
CCL22
VEGFD
VEGFA
FCER2
VTCN1
VEGFC
FASLG
WNT7B
MARCO
IL1R2
IL10
FN1
MRC1
CCL4
M1
0.4
0.3
0.2
0.1
0
M2
0.2
0.15
0.1
0.05
0
signature gene
cell type

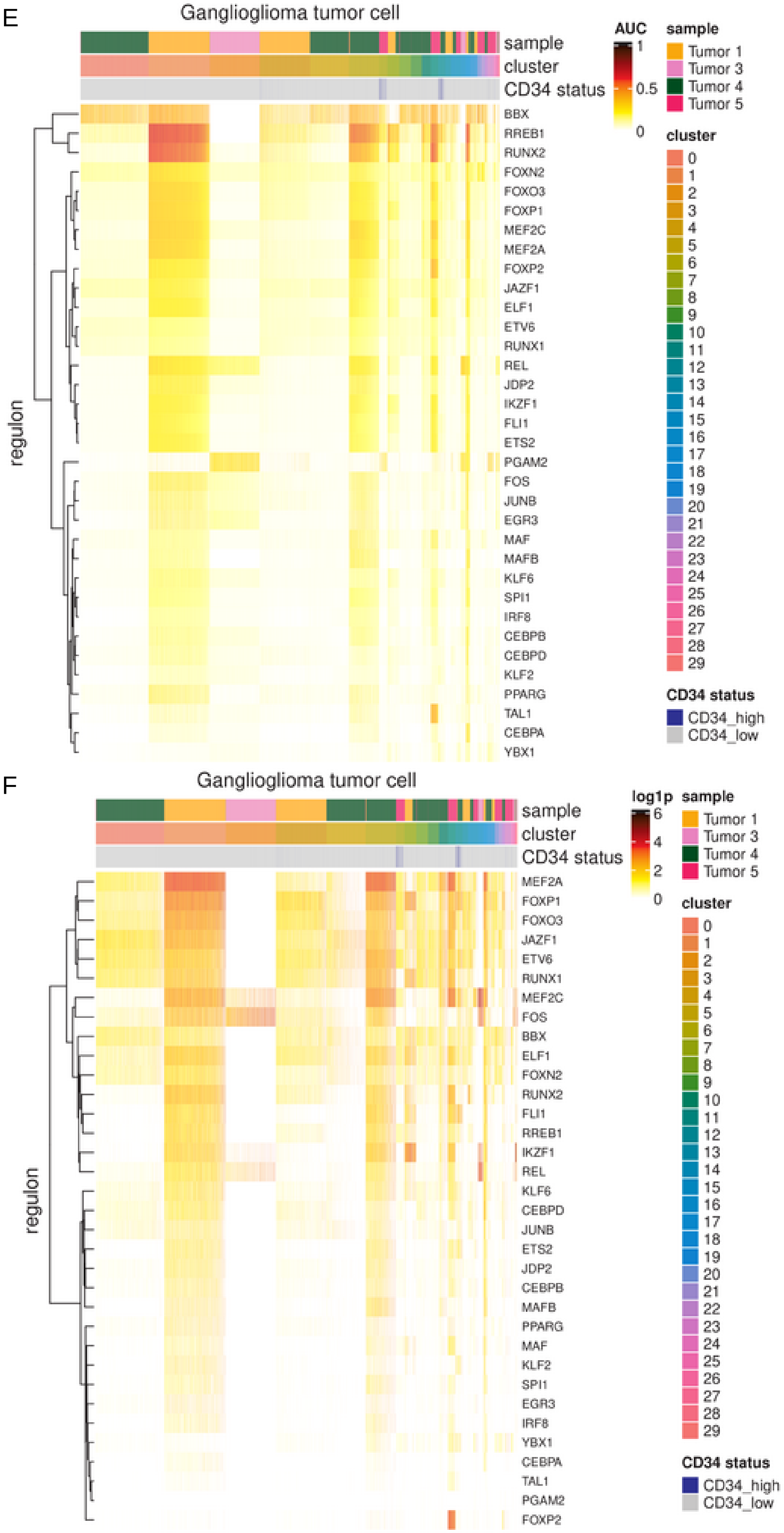

Supplementary Figure 6: Ganglioglioma-associated myeloid cells.
(A) Heatmap of RNA log1p (base 2) for top differentially expressed genes (by likelihood ratio test) for each of the myeloid clusters.
(B-D) Ganglioglioma microglia UCell scoring for Azizi *et al*. M1 vs M2 signature scatterplots (B), UCell scores by ganglioglioma cluster (C), and heatmap of signature component gene log1p (D).
(E-F) Top microglia cluster transcription factors as nominated by SCENIC, shown in a heatmap with SCENIC regulon AUC (E) or transcription factor transcript log1p (F).

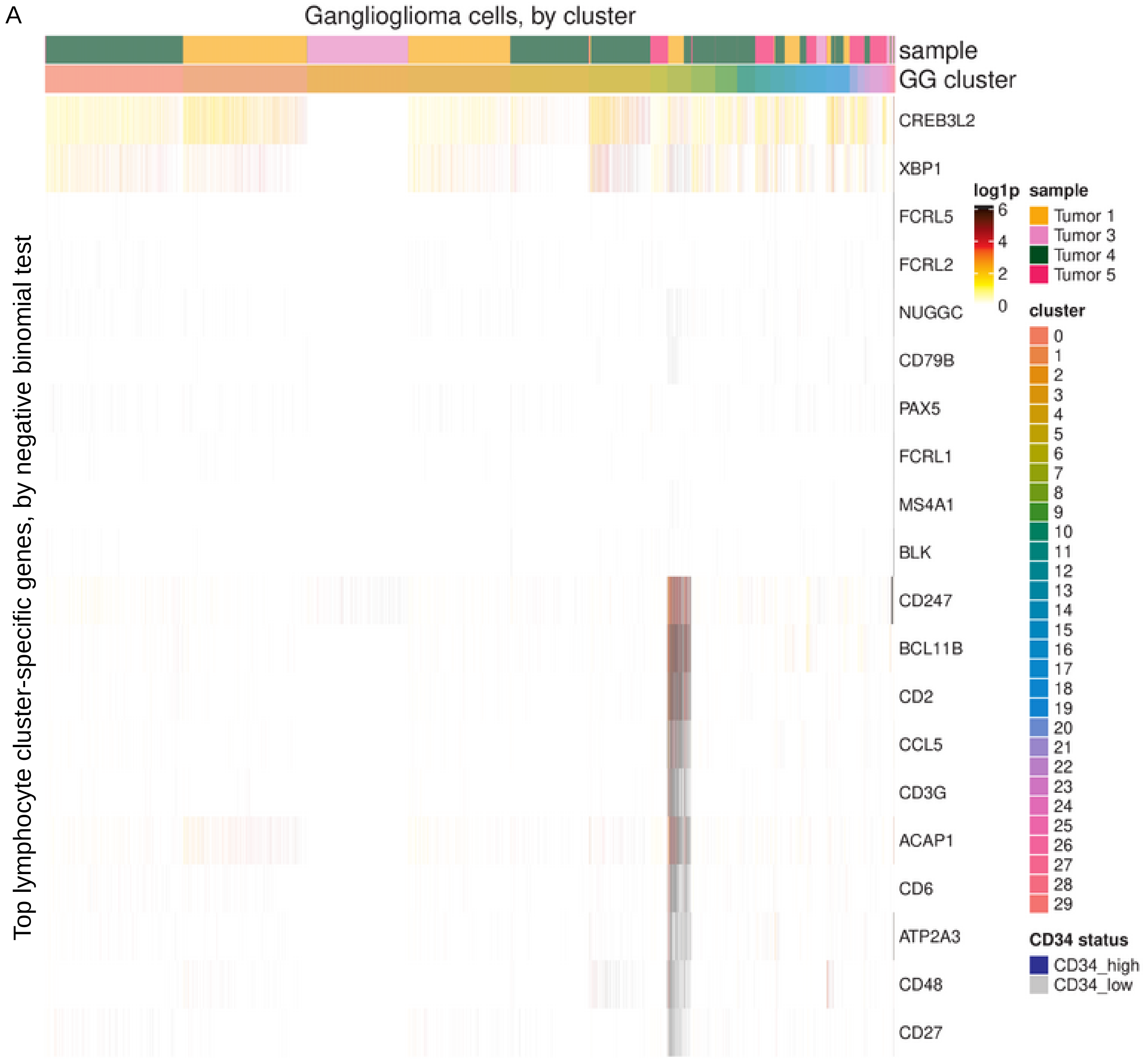
A
Ganglioglioma cells, by cluster
sample
GG cluster
CREB3L2
XBP1
FCRL5
FCRL2
NUGGC
CD79B
PAX5
FCRL1
MS4A1
BLK
CD247
BCL11B
CD2
CCL5
CD3G
ACAP1
CD6
ATP2A3
CD48
CD27
Top lymphocyte cluster-specific genes, by negative binomial test
log1p
6
4
2
0
sample
Tumor 1
Tumor 3
Tumor 4
Tumor 5
cluster
CD34 status
CD34_high
CD34_low

B

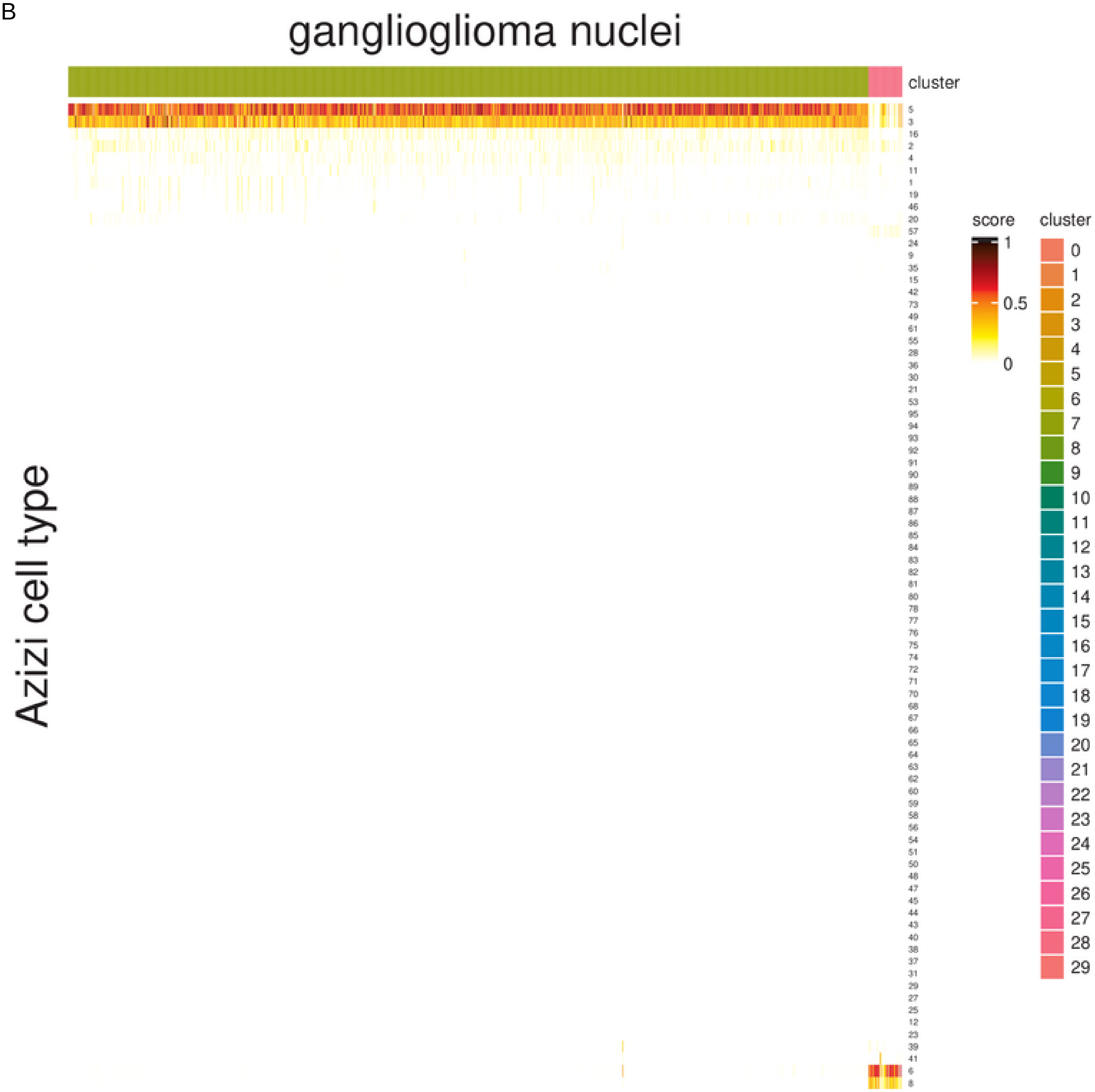
ganglioglioma nuclei
cluster
Azizi cell type
score
1
0.5
0
cluster

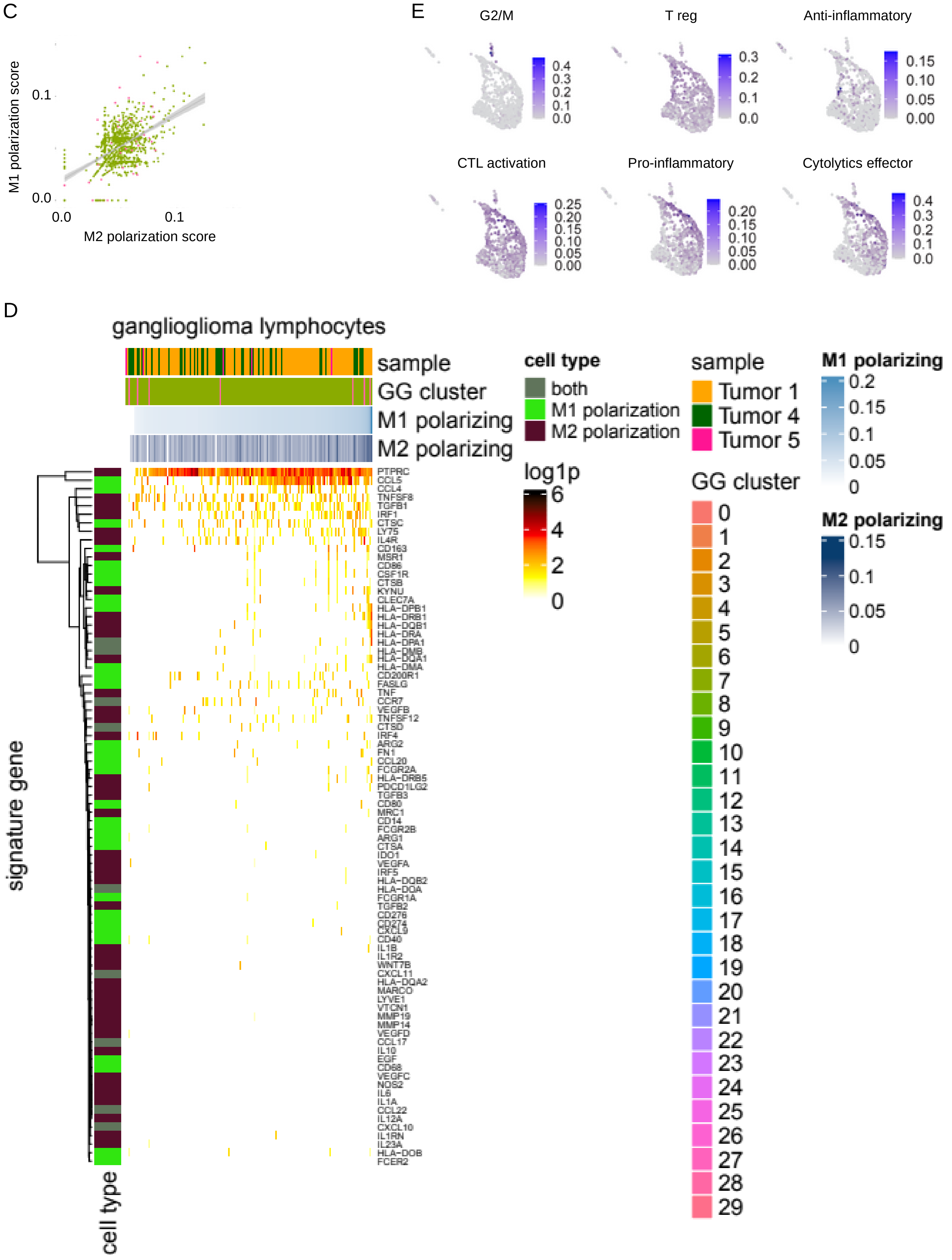
C
M1 polarization score
M2 polarization score
0.0
0.1
E
G2/M
T reg
Anti-inflammatory
CTL activation
Pro-inflammatory
Cytolytics effector
D
ganglioglioma lymphocytes
sample
GG cluster
M1 polarizing
M2 polarizing
signature gene
cell type
PTPRC
CCL5
CCL4
TNFSF8
TGFB1
IRF1
CTSC
LY75
IL4R
CD163
MSR1
CD86
CSF1R
CTSB
KYNU
CLEC7A
HLA-DPB1
HLA-DRB1
HLA-DQB1
HLA-DRA
HLA-DPA1
HLA-DMB
HLA-DQA1
HLA-DMA
CD200R1
FASLG
TNF
CCR7
VEGFB
TNFSF12
CTSD
IRF4
ARG2
FN1
CCL20
FCGR2A
HLA-DRB5
PDCD1LG2
TGFB3
CD80
MRC1
CD14
FCGR2B
ARG1
CTSA
IDO1
VEGFA
IRF5
HLA-DQB2
HLA-DOA
FCGR1A
TGFB2
CD276
CD274
CXCL9
CD40
IL1B
IL1R2
WNT7B
CXCL11
HLA-DQA2
MARCO
LYVE1
VTCN1
MMP19
MMP14
VEGFD
CCL17
IL10
EGF
CD68
VEGFC
NOS2
IL6
IL1A
CCL22
IL12A
CXCL10
IL1RN
IL23A
HLA-DOB
FCER2
cell type
both
M1 polarization
M2 polarization
log1p
sample
Tumor 1
Tumor 4
Tumor 5
GG cluster
M1 polarizing
M2 polarizing

F

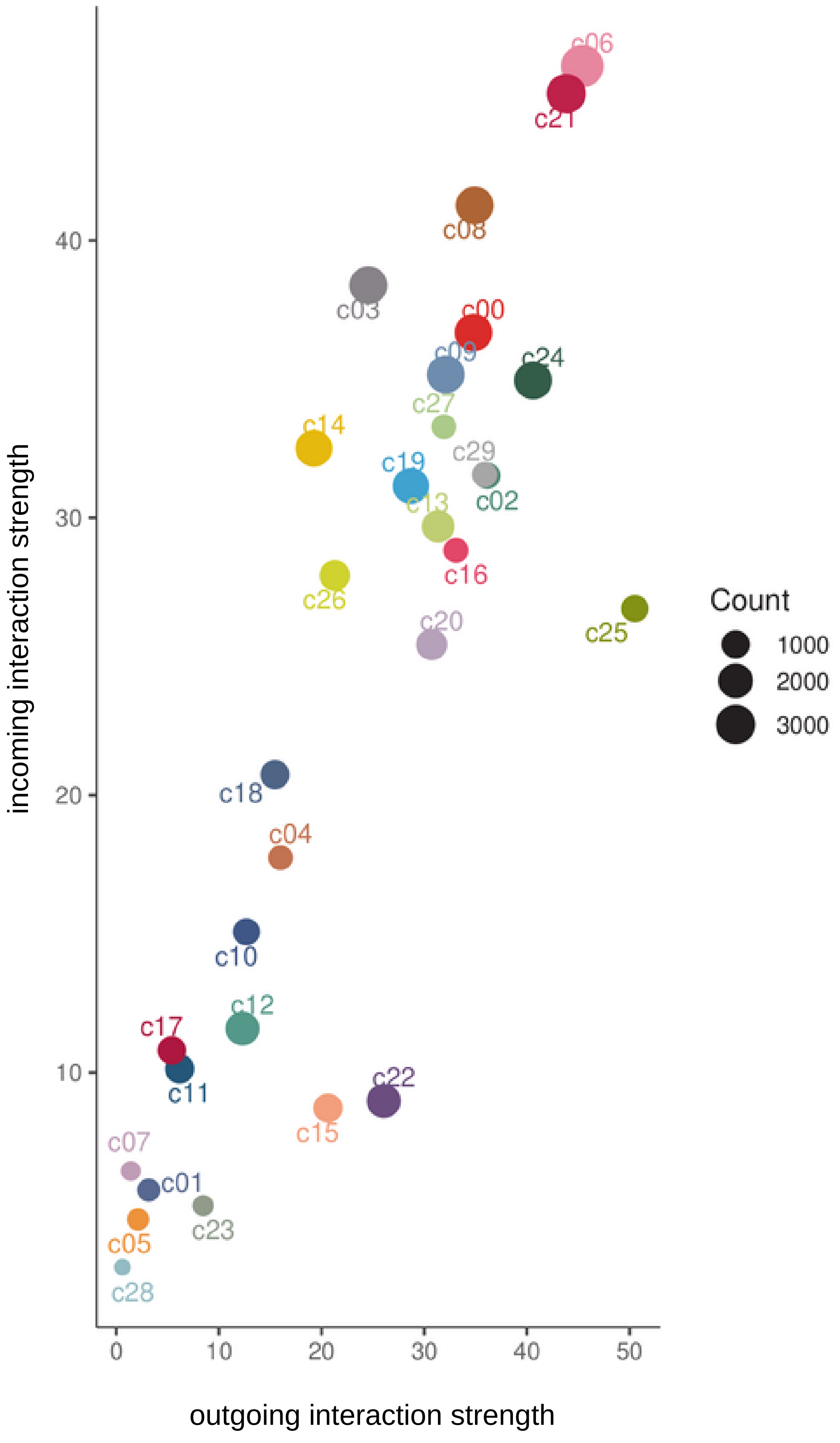

incoming interaction strength
outgoing interaction strength
Count
1000
2000
3000
c00
c01
c02
c03
c04
c05
c06
c07
c08
c09
c10
c11
c12
c13
c14
c15
c16
c17
c18
c19
c20
c21
c22
c23
c24
c25
c26
c27
c28
c29

G

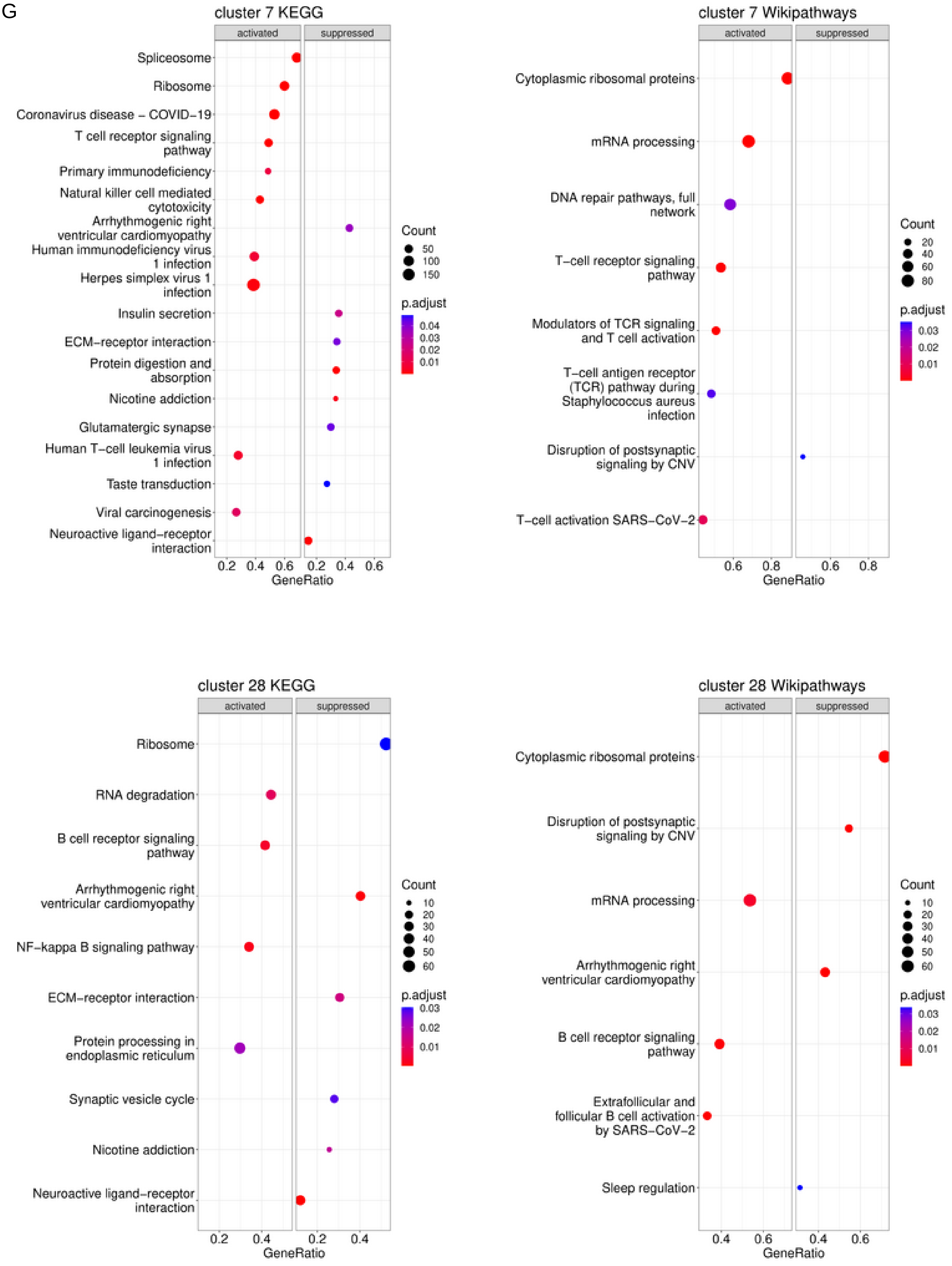
cluster 7 KEGG
activated
suppressed
Spliceosome
Ribosome
Coronavirus disease – COVID−19
T cell receptor signaling pathway
Primary immunodeficiency
Natural killer cell mediated cytotoxicity
Arrhythmogenic right ventricular cardiomyopathy
Human immunodeficiency virus 1 infection
Herpes simplex virus 1 infection
Insulin secretion
ECM−receptor interaction
Protein digestion and absorption
Nicotine addiction
Glutamatergic synapse
Human T−cell leukemia virus 1 infection
Taste transduction
Viral carcinogenesis
Neuroactive ligand−receptor interaction
0.2 0.4 0.6 0.2 0.4 0.6
GeneRatio
Count
50
100
150
p.adjust
0.04
0.03
0.02
0.01
cluster 7 Wikipathways
activated
suppressed
Cytoplasmic ribosomal proteins
mRNA processing
DNA repair pathways, full network
T−cell receptor signaling pathway
Modulators of TCR signaling and T cell activation
T−cell antigen receptor (TCR) pathway during Staphylococcus aureus infection
Disruption of postsynaptic signaling by CNV
T−cell activation SARS−CoV−2
0.6 0.8 0.6 0.8
GeneRatio
Count
20
40
60
80
p.adjust
0.03
0.02
0.01
cluster 28 KEGG
activated
suppressed
Ribosome
RNA degradation
B cell receptor signaling pathway
Arrhythmogenic right ventricular cardiomyopathy
NF−kappa B signaling pathway
ECM−receptor interaction
Protein processing in endoplasmic reticulum
Synaptic vesicle cycle
Nicotine addiction
Neuroactive ligand−receptor interaction
0.2 0.4 0.2 0.4
GeneRatio
Count
10
20
30
40
50
60
p.adjust
0.03
0.02
0.01
cluster 28 Wikipathways
activated
suppressed
Cytoplasmic ribosomal proteins
Disruption of postsynaptic signaling by CNV
mRNA processing
Arrhythmogenic right ventricular cardiomyopathy
B cell receptor signaling pathway
Extrafollicular and follicular B cell activation by SARS−CoV−2
Sleep regulation
0.4 0.6 0.4 0.6
GeneRatio
Count
10
20
30
40
50
60
p.adjust
0.03
0.02
0.01

H

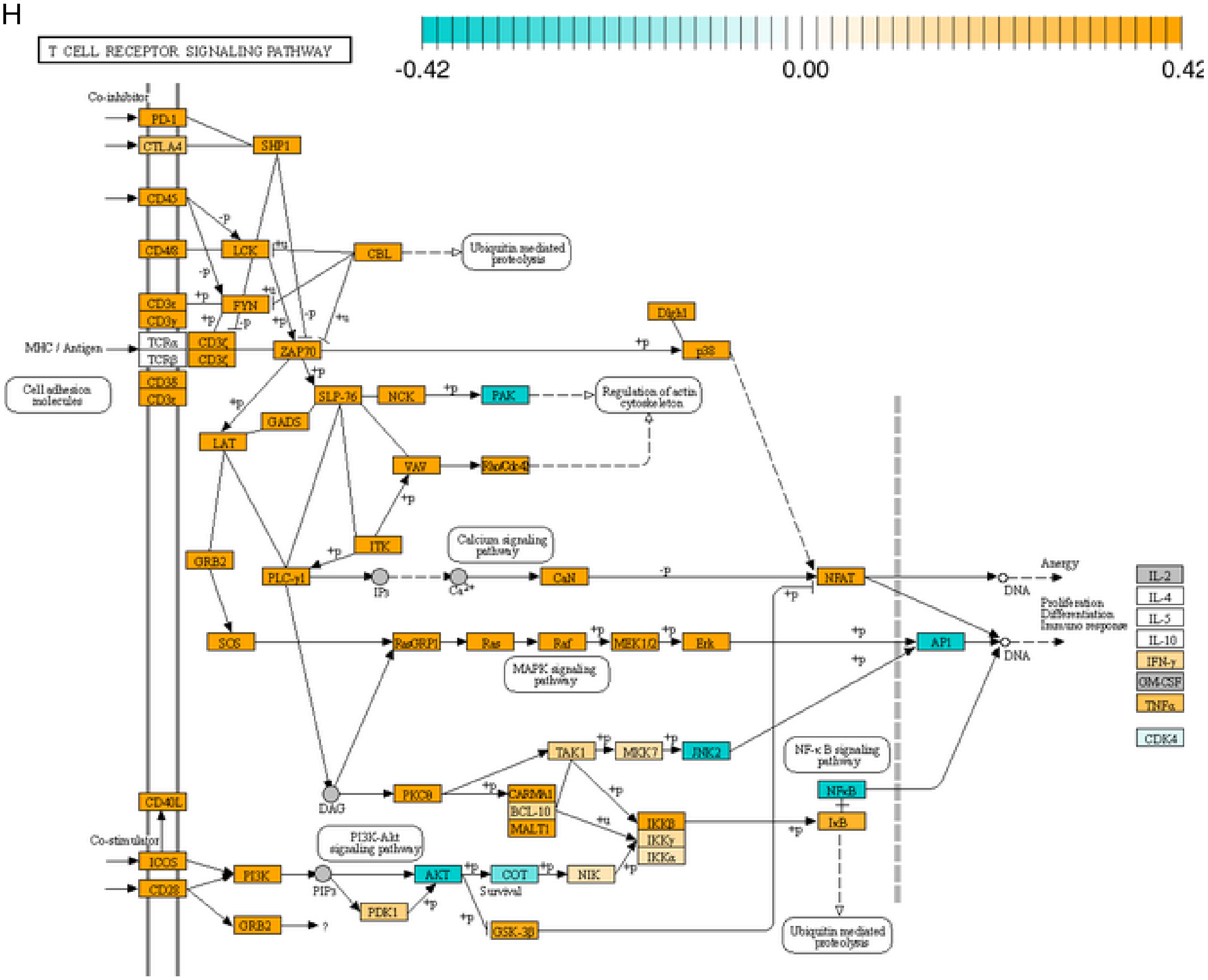
T CELL RECEPTOR SIGNALING PATHWAY
-0.42
0.00
0.42
Co-inhibitor
PD-1
CTLA4
SHP1
CD45
CD4/8
LCK
CBL
Ubiquitin mediated proteolysis
CD3ε
CD3γ
FYN
Dlgh1
MHC / Antigen
TCRα
TCRβ
CD3ζ
ZAP70
p38
Cell adhesion molecules
CD3δ
CD3ε
SLP-76
NCK
PAK
Regulation of actin cytoskeleton
GADS
LAT
VAV
Rho/Cdc42
ITK
Calcium signaling pathway
GRB2
PLC-γ1
IP3
Ca2+
CaN
NFAT
Anergy
DNA
IL-2
IL-4
IL-5
IL-10
IFN-γ
GM-CSF
TNFα
CDK4
Proliferation
Differentiation
Immune response
SOS
RasGRP1
Ras
Raf
MEK1/2
Erk
AP1
MAPK signaling pathway
TAK1
MKK7
JNK2
NF-κB signaling pathway
DAG
PKCθ
CARMA1
BCL-10
MALT1
IKKβ
IKKγ
IKKα
NFκB
IκB
CD40L
Co-stimulator
ICOS
CD28
PI3K
PIP3
PI3K-Akt signaling pathway
AKT
COT
NIK
Survival
PDK1
GRB2
GSK-3β

I

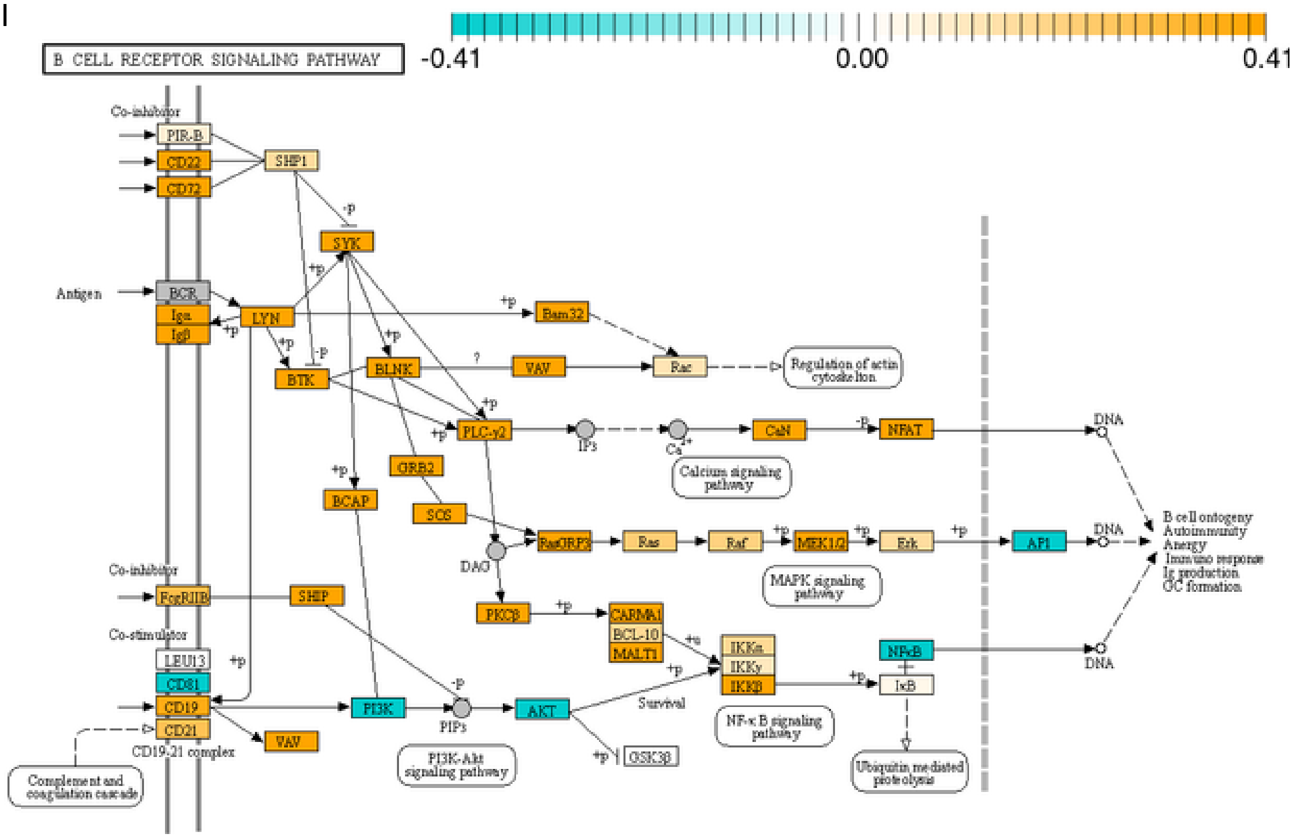
B CELL RECEPTOR SIGNALING PATHWAY
-0.41
0.00
0.41
Co-inhibitor
PIR-B
CD22
CD72
SHP1
SYK
Antigen
BCR
Igα
Igβ
LYN
Bam32
BTK
BLNK
VAV
Rac
Regulation of actin cytoskeleton
PLC-γ2
IP3
Ca2+
CaN
NFAT
DNA
GRB2
Calcium signaling pathway
BCAP
SOS
RasGRP3
Ras
Raf
MEK1/2
Erk
AP1
B cell ontogeny
Autoimmunity
Anergy
Immune response
Ig production
GC formation
DAG
MAPK signaling pathway
Co-inhibitor
FcγRIIB
SHIP
PKCβ
CARMA1
BCL-10
MALT1
IKKα
IKKγ
IKKβ
NFκB
IκB
Co-stimulator
LEU13
CD81
CD19
CD21
CD19-21 complex
PI3K
PIP3
AKT
Survival
NF-κB signaling pathway
VAV
GSK3β
Ubiquitin mediated proteolysis
PI3K-Akt signaling pathway
Complement and coagulation cascade

J

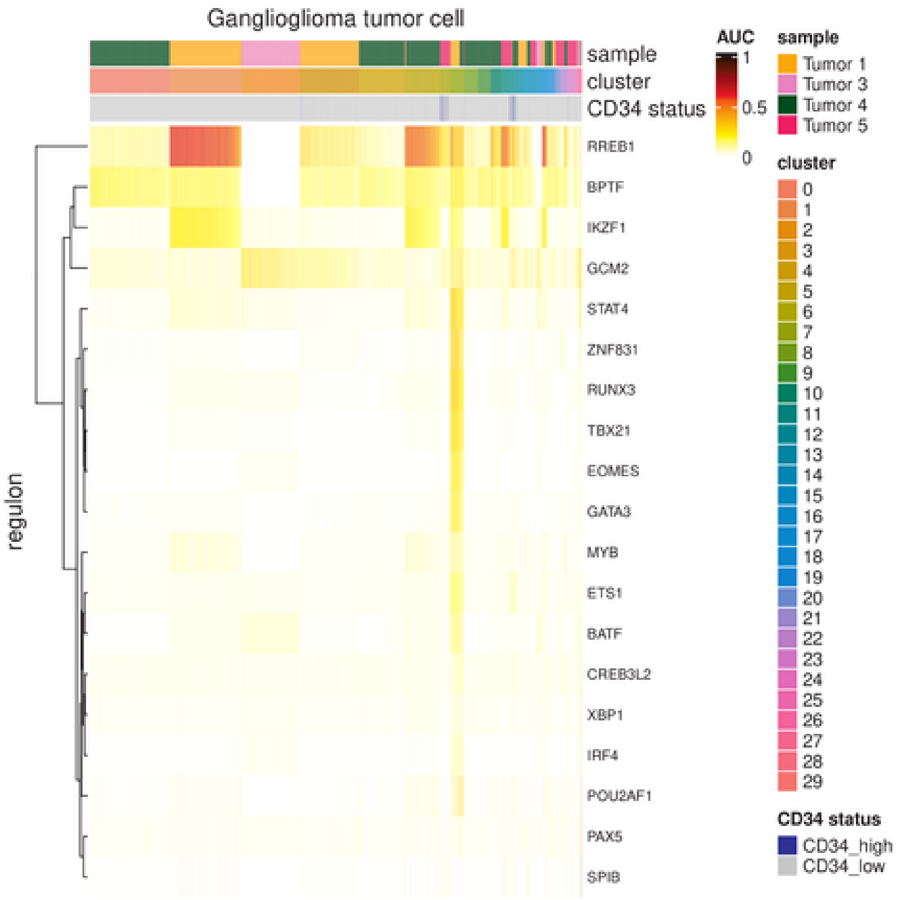

K

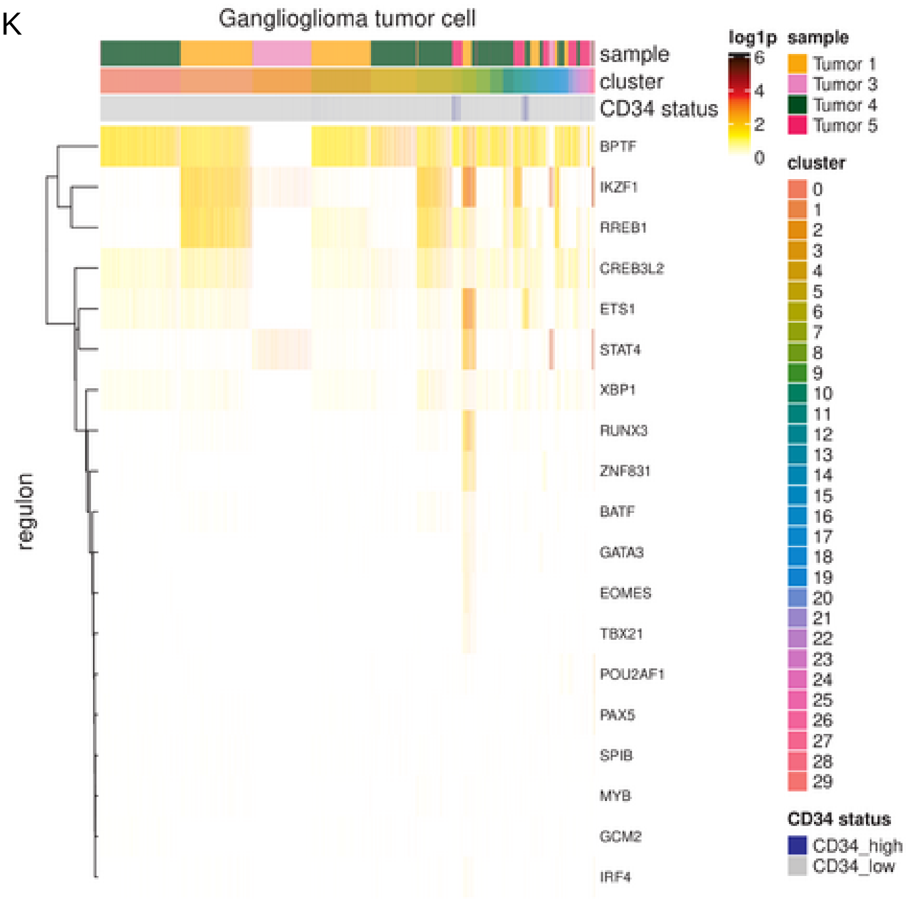

Supplementary Figure 7: Ganglioglioma-infiltrating lymphocytes.
(A) Heatmap of RNA log1p (base 2) for top differentially expressed genes (by negative binomial test) for each of the lymphocyte clusters.
(B) Heatmap of Seurat label based transfer prediction score using Azizi *et al*. immune cell reference atlas. Note Azizi clusters 5 and 3 are T lymphocytes and clusters 6 and 8 B lymphocytes.
(C-E) UCell signature scoring for Azizi *et al*. T cell state signatures.
(C) Scatter plot of lymphocyte scores for M1 polarizing vs M2 polarizing signature.
(D) Heatmap of M1 polarizing and M2 polarizing signature component gene log1p for ganglioglioma lymphocytes.
(E) Feature plots of lymphocyte cells' signature scores identifying dividing cells (G2/M), regulatory T cells (T reg, anti-inflammatory), and cytotoxic T lymphocytes (CTL activation, pro-inflammatory, cytolytics effector).
(F) Scatter plot of CellChat incoming and outgoing interaction strength by ganglioglioma cluster. Note the relatively low interaction of lymphocyte clusters and microglial clusters and the very high interaction of potent neoplastic cluster 6.
(G) Most activated or suppressed KEGG or Wikipathways for lymphocyte clusters.
(H) Pathview depiction of log2-fold-change in the KEGG T cell receptor pathway for cluster 7 versus other ganglioglioma cells.
(I) Pathview depiction of log2-fold-change in the KEGG B cell receptor pathway for cluster 28 versus other ganglioglioma cells.
(J-K) Top lymphocyte cluster transcription factors as nominated by SCENIC, shown in a heatmap with SCENIC regulon AUC (J) or transcription factor transcript log1p (K).

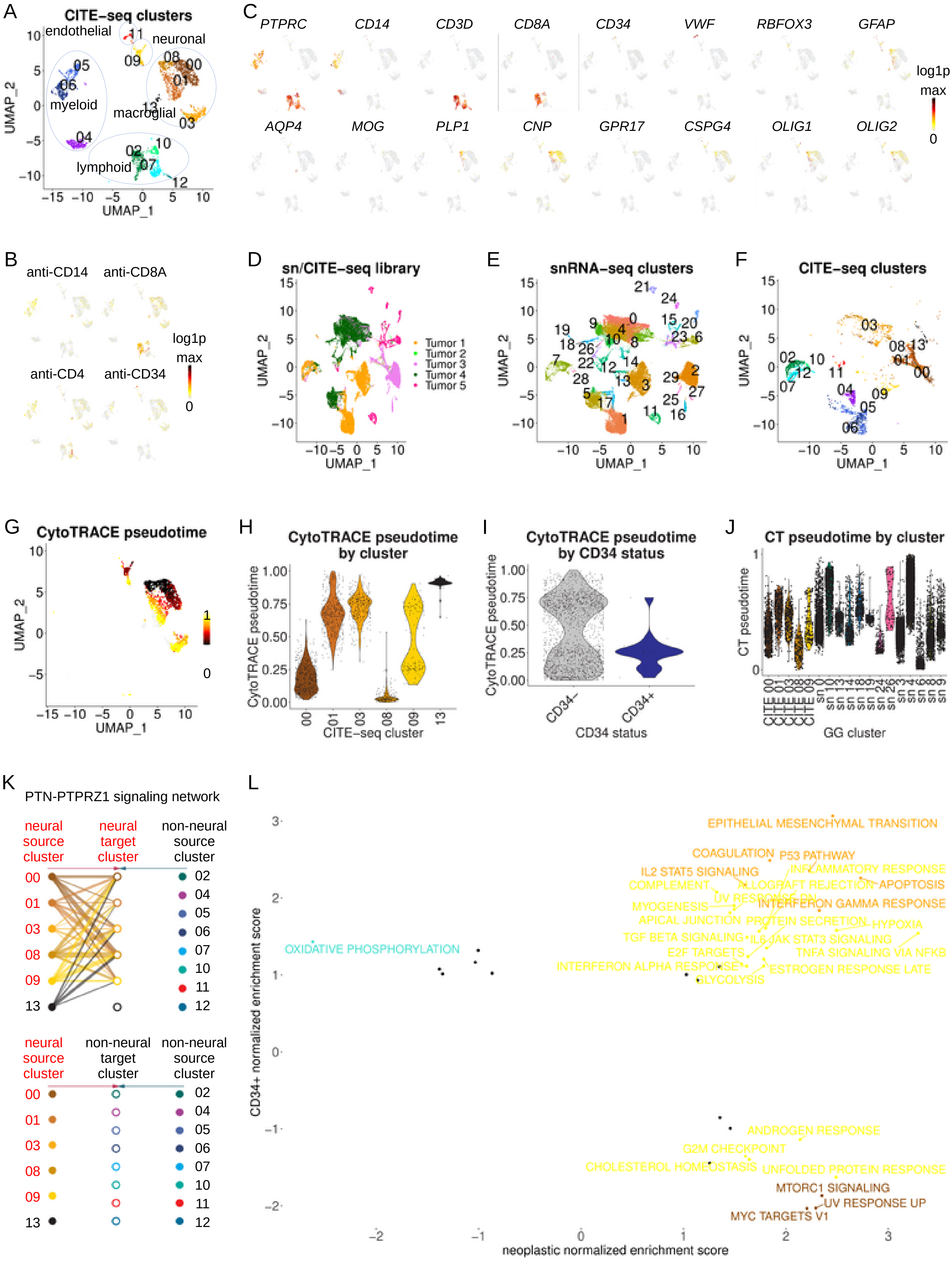
A
CITE-seq clusters
endothelial
neuronal
myeloid
macroglial
lymphoid
UMAP_1
UMAP_2
C
PTPRC CD14 CD3D CD8A CD34 VWF RBFOX3 GFAP
AQP4 MOG PLP1 CNP GPR17 CSPG4 OLIG1 OLIG2
log1p
max
0
B
anti-CD14 anti-CD8A
anti-CD4 anti-CD34
log1p
max
0
D
sn/CITE-seq library
Tumor 1
Tumor 2
Tumor 3
Tumor 4
Tumor 5
E
snRNA-seq clusters
F
CITE-seq clusters
G
CytoTRACE pseudotime
H
CytoTRACE pseudotime by cluster
CytoTRACE pseudotime
CITE-seq cluster
I
CytoTRACE pseudotime by CD34 status
CD34-
CD34+
CD34 status
J
CT pseudotime by cluster
CT pseudotime
GG cluster
K
PTN-PTPRZ1 signaling network
neural source cluster
neural target cluster
non-neural source cluster
neural source cluster
non-neural target cluster
non-neural source cluster
L
EPITHELIAL MESENCHYMAL TRANSITION
COAGULATION
P53 PATHWAY
IL2 STAT5 SIGNALING
INFLAMMATORY RESPONSE
COMPLEMENT
ALLOGRAFT REJECTION
APOPTOSIS
UV RESPONSE DN
MYOGENESIS
INTERFERON GAMMA RESPONSE
APICAL JUNCTION
PROTEIN SECRETION
HYPOXIA
TGF BETA SIGNALING
IL6 JAK STAT3 SIGNALING
OXIDATIVE PHOSPHORYLATION
E2F TARGETS
TNFA SIGNALING VIA NFKB
INTERFERON ALPHA RESPONSE
ESTROGEN RESPONSE LATE
GLYCOLYSIS
ANDROGEN RESPONSE
G2M CHECKPOINT
CHOLESTEROL HOMEOSTASIS
UNFOLDED PROTEIN RESPONSE
MTORC1 SIGNALING
UV RESPONSE UP
MYC TARGETS V1
CD34+ normalized enrichment score
neoplastic normalized enrichment score

M

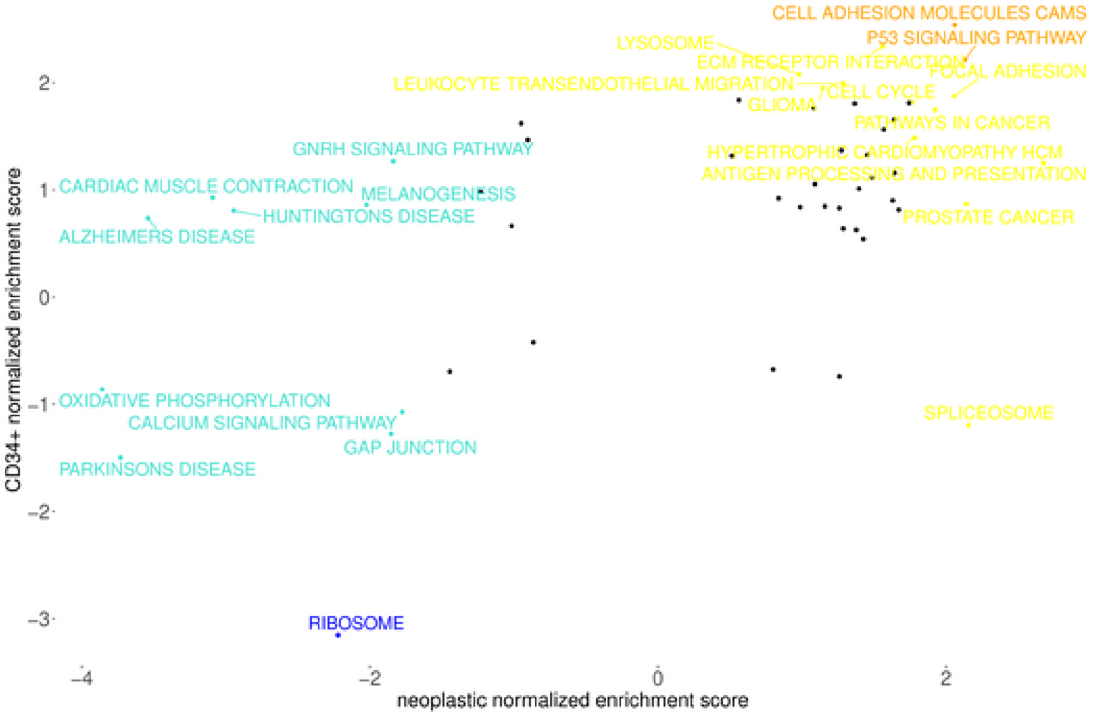
CELL ADHESION MOLECULES CAMS
P53 SIGNALING PATHWAY
LYSOSOME
ECM RECEPTOR INTERACTION
FOCAL ADHESION
LEUKOCYTE TRANSENDOTHELIAL MIGRATION
CELL CYCLE
GLIOMA
PATHWAYS IN CANCER
GNRH SIGNALING PATHWAY
HYPERTROPHIC CARDIOMYOPATHY HCM
ANTIGEN PROCESSING AND PRESENTATION
CARDIAC MUSCLE CONTRACTION
MELANOGENESIS
HUNTINGTONS DISEASE
PROSTATE CANCER
ALZHEIMERS DISEASE
OXIDATIVE PHOSPHORYLATION
CALCIUM SIGNALING PATHWAY
SPLICEOSOME
GAP JUNCTION
PARKINSONS DISEASE
RIBOSOME
CD34+ normalized enrichment score
neoplastic normalized enrichment score
2
1
0
−1
−2
−3
−4
−2
0
2

N

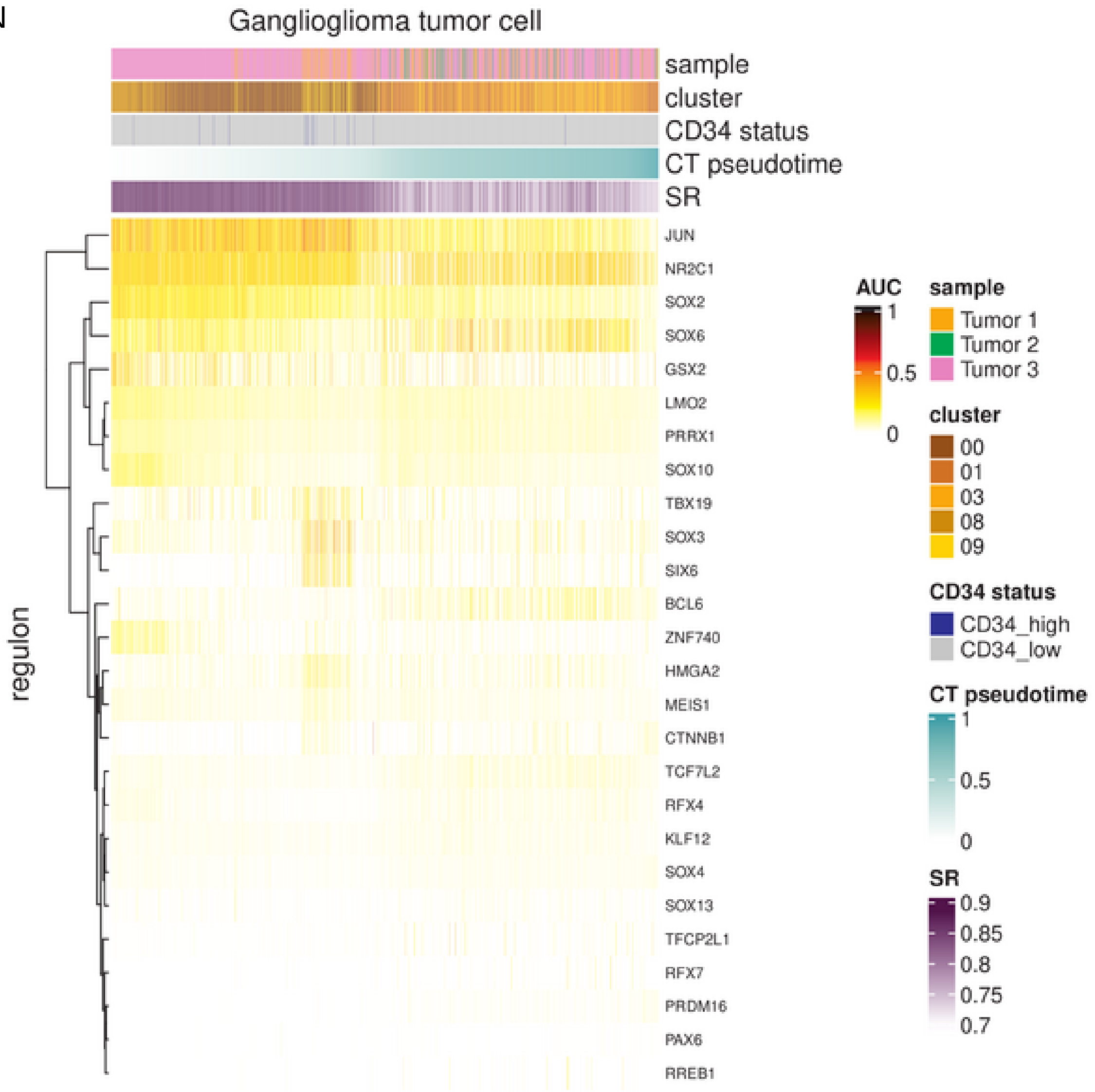
Ganglioglioma tumor cell
sample
cluster
CD34 status
CT pseudotime
SR
JUN
NR2C1
SOX2
SOX6
GSX2
LMO2
PRRX1
SOX10
TBX19
SOX3
SIX6
BCL6
ZNF740
HMGA2
MEIS1
CTNNB1
TCF7L2
RFX4
KLF12
SOX4
SOX13
TFCP2L1
RFX7
PRDM16
PAX6
RREB1
regulon
AUC
1
0.5
0
sample
Tumor 1
Tumor 2
Tumor 3
cluster
00
01
03
08
09
CD34 status
CD34_high
CD34_low
CT pseudotime
1
0.5
0
SR
0.9
0.85
0.8
0.75
0.7

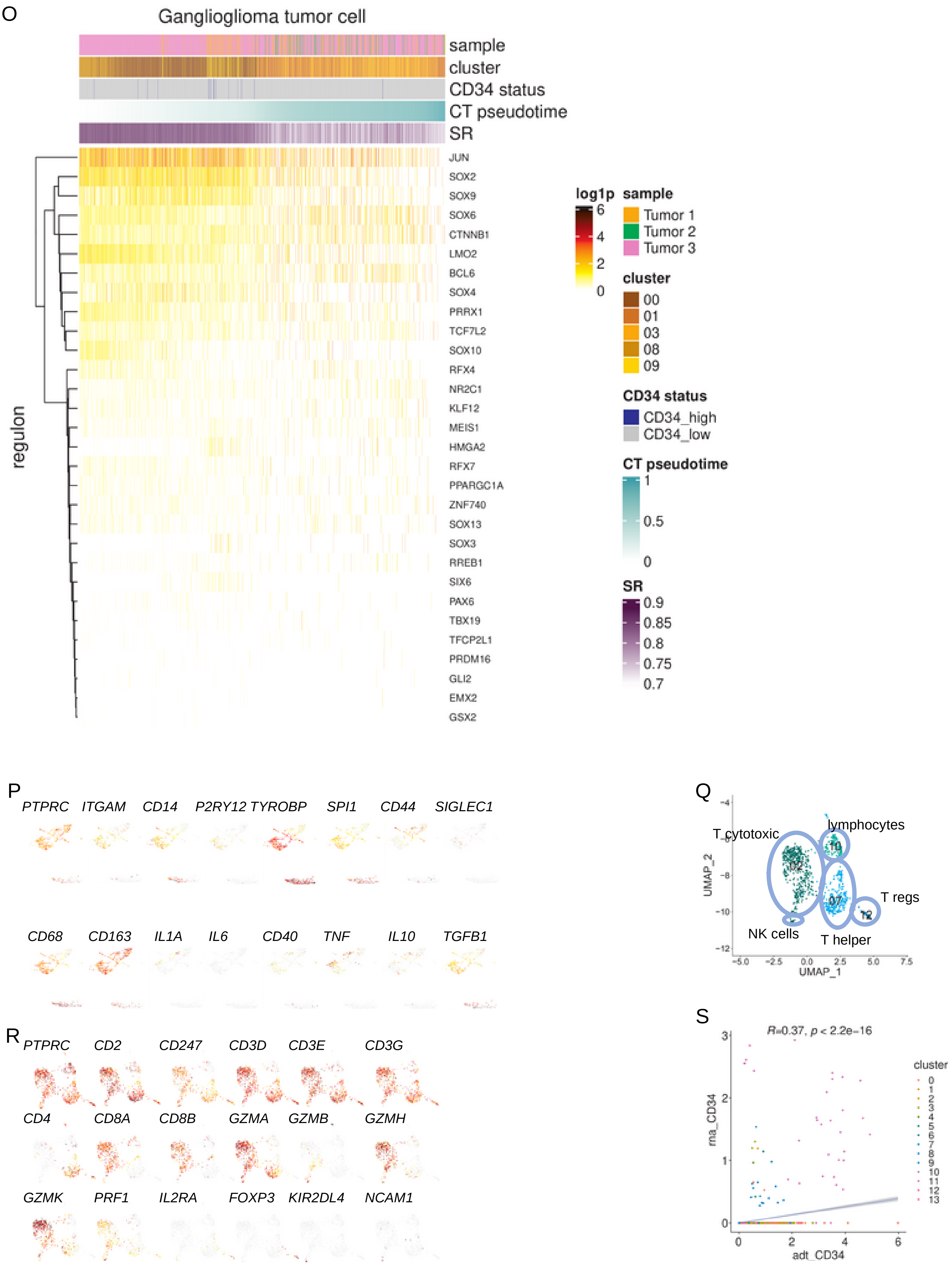
O
Ganglioglioma tumor cell
sample
cluster
CD34 status
CT pseudotime
SR
regulon
JUN
SOX2
SOX9
SOX6
CTNNB1
LMO2
BCL6
SOX4
PRRX1
TCF7L2
SOX10
RFX4
NR2C1
KLF12
MEIS1
HMGA2
RFX7
PPARGC1A
ZNF740
SOX13
SOX3
RREB1
SIX6
PAX6
TBX19
TFCP2L1
PRDM16
GLI2
EMX2
GSX2
log1p
6
4
2
0
sample
Tumor 1
Tumor 2
Tumor 3
cluster
00
01
03
08
09
CD34 status
CD34_high
CD34_low
CT pseudotime
1
0.5
0
SR
0.9
0.85
0.8
0.75
0.7
P
PTPRC ITGAM CD14 P2RY12 TYROBP SPI1 CD44 SIGLEC1
CD68 CD163 IL1A IL6 CD40 TNF IL10 TGFB1
Q
T cytotoxic
lymphocytes
T regs
NK cells
T helper
UMAP_2
UMAP_1
R
PTPRC CD2 CD247 CD3D CD3E CD3G
CD4 CD8A CD8B GZMA GZMB GZMH
GZMK PRF1 IL2RA FOXP3 KIR2DL4 NCAM1
S
R=0.37, p < 2.2e-16
rna_CD34
adt_CD34
cluster
0
1
2
3
4
5
6
7
8
9
10
11
12
13

Supplementary Figure 8: Ganglioglioma validation by CITE-seq.
(A) UMAP plot colored by nearest neighbor clusters (numbered 00-13) with broad cell type, supercluster annotations superimposed.
(B) Feature plots of CITE-seq epitope probing to identify ganglioglioma immune cells and CD34-positive cells.
(C) Feature plots of CITE-seq immune, endothelial, neuronal, astrocyte, oligodendrocyte, and oligodendrocyte precursor cell markers to distinguish major ganglioglioma cell types.
(D-F) Dim plots of results of ganglioglioma CITE-seq integration with snRNA-seq data using Seurat CCA v4. (D) Combined data, colored by tumor. (E) SnRNA-seq, colored and labeled by cluster. (F) CITE-seq, colored and labeled by cluster.
(G-J) Ganglioglioma neural(-like) cellular hierarchy by CITE-seq. (G) Feature plot of ganglioglioma CITE-seq neural(-like) cell CellRank/CytoTRACE pseudotime.
(H-I) Violin plots of CytoTRACE pseudotime by ganglioglioma CITE-seq cluster (H) or *CD34* status (I).
(J) Violin plots of CytoTRACE pseudotime by ganglioglioma CITE-seq (CITE) or snRNA-seq (sn) cluster, as calculated after integration of the datasets by Seurat CCA as above.
(K) Ganglioglioma CITE-seq neural cell communication via PTN (ligand) interactions with/antagonism of PTPRZ1 (receptor) as nominated by CellChat. Hierarchy plots of significant PTN-PTPRZ1 interaction shown for ganglioglioma neural cell targets (left) and non-neural cell targets (right). Line thickness reflects strength of observed interaction. Neoplastic clusters noted in red; stromal clusters noted in black.
(L) Plot of Hallmark pathway normalized enrichment score (NES) when comparing neoplastic CITE-seq *CD34*+ and *CD34*- cells vs when comparing neoplastic-appearing vs normal neural-appearing cells. Pathways with NES>1 and FDR q-value<0.05 in both comparisons shown in orange. Pathways with NES<-1 and q-value<0.05 in both comparisons shown in dark blue. Pathways with |NES|>1 but in opposing directions with q-value<0.05 in both comparisons shown in chocolate. Pathways with |NES|>1 and q-value<0.05 in only one comparison shown in yellow or turquoise. Pathways otherwise shown in black or (mostly) not shown.
(M) Analogous to panel E but showing KEGG pathways.
(N-O) Heatmaps of SCENIC AUC scores (N) and corresponding transcription factor concentrations (O) for neoplastic CITE-seq cells and transcription factors of interest due to SCENIC nomination among potent neoplastic cells and/or *CD34*+ neoplastic snRNA-seq or CITE-seq cells. Note cells (columns) are ordered by CytoTRACE pseudotime.
(P) Feature plots of CITE-seq myeloid cell expression of myeloid markers of *a priori* interest.
(Q) UMAP with CITE-seq lymphocyte cell typing results. Cells colored by cluster.
(R) Feature plots of CITE-seq lymphocyte expression of lymphocyte markers of *a priori* interest.
(S) Scatterplot of CITE-seq *CD34* RNA concentration versus CITE-seq anti-CD34 concentration showing correlation thereof.

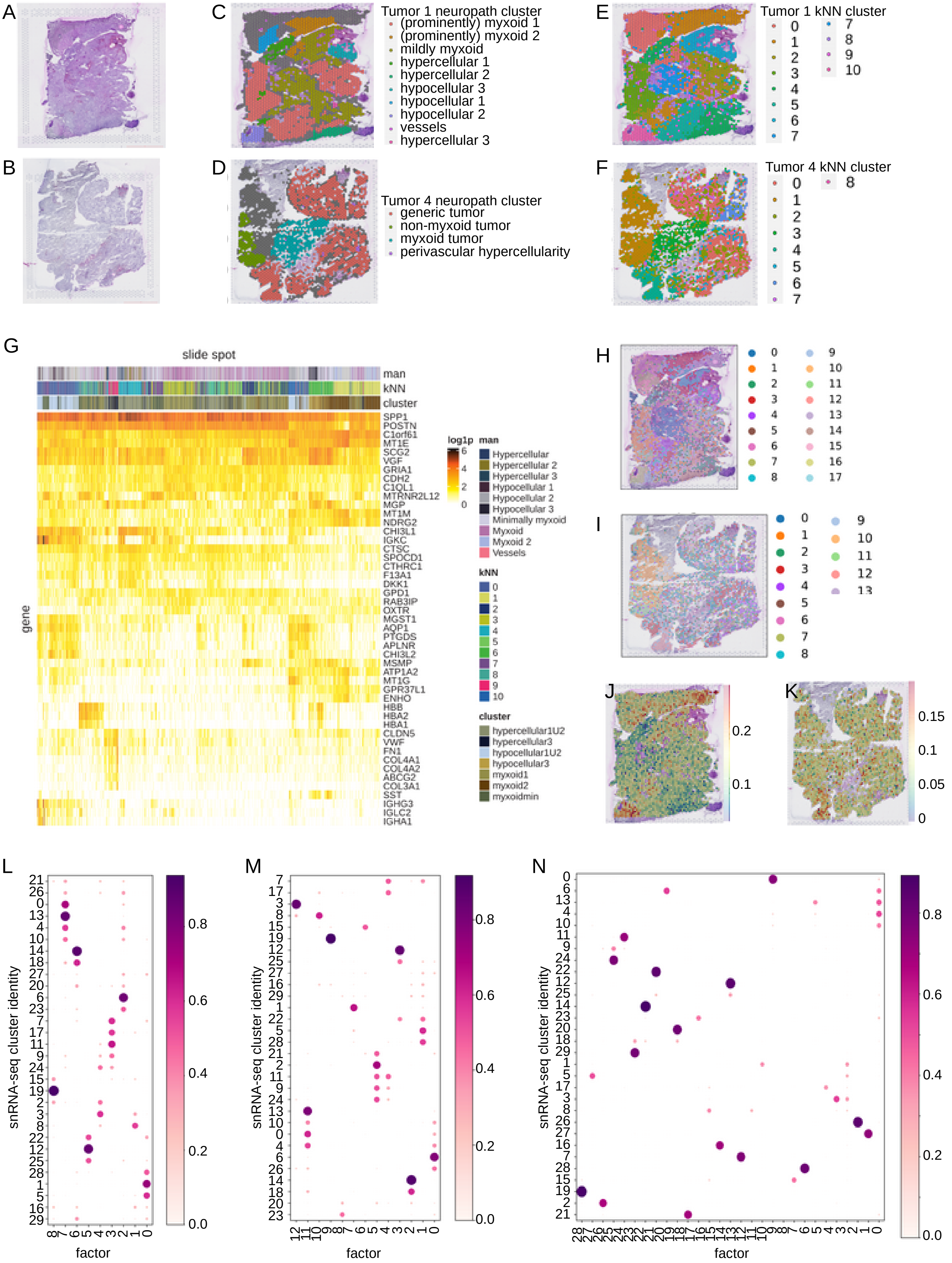
A
B
C
Tumor 1 neuropath cluster
(prominently) myxoid 1
(prominently) myxoid 2
mildly myxoid
hypercellular 1
hypercellular 2
hypocellular 3
hypocellular 1
hypocellular 2
vessels
hypercellular 3
D
Tumor 4 neuropath cluster
generic tumor
non-myxoid tumor
myxoid tumor
perivascular hypercellularity
E
Tumor 1 kNN cluster
0 1 2 3 4 5 6 7 7 8 9 10
F
Tumor 4 kNN cluster
0 1 2 3 4 5 6 7 8
G
slide spot
man
kNN
cluster
gene
SPP1
POSTN
C1orf61
MT1E
SCG2
VGF
GRIA1
CDH2
C1QL1
MTRNR2L12
MGP
MT1M
NDRG2
CHI3L1
IGKC
CTSC
SPOCD1
CTHRC1
F13A1
DKK1
GPD1
RAB3IP
OXTR
MGST1
AQP1
PTGDS
APLNR
CHI3L2
MSMP
ATP1A2
MT1G
GPR37L1
ENHO
HBB
HBA2
HBA1
CLDN5
VWF
FN1
COL4A1
COL4A2
ABCG2
COL3A1
SST
IGHG3
IGLC2
IGHA1
log1p
6 4 2 0
man
Hypercellular
Hypercellular 2
Hypercellular 3
Hypocellular 1
Hypocellular 2
Hypocellular 3
Minimally myxoid
Myxoid
Myxoid 2
Vessels
kNN
0 1 2 3 4 5 6 7 8 9 10
cluster
hypercellular1U2
hypercellular3
hypocellular1U2
hypocellular3
myxoid1
myxoid2
myxoidmin
H
0 1 2 3 4 5 6 7 8 9 10 11 12 13 14 15 16 17
I
0 1 2 3 4 5 6 7 8 9 10 11 12 13
J
0.2
0.1
K
0.15
0.1
0.05
0
L
snRNA-seq cluster identity
21 26 0 13 4 10 14 18 27 20 6 23 7 17 11 9 24 15 19 2 3 8 22 12 25 28 1 5 16 29
8 7 6 5 4 3 2 1 0
factor
0.8 0.6 0.4 0.2 0.0
M
snRNA-seq cluster identity
7 17 3 8 15 19 12 25 27 16 29 1 22 5 28 21 2 11 9 24 13 10 0 4 6 26 14 18 20 23
12 11 10 9 8 7 6 5 4 3 2 1 0
factor
0.8 0.6 0.4 0.2 0.0
N
snRNA-seq cluster identity
0 6 13 4 10 11 9 24 22 12 25 14 23 20 18 29 1 5 17 3 8 26 27 16 7 28 15 19 2 21
29 28 27 26 25 24 23 22 21 20 19 18 17 16 15 14 13 12 11 10 9 8 7 6 5 4 3 2 1 0
factor
0.8 0.6 0.4 0.2 0.0

Supplementary Figure 9: Ganglioglioma spatial transcriptomics shown for two samples from tumor 1 and tumor 4.
(A-B) H&E stained slide for tumor 1 (A) and tumor 4 (B).
(C-D) Spatial feature plots of neuropathologist (GYL) annotations for tumor 1 (C) and tumor 4 (D).
(E-F) Spatial feature plots of kNN clusters for tumor 1 (E) and tumor 4 (F).
(G) Heatmap of expression for the top (by fold-change) spatial cluster differentially expressed transcripts for tumor 1.
(H-I) Cell2location tissue regions derived from cell abundance kNN for tumor 1 (H) and tumor 4 (I).
(J) SnRNA-seq cluster top 30 differentially expressed genes (by Wilcox) were used to signature score each spatial transcriptomics spot (UCell). Shown is cluster 6 neuroectoderm neural precursor-like cell cluster scoring for tumor 1, showing two areas of particular interest for these cells.
(K) Analogous to (J) but for tumor 4.
(L-N) Cell type colocalization by cell2location non-negative matrix factorization with 9 (L), 13 (M), and 29 (N) factors shown for tumor 1 spots deconvoluted using cell2location with ganglioglioma snRNA-seq clusters as reference.

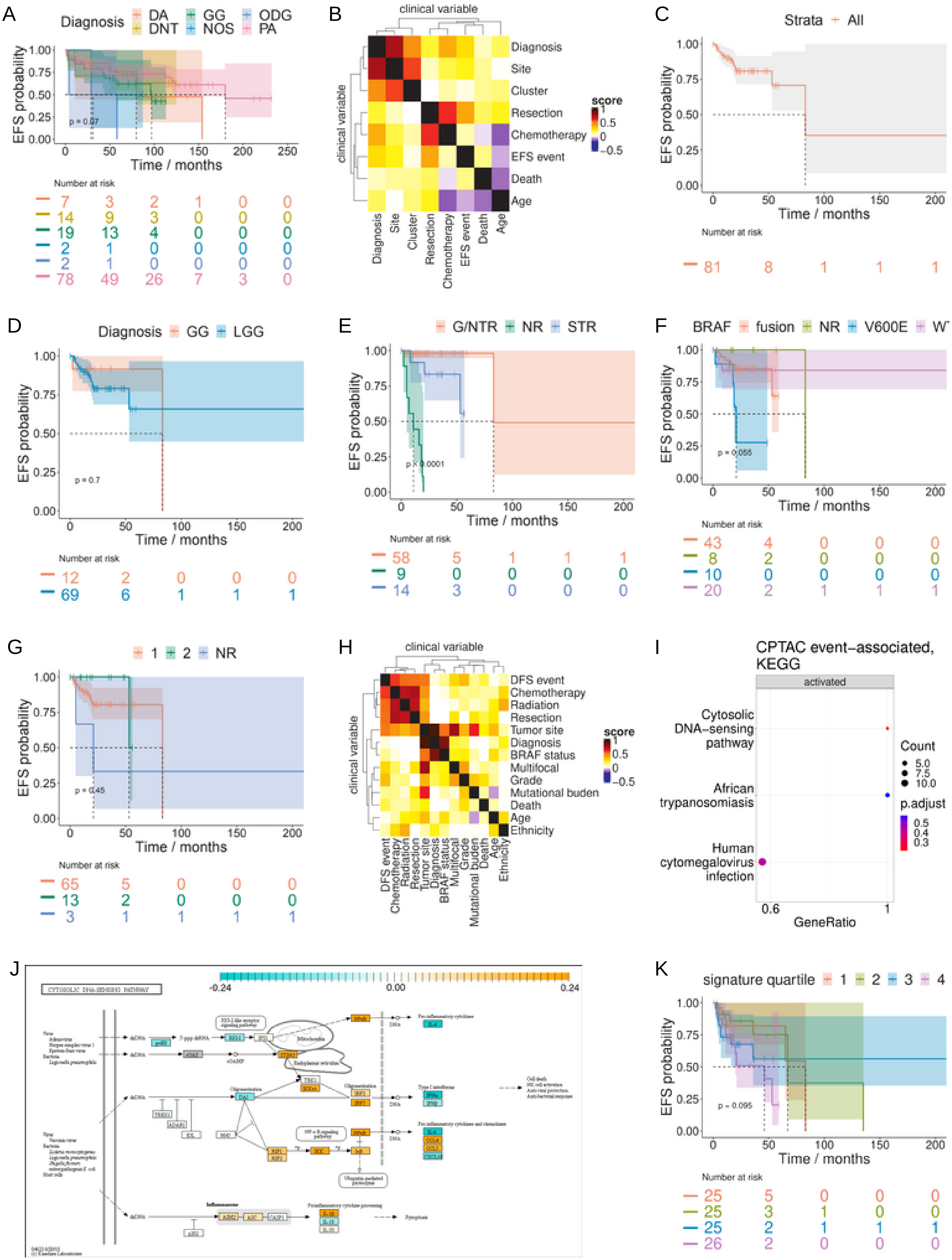
A
Diagnosis DA DNT GG NOS ODG PA
EFS probability
p = 0.07
Time / months
Number at risk
7 3 2 1 0 0
14 9 3 0 0 0
19 13 4 0 0 0
2 1 0 0 0 0
2 1 0 0 0 0
78 49 26 7 3 0
B
clinical variable
Diagnosis
Site
Cluster
Resection
Chemotherapy
EFS event
Death
Age
score
C
Strata All
EFS probability
Time / months
Number at risk
81 8 1 1 1
D
Diagnosis GG LGG
EFS probability
p = 0.7
Time / months
Number at risk
12 2 0 0 0
69 6 1 1 1
E
G/NTR NR STR
EFS probability
p < 0.0001
Time / months
Number at risk
58 5 1 1 1
9 0 0 0 0
14 3 0 0 0
F
BRAF fusion NR V600E WT
EFS probability
p = 0.055
Time / months
Number at risk
43 4 0 0 0
8 2 0 0 0
10 0 0 0 0
20 2 1 1 1
G
1 2 NR
EFS probability
p = 0.45
Time / months
Number at risk
65 5 0 0 0
13 2 0 0 0
3 1 1 1 1
H
clinical variable
DFS event
Chemotherapy
Radiation
Resection
Tumor site
Diagnosis
BRAF status
Multifocal
Grade
Mutational buden
Death
Age
Ethnicity
score
I
CPTAC event-associated, KEGG
activated
Cytosolic DNA-sensing pathway
African trypanosomiasis
Human cytomegalovirus infection
Count
p.adjust
GeneRatio
J
CYTOSOLIC DNA-SENSING PATHWAY
-0.24
0.00
0.24
K
signature quartile 1 2 3 4
EFS probability
p = 0.095
Time / months
Number at risk
25 5 0 0 0
25 3 1 0 0
25 2 1 1 1
26 2 0 0 0

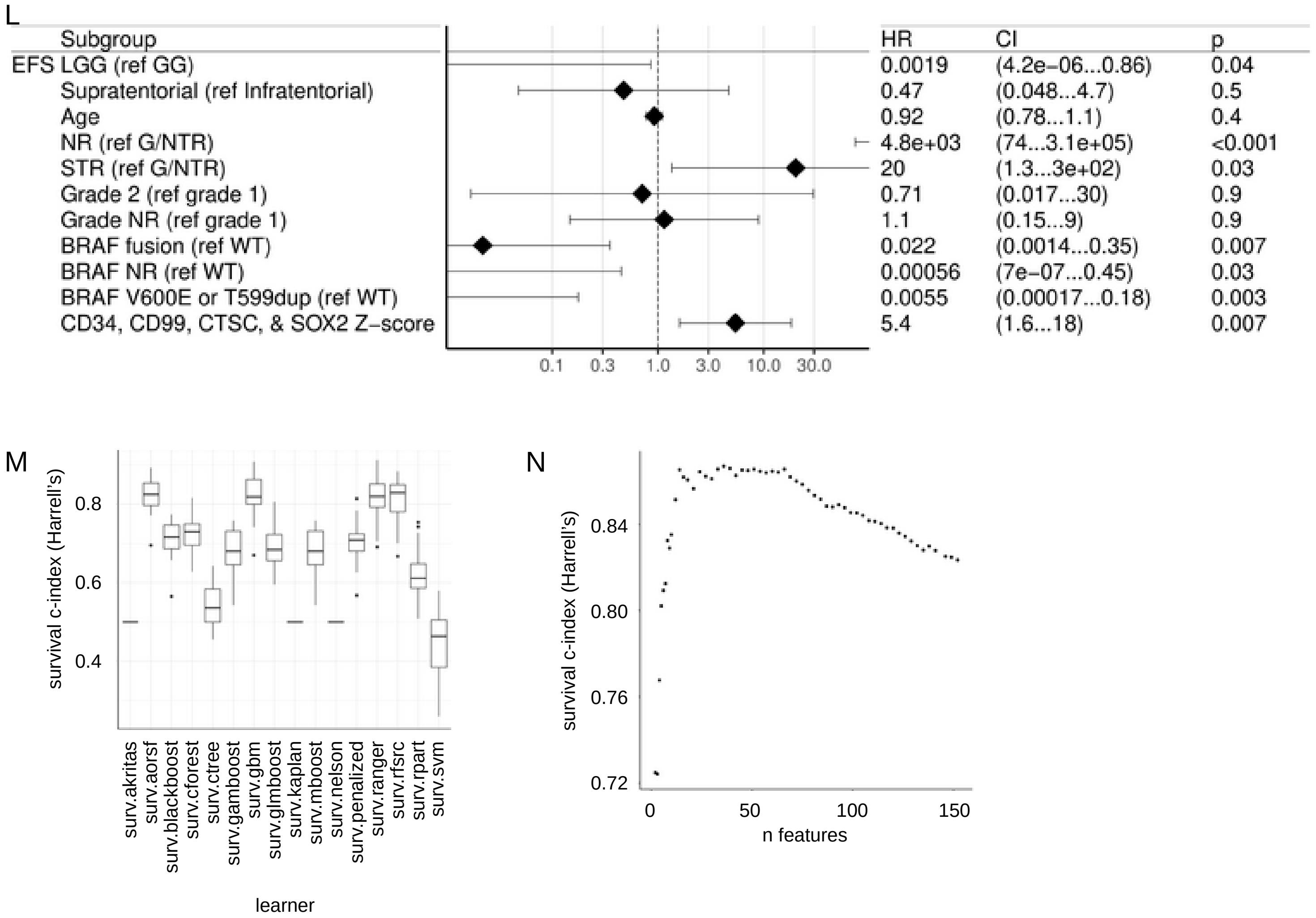
L
Subgroup
EFS LGG (ref GG)
Supratentorial (ref Infratentorial)
Age
NR (ref G/NTR)
STR (ref G/NTR)
Grade 2 (ref grade 1)
Grade NR (ref grade 1)
BRAF fusion (ref WT)
BRAF NR (ref WT)
BRAF V600E or T599dup (ref WT)
CD34, CD99, CTSC, & SOX2 Z−score
0.1 0.3 1.0 3.0 10.0 30.0
HR
0.0019
0.47
0.92
4.8e+03
20
0.71
1.1
0.022
0.00056
0.0055
5.4
CI
(4.2e−06...0.86)
(0.048...4.7)
(0.78...1.1)
(74...3.1e+05)
(1.3...3e+02)
(0.017...30)
(0.15...9)
(0.0014...0.35)
(7e−07...0.45)
(0.00017...0.18)
(1.6...18)
p
0.04
0.5
0.4
<0.001
0.03
0.9
0.9
0.007
0.03
0.003
0.007
M
survival c-index (Harrell's)
0.4
0.6
0.8
surv.akritas
surv.aorsf
surv.blackboost
surv.cforest
surv.ctree
surv.gamboost
surv.gbm
surv.glmboost
surv.kaplan
surv.mboost
surv.nelson
surv.penalized
surv.ranger
surv.rfsrc
surv.rpart
surv.svm
learner
N
survival c-index (Harrell's)
0.72
0.76
0.80
0.84
0 50 100 150
n features

Supplementary Figure 10: Low grade glioma outcomes analysis.
(A) Kaplan-Meier EFS (95% CI shaded, dotted line for median) for Bergthold *et al*. patients by diagnosis.
(B) Correlogram of available Bergthold *et al*. clinical annotations including diagnosis, site (infratentorial versus supratentorial), cluster (as determined by SOM by Bergthold *et al*.), extent of resection (biopsy, subtotal resection, near total resection, and complete resection), receipt of chemotherapy, EFS event, death, and age (continuous). Strength of association is calculated for nominal vs nominal with a bias corrected Cramer's V, numeric vs numeric with Spearman (default), and nominal vs numeric with ANOVA.
(C-G) Kaplan-Meier EFS (95% CI shaded, dotted line for median) for CPTAC patients overall (C), by diagnosis (D), by resection status (E), by BRAF mutational status (F), and by tumor grade (G).
(H) Correlogram for CPTAC patients, analogous to (B) for Bergthold *et al*. patients.
(I) Dotplot of top differentially expressed KEGG pathways in comparing CPTAC event-associated tumors to those with no recorded event.
(J) Pathview representation of cGAS/STING signaling pathway colored by log2-fold-change in RNA concentrations comparing CPTAC event-associated tumors to those with no recorded event.
(K) Kaplan-Meier EFS (95% CI shaded, dotted line for median) for CPTAC patients by UCell score quartile for the gene signature of CD34, SOX2, CD99, and CTSC.
(L) Forest plot of COXPH EFS HR for CPTAC patients with continuous variables kept continuous. Includes HR for Z-score for the gene signature of CD34, SOX2, CD99, and CTSC.
(M) Supervised learning algorithm selection results (Harrell's c-index), training on Bergthold *et al*. data with 2.5% most important features by surv.rfsrc.
(N) Supervised learning feature selection results (Harrell's c-index), training on Bergthold *et al*. data with 2.5% most important features by surv.rfsrc using surv.aorsf as learner.
EFS=event-free survival, NR=none reported, GTR=gross total resection, NTR=near total resection, G/NTR=gross or near total resection, STR=subtotal resection, WT=wild-type. DA=diffuse astrocytoma, GG=ganglioglioma, ODG=oligodendroglioma, DNT=dysplastic neuroepithelial tumor, NOS or LGG=low-grade glioma, not otherwise specified. PA=pilocytic astrocytoma.
