## Supplementary information 1 Supplementary text.docx for "Ganglioglioma deep transcriptomics reveals primitive neuroectoderm neural precursor-like population"

Supplementary results

Supplementary cluster assignment and differentially expressed gene analysis for ganglioglioma snRNA-seq

To try to mitigate undesirable technical artifacts, integration was performed for snRNA-seq data (by tumor-of-origin) using several methods (with widely varying parameters, including Seurat v4 CCA[21], Harmony[30], Scanorama[24], scVI[34], scANVI[71], scGen[35], BBKNN[50], and MNN[20]) including previously identified top performers[36]. Our top performers BBKNN and Scanorama managed to preserve segregation of the immune cells (microglial clusters 1, 5, 11, and 17 and lymphocyte clusters 7 and 28, s**Figure 1F-G** for BBKNN and not shown for Scanorama). BBKNN was also capable of segregating vascular cells (endothelial cluster 12 and vascular leptomeningeal cell (VLMC) cluster 22), oligodendrocyte cluster 20, and OPC cluster 21 (s**Figure 1F-G**). Beyond that, data integration attempts struggled to adequately balance preservation of meaningful biological variation over mitigation of undesirable batch effects (not shown except for BBKNN); for example, even BBKNN tended to only somewhat segregate neuron cluster 15 from astrocyte cluster 23 and the tumor neoplastic cell supercluster (s**Figure 1F-G**). In order to optimally preserve meaningful biological variation, we preferred the non-integrated data, and that is what we discuss throughout the main text and below.

Preliminary snRNA-seq cell assignments were confirmed by analysis of top differentially expressed genes (**sTable 2**). Cluster 1, 5, 11, and 17 top 5 DEG by Wilcoxon rank sum test encoded machinery for microglia-/macrophage-specific hemoglobin/haptoglobin uptake (*CD163*) and iron metabolism (*SLC11A1*) (both cluster 5) and MHC class II antigen processing (*CD74*) and presentation (*HLA-DRA* and *HLA-DPA1*) (both cluster 17)[51, 75]. Each cluster 1, 5, 11, and 17 top 5 included transcripts that represent microglia-specific markers in normal brain such as *USP53* and *SFMBT2* in cluster 1, *SLCO2B1* and *PLXDC2* in cluster 5, and *ADAM28* in cluster 11[51, 75]. Cluster 7 top 4 DEG encoded proteins responsible for T cell adhesion and function (*CD2*), T cell receptor signaling enhancement (*SKAP1*), and regulation of T cell development (*THEMIS*, *BCL11B*)[51, 75], consistent with this cluster being composed of T lymphocytes. Cluster 28 top 3 DEG encoded proteins responsible for B cell receptor function (*BANK1*), immunoglobulin (*IGKC*), and lymphoid development (*AFF3*)[51, 75], consistent with B lymphocytes populating cluster 28. Cluster 12 top 5 DEG included transmembrane transporters (*ABCB1* (involved in transport across the blood brain barrier) and *ATP10A*), a vascular injury hemostatic facilitator (*VWF*), and a modulator of development, cell proliferation, and differentiation (*MECOM*)[51, 75], consistent with this cluster representing endothelial cells. Cluster 22 top DEG included transcripts encoding VLMC markers (*DCN*, #1; *C7*, #7; *FLVCR2*, #18)[14, 76]. Cluster 15 top 5 DEG included transcripts encoding regulators of synaptic neurotransmitter release (*SYT1* and *SNAP25*), synaptic development and remodeling (*NRGN*), and numerous functions in myelinated axons (*CNTNAP2*)[51, 75]. Cluster 20 top 4 DEG were oligodendrocyte-specific markers in the context of the brain (*TMEM144*, *TF*, *EDIL3*, and *ST18*)[51, 75]. Cluster 21 top 5 DEG included 2 oligodendrocyte precursor cell-specific markers in the context of the brain (*PCDH15* and *DSCAM* relatively so) while the remainder were at least expected to be highly expressed in OPCs[51, 75]. Cluster 23 top 5 DEG included astrocyte-specific markers in the brain context (*CST3*, *MT3*, and *AQP4*) and were otherwise at least expected to be highly expressed in astrocytes[51, 75]. These results overall supported our prior provisional snRNA-seq cluster typing.

Moreover, preliminary snRNA-seq cell assignments were confirmed by inference of chromosomal copy number changes using inferCNV (**sFigure 1I**)[48]. Controls included oligodendroglioma samples (characterized by 1p,19q co-deletion; MGH36, MGH53, MGH93, and MGH97), which also contained other known alterations including amplifications and deletions[63]. Unlike oligodendroglioma, ganglioglioma does not have a defining copy number alteration, though a few recurrent copy number alterations have been identified in ganglioglioma, including del1p, del9, amp5, amp7, amp9, amp12, and amp19q[17, 49]. Using oligodendrocytes from the oligodendroglioma samples and ganglioglioma stromal neural cells as the normal copy number reference, this method confidently recalled the previously described copy number variations in the control oligodendrogliomas without any copy number variations identified in the normal reference cells. Among the ganglioglioma neoplastic-appearing cells, inferCNV identified clonal del1p and subclonal del14 exclusively in the putative neoplastic cell clusters in tumor 4, providing support that these clusters represented neoplastic cells.

Validation of provisional snRNA-seq cell typing using curated cell type gene signature sets

Cell types were systematically verified using the curated cell type signatures in MSigDB C8[60]. Validation was first approached on the level of cell typing by cluster. Transcript concentrations were calculated by cluster relative to all of the other clusters (i.e. average log2 fold-change), and these ganglioglioma cluster signatures were subject to gene set enrichment analysis (fgsea)[29]. GSEA was performed using all 32880 MSigDB C8 cell type gene signatures available as of Feb 2023 (i.e. unfiltered by organ, tissue, pathology, etc...). Top enriched gene sets were primarily microglial (clusters 1, 5, 11, and 17), T cell (cluster 7), B cell (cluster 28), or endothelial (cluster 12 and 22), confirming these stromal cell type assignments (**sTable 8**). For each of the neural stromal clusters (neuron cluster 15, oligodendrocyte cluster 20, OPC cluster 21, and astrocyte cluster 23), top enriched gene sets included the expected cell type, and signatures representing the expected cell type were generally enriched, consistent with expectations (**sTable 8**). Neoplastic clusters were typically associated with enrichment for signatures from more varied cell types: A mix of neuron and oligodendrocyte signatures (cluster 4); a mix of neuron and astrocyte signatures (clusters 3 and 8); a mix of neuron, oligodendrocyte, and OPC signatures (clusters 0, 9, 13, 18 and 26); a mix of astrocyte, neuron, and OPC signatures (cluster 6); or more varied (or less descript) signatures (clusters 2, 10, 16, 24, 25, 27, and 29; **sTable 8**). Among neoplastic clusters, cluster 14 was associated with enrichment of astrocyte signatures (**sTable 8**) despite relatively low expression of most astrocyte markers. This appears most consistent with a relatively astrocyte-like neoplastic cell cluster. Similarly, cluster 19 was associated with enrichment for neuron signatures (**sTable 8**) despite lack of significant expression of top neuron markers (including pan-neuron marker *RBFOX3*). This appears consistent with a relatively neuron-like neoplastic cell cluster. Hence, cluster-level probing of ganglioglioma cell types by gene set enrichment analysis appeared consistent with the provisional cell typing.

To further build upon ganglioglioma cell typing by the MSigDB C8 curated cell type signature sets, we subjected the individual ganglioglioma cell transcriptomic profiles to signature scoring by UCell[2, 60]. This approach confirmed the results above. Among neural C8 gene sets, top enriched gene sets included Lein neuron markers (cluster 15), Lein oligodendrocyte markers (cluster 20), Zhong PFC major types OPC and Fan embryonic OPC (cluster 21), and Fan embryonic CTX astrocyte 2 and GOCC astrocyte end foot (cluster 6 more so than cluster 23 for the former; **sTable 9**). Hence, interrogation of the ganglioglioma snRNA-seq data on a cluster- or cell-level using the large MSigDB collection of curated cell type gene signatures appeared to support our provisional cell typing.

Supplementary inference of snRNA-seq cellular hierarchy

CellRank/CytoTRACE results for snRNA-seq cluster 6 and/or neoplastic *CD34*+ cells appeared robust to varying preprocessing and data subsetting (including all data, exclusion of low complexity tumor 3 nuclei alone (the inclusion of which resulted in aberrantly high pseudotime for the tumor 3 nuclei), and subsetting by tumor of origin; not shown)[19, 31]. For RNA velocity analyses, ganglioglioma neoplastic cellular hierarchy was inferred using scVelo dynamical, stochastic, and steady state modes, which generally agreed with the high stemness of cluster 6 and/or *CD34*+ neoplastic cells (**Figure 2J-M**) though RNA velocity analysis was complicated by a significant proportion of uninformative phase portraits (not shown)[4].

Supplementary snRNA-seq ligand-receptor interactions

CellChat considers the geometric means of ligand and receptor expression, weighted by their agonists/antagonists, and has the advantages of working from a curated database and considering together complex subunits and cofactors (i.e. uses a macromolecular complex weighting network)[26].

Interestingly, it has been shown that PTPRZ1 plays a role in aggressive glioblastoma cell phenotypes[5]. Higher PTPRZ1 expression has been associated with poorer prognosis in both grade 2-3 glioma and glioblastoma[70]. Interestingly, we found evidence of substantial PTPRZ1 antagonism across gangliogliomas and a wide array of gliomas, including pilocytic astrocytoma and glioblastoma (**Figure 3A** and not shown)[15, 52]. Additional PTN pathway interactions with interesting patterns but unclear significance emerged including PTN - SDC2 (neoplastic specific but not sensitive), - SDC4 (cluster 6 specific), and - ITGAV+ITGB3 (neoplastic specific but not sensitive; not shown).

Results by CellChat were corroborated by parallel analysis with CellPhoneDB (not shown)[18]. Similar to CellChat, CellPhoneDB has the advantages of working from a curated interaction database and of accounting for multi-subunit complex formation. In contrast to CellChat, it infers likelihood of cell communication by co-overexpression of ligand-receptor pairs over generated null distributions.

Supplementary snRNA-seq neoplastic cell signaling

Among the most recurrently activated pathways among snRNA-seq neoplastic-appearing cells when comparing by cluster or overall were pro-inflammatory, complement and coagulation cascade, cytokine-cytokine receptor interaction, transcriptional misregulation in cancer, protein serine/threonine kinase, and various cancer site specific pathways (**Figure 4** and **sFigure 4A-B**). KEGG pathways of interest based on *a priori* interest and GSEA results were further visualized using Pathview (selected in **sFigure 4C-F**)[37]. Interpretation of such pathway-based diagrams for RNA was not expected to be straightforward given the complex negative and positive feedback loops in cells. However, of note, BRAF and PKB/AKT were both relatively overexpressed in neoplastic-appearing versus normal-appearing neural cells.

Supplementary snRNA-seq neoplastic transcriptional programs and putative drivers

Normal brain development is associated with a cascade of SOX family member activity[58]. Interestingly, among snRNA-seq neoplastic ganglioglioma cells, there appeared to be a cascade of SOX family member activities as well. Like *SOX2*, *SOX9* expression was associated with cluster 6 and/or *CD34*-expressing neoplastic cells as well as early pseudotime (and high potency, **sFigure 5G**). SOX9 (like SOX2) is expressed in neural stem and progenitor cells and is essential for stem cell maintenance and multipotentiality[58]. Depending upon context, SOX9 is further involved in facilitating gliogenesis, inhibiting neurogenesis, promoting oligodendrocyte differentiation, or terminal differentiation of astrocytes[58]. In normal adult brain, SOX9 is an astrocyte-specific marker[51, 75]. Interestingly, *SOX13* exhibited pseudokinetics similar to *SOX2* and *SOX9* (**sFigure 5H**). SOX13 is involved in differentiation of post-mitotic neurons, may be involved in oligodendroglial differentiation, and is normally found predominantly in adult OPCs and certain neurons more so than astrocytes and oligodendrocytes[51, 58, 75]. In contrast to *SOX2* and *SOX9* expression and regulon activity at early pseudotimes, *SOX4* and *SOX10* expression and activity appeared predominantly at intermediate pseudotimes (and potencies, **sFigure 5I-J**). Both these factors act later in lineage development normally as well[58]. SOX4 is a SOX C member important in transcriptional programming in early neuronal development as well as some glial/mesenchymal progenitors, particularly OPCs in the adult brain[51, 58, 75]. SOX10 is a SOX E member that promotes oligodendrocyte differentiation and is normally found in OPCs and oligodendrocytes[51, 58, 75].

Beyond the TCF7L2/MEIS1-PAX6 and SOX transcriptional cascades, top transcription factors for both cluster 6 and *CD34*+ neoplastic cells also included a number of other transcription factors that appeared specific for the primitive neoplastic cells. PR domain containing 16 (PRDM16), peroxisome proliferator activated receptor gamma co-activator 1 alpha (PPARGC1A), empty spiracles homeobox 2 (EMX2), and Mesenchyme Homeobox 2 (MEOX2) all had expression and regulon activity appear strongly *CD34*- and cluster 6-specific with briskly falling pseudokinetics (by CytoTRACE or SCENT, **sFigure 5K-N**). PRDM16 appears to control neural stem cell development and maintenance, appears essential for normal radial glial development[22]. PPARGC1A is a transcriptional co-activator for steroid receptors and nuclear receptors including thyroid hormone receptor, cAMP response element binding protein (CREB), and nuclear respiratory factors (NRFs). It is thereby a master regulator of mitochondrial biogenesis as well as normal neuron development and maintenance[13, 69]. This appears consistent with the increase in oxidative respiration pathways in the primitive neoplastic cells. EMX2 is normally astrocyte-specific in adult brains and is involved in counteracting SOX2 to control cell fate during normal development[7, 39]. MEOX2 is a good pan-neuron marker in the healthy adult brain[51, 75]. Its role in brain development is unclear, though it is increasingly appreciated to be capable of promoting glioma-associated aggressive features[56].

Other transcription factors involved in neural progenitor cell functions appeared to be most prevalent and active in the most potent neoplastic cells but only gradually decreased or plateaued with differentiation (**sFigure 5O-Q**). For example, Ras responsive element binding protein 1 (RREB1) potentiates the transcriptional activity of NEUROD1, which counteracts SOX2 activity, and RREB1 may be involved in Ras/Raf-mediated cell differentiation[40]. In the healthy adult brain, it normally is found more so in microglia than in astrocytes and to a limited extent in some neurons[51, 75]. Also included in this group are the transcription factors glioma-associated oncogene family zinc finger 2 (GLI2, SHH signaling during development[41]), CP2-Like 1 (TFCP2L1, facilitates pluripotency in embryonic stem cells[61]), B-Cell Lymphoma 6 (BCL6, promotes neuronal differentiation[6]), Kruppel-like zinc finger 12 (KLF12, AP2 repressor in neural progenitors[45]), regulatory factors X 4 and 7 (RFX4 and RFX7; RFX genes are master regulators of neurogenesis[27]).

Supplementary snRNA-seq immune landscape

SnRNA-seq myeloid cells

Top snRNA-seq microglial transcriptional programs were nominated in an unbiased manner using SCENIC (**sFigure 6E-F**). Most of the top nominated transcription factors among microglia were in common to all microglia clusters, which included RUNX2, RREB1, FOXO3, FOXP1, FOXP2, MEF2A, MEF2C, BBX, IKZF1, ETS2, FLI1, REL, ELF1, MAF, SPI1, and JDP2. RUNX2 is constitutively expressed in microglia[44]. FOXO3 is an activator of microglia-induced inflammation[57]. FOXP1 and FOXP2 each appear to play important roles in brain development and human neuropathology, but their functions have largely been studied in neural cells rather than microglia[1, 46]. Clusters 1 and 5 did not have any standout cluster-specific transcription factors. In contrast, cluster 11 specific transcription factors included TAL1 and RUNX1 which stood out as the top two nominated transcription factors and are associated with primitive microglial programming including during very early embryonic development[67]. This is consistent with the cellular hierarchy determination discussed earlier in which cluster 11 cells occupied the earliest pseudotimes. Cluster 17-specific factors CEBPA and PGAM2 were among the top 5 nominated transcription factors for that cluster. CEBPA in the context of SPI1 has been found to be sufficient for differentiation of induced pluripotent stem cells to microglia[11]. PGAM2 has been found in experimental models to be crucial to microglia-mediated necroptosis and inflammation[65].

SnRNA-seq lymphocytes

Review of top snRNA-seq differentially expressed genes concurred with single marker-based typing (**sTable 2**). Cluster 7 top DEG included the classic T cell markers *CD2*, *CD3G*, and *CD247* (*CD3Z/H/Q*), and top DEG otherwise are expected to be relatively highly expressed on T lymphocytes[51, 75]. Top DEG for cluster 28 included *CD20* (*MS4A1*), B lymphocyte kinase (*BLK*), B cell specific glycoprotein B29 (*CD79B*), B cell specific transcription factor (*PAX5*), and Fc receptors (*FCRL1*, *FCRL2*, *FCRL5*) entirely consistent with cluster 28 representing B cells.

Lymphocyte significant regulons and transcription factors were nominated in an unbiased fashion using SCENIC (**Figure 7J-K**). Cluster 7 top SCENIC nominated regulons included the important CTL exhaustion-associated transcription factor EOMES as well as the pre-exhaustion transcription factor TBX21, again consistent with a spectrum of CTL states between precursor and exhaustion. Other top transcription factors included RUNX3 which plays a crucial role in CTL development; ZNF831 which is specific to T cells and NK cells but is otherwise not well understood; STAT4 - a key regulator of helper T cell differentiation; GATA3 - an important T cell developmental regulator; ETS1 which controls lymphocyte differentiation, survival, and proliferation; and other factors with roles in T lymphocyte development-RREB1, BATF, IKZF1, and MYB. Cluster 28 most significant regulons by SCENIC were SPIB, which may be required for B cell receptor signaling and B cell development; PAX5, which commits to and maintains in B cell lineage; and POU2AF1- a factor essential for B cell antigen response.

Supplementary validation with CITE-seq

Overall CITE-seq cellular composition

To validate results from snRNA-seq, we performed CITE-seq on freshly-dissociated cells from 3 gangliogliomas. CITE-seq data passed quality control for 3725 cells from 3 tumors (2 also sequenced with snRNA-seq and 1 unique, see **Supplementary Table 1** for sample information). Non-linear dimensional reduction and k-nearest neighbor search[8, 21, 55, 59] yielded 14 clusters (**sFigure 8A**). Cell type was identified using a multi-pronged approach. Parallel antibody probing for CD14 (myeloid marker), CD8A (cytotoxic T lymphocyte marker), CD4 (helper T cell marker, also present on some APCs), and CD34 (endothelial marker but also of interest to us due to expression in the neuroectoderm) identified a myeloid supercluster (clusters 04, 05, and 06), a lymphoid supercluster (clusters 02, 07, 10, and 12), and an endothelial cluster (cluster 11) (**sFigure 8B**). RNA expression of other general cell type markers (**sFigure 8C**) further supported/refined the presence of a myeloid supercluster (*PTPRC*- and *CD14*-expressing, clusters 04, 05, and 06), a lymphocyte supercluster (*PTPRC*-expressing but not *CD14*-expressing, clusters 02, 07, 10, and 12), and an endothelial cluster (*VWF*-expressing, cluster 11), leaving the macroglial- and neural-like cells (cluster 00, 01, 04, 08, 09, and 13) [51, 75]. Among the neural(-like) cells, cluster 13 cells tended to express astrocyte markers (*AQP4, SLC1A2, GFAP*) but not other neural lineage specific markers (e.g. neuronal *RBFOX3*, oligodendrocyte *MOBP/CNP/MOG/MBP*, or OPC *GPR17/CSPG4/OLIG1/OLIG2*)[51, 75]. The overall pattern of marker expression appeared most consistent with that of normal astrocytes. Cluster 09 cells, in contrast tended to express the pan-neuron-specific marker *RBFOX3*, with moderate expression of macroglial lineage markers including *GFAP*, *CSPG4*, *OLIG1*, and *OLIG2* and an actively proliferating component, consistent with this cluster representing neoplastic neuronal-like cells[51, 75]. In all, there were 15 neoplastic *CD34*+ cells, 10 from tumor 1 and 5 from tumor 3. Cluster 09 was particularly enriched for *CD34*, with 10/130 *CD34*+. Cluster 08 cells tended to express a number of oligodendrocyte/OPC and OPC markers including the OPC-specific marker *GPR17* but also tended to express *GFAP*, consistent with this cluster representing abnormal OPC-like tumor cells[51, 75]. The other neural clusters (00, 01, and 03) exhibited atypical patterns of glial lineage markers but no *RBFOX3*, consistent with these representing various macroglia-like tumor cells with nonspecific marker expression. Analysis of top DEG and cell typing by label transfer with reference atlases concurred with these individual marker-based assessments (**Supplementary Table 6**, not shown).

Copy number variation was inferred using the inferCNV package (not shown)[48]. When querying ganglioglioma cluster 00, 01, 03, 08, and 09 cells, there appeared to be alterations in cells from tumor 3. In particular, neoplastic cells appeared to have del1p. Subclones from clusters 00 and 08 had del1 and subclones of this had del1;del9. Del1q and del9 have been previously identified in ganglioglioma[17, 49]. This reinforces the identification of cells from clusters 00, 01, 03, and 08 as neoplastic macroglia-like cells.

CITE-seq data integration (batch=tumor) was performed analogously to that performed for snRNA-seq. Similar to what was found for snRNA-seq, the benefits of CITE-seq data integration were consistently outweighed by the loss of meaningful biological variation. Consequently, we continued to use unmanipulated CITE-seq data for downstream analysis.

CITE-seq integration with snRNA-seq data (batch=technique) was interrogated using several methods (Seurat v4 CCA[21], Harmony[30], Scanorama[24], scVI[34], scANVI[71], scGen[35], BBKNN[50], and MNN[20]). Seurat CCA was unique in preservation of meaningful biological variation (**sFigure 8D-F**). SnRNA/CITE seq microglial clusters co-clustered, snRNA/CITE-seq lymphocyte clusters co-clustered, and snRNA/CITE-seq endothelial clusters co-clustered. As expected, neoplastic cells did not necessarily co-cluster as neatly as their stromal counterparts. Notable is the lack of significant CITE-seq cells co-clustering with snRNA-seq neuroectoderm neural precursor cell-like cluster 6. Such an integration product may serve as a reasonable ganglioglioma reference atlas for subsequent studies. However, except as noted explicitly below, for the sake of downstream analysis, we preferred to continue with our test-validation approach (test=snRNA-seq, validation=CITE-seq), so downstream analysis was continued with unintegrated CITE-seq data.

CITE-seq cell types were systematically verified using the curated cell type signatures in MSigDB C8[60]. Validation was first approached on the level of cell typing by cluster as done previously with the snRNA-seq data. Top enriched gene sets were primarily microglial, macrophage, or myeloid (clusters 04, 05, and 06); T cell , NKT cell, or lymphocyte (clusters 02, 07, 10, and 12), or endothelial (cluster 11), confirming these stromal cell type assignments (**sTable 10**). For astrocyte cluster 13, top enriched gene sets included the expected cell type, and signatures representing the expected cell type were generally enriched, consistent with expectations (**sTable 10**). Neoplastic clusters were typically associated with enrichment for signatures from more varied cell types: A mix of neuron and astrocyte signatures (cluster 03); a mix of neuron, oligodendrocyte, OPC, and astrocyte signatures (clusters 01 and 08); or a mix of neuron, OPC, and astrocyte signatures (clusters 00 and 09; **sTable 10**). Hence, cluster-level probing of ganglioglioma CITE-seq cell types by gene set enrichment analysis appeared consistent with the provisional cell typing.

To further build upon ganglioglioma CITE-seq cell typing using the MSigDB C8 curated cell type signature sets, we subjected the individual ganglioglioma cell transcriptomic profiles to signature scoring by UCell[2, 60]. This approach confirmed the results above. Among brain/neural C8 gene sets, top enriched gene sets included Zhong PFC major types OPC (cluster 08) and GOCC astrocyte end foot (clusters 08 and 13; **sTable 11**). Hence, interrogation of the ganglioglioma CITE-seq data on a cluster- or cell-level using the large MSigDB collection of curated cell type gene signatures appeared to support our provisional cell typing.

CITE-seq neoplastic cellular hierarchy

CellRank/CytoTRACE of CITE-seq neoplastic cells revealed cluster 08, OPC-like cells to be relatively primitive (**sFigure 8G-H**)[31]. Comparison of neoplastic *CD34*+ cells to *CD34*- cells revealed the former to generally be of earlier pseudotime (**sFigure 8I**). When combining the snRNA-seq and CITE-seq data (using integration results from Seurat CCA as above), the most primitive snRNA-seq cells (i.e. cluster 6 and/or neoplastic *CD34*+ cells) were at earlier pseudotimes than CITE-seq cluster 08 and did not appear to have a correlate in the CITE-seq dataset (**sFigure 8J**). This may be due to undersampling in the CITE-seq data given the order of magnitude difference in the dataset sizes. This underscores the value of sequencing more cells/nuclei to detect rare tumor cells of interest, which we expected tumor stem cells to be. These findings are consistent with our *a priori* hypothesis and our snRNA-seq findings regarding a *CD34*+ neuroectoderm neural precursor-like stem cell population.

CITE-seq neoplastic cell ligand-receptor communication

Ganglioglioma CITE-seq cell significant ligand-receptor interactions were interrogated by CellChat[26]. Similar to what was found in the snRNA-seq data, this analysis suggested neoplastic tumor cell PTPRZ1 targeting by PTN, the latter largely produced by the neoplastic cells (**sFigure 8K**). This interaction appeared particularly strong in its targeting of PTPRZ1 in the relatively primitive, OPC-like cluster 08. These results by CellChat were corroborated by parallel analysis with CellPhoneDB (not shown)[18].

CITE-seq neoplastic cell signaling

CITE-seq cells were subject to GSEA for GO, KEGG, Hallmark, Oncogenic, Immune, Reactome, and Wikipathways[60, 68]. When comparing Hallmark pathway enrichment of neoplastic-appearing ganglioglioma CITE-seq cells (clusters 00, 01, 03, 08 and 09) to normal-appearing neural cells (cluster 13), a number of cancer-associated pathways appeared to have significant enrichment, including epithelial mesenchymal transition, hypoxia, interferon gamma response, p53 signaling, TNFα via NF-κB signaling, apoptosis, mTORC1 signaling, and unfolded protein response (**sFigure 8L**). Also of note, the neoplastic cells exhibited significantly decreased oxidative phosphorylation, consistent with the Warburg effect typical of neoplastic cells and offering further support for the provisional identification of cells as neoplastic or normal[66]. Among KEGG pathways, the antigen processing and presentation pathway was particularly enriched followed by p53 signaling, cell adhesion, spliceosome, cell cycle, and pathways in cancer (**sFigure 8M**). Oxidative phosphorylation was again significantly depleted. Similar themes emerged when analyzing Oncogenic, Immune, and Reactome pathways (not shown).

When comparing neoplastic *CD34*+ cells to *CD34*- cells, some similarities and differences emerged when comparing with the bulk neoplastic cell enriched pathways (**sFigure 8L-M**). For example, among the Hallmark pathways, particularly strong similarities included epithelial mesenchymal transition, p53 pathway, and apoptosis. Interestingly oxidative phosphorylation pathway depletion was not found in comparing *CD34*+ to *CD34*- cells, and there was in fact a trend in the opposite direction. Also interesting is that for the mTORC1 signaling, significant depletion was observed in *CD34*+ compared to *CD34*- cells, opposite what was seen when comparing neoplastic to normal appearing neural cells.

KEGG pathways of interest based on *a priori* interest and GSEA results were further visualized using Pathview[37]. Of interest was the intact expression of *BRAF* itself and central downstream mediators (*JUN* and *FOS*) and expected transcriptional programs (e.g. *VEGF*, *MMP*, and *IL8*) among the neoplastic cells compared to normal-appearing neural cells. Comparison of *CD34*+ to *CD34*- neoplastic cells hinted at some similar dysregulation in MAPK signaling.

These results were further confirmed by analysis of differential gene expression among the neural(-like) clusters. For example, by Wilcoxon rank sum test, comparison of neoplastic cells with normal appearing neural cells revealed enrichment (in the former) of MAPK downstream transcription factor-encoding transcripts *JUNB*, *FOS*, and *FOSB* and downstream induction of expression of *VEGFA*. Other transcripts notably increased in neoplastic versus stromal cells included stem/glioma cell marker *POSTN*; heat shock/unfolded protein stress response *HSPA1A*; cell motility/metastasis *MALAT1* and *CD9*; p53 signaling *NDRG1*, *EGR1*, and *MDM2*; *APOD* which is overexpressed in low grade gliomas (as well as the aging brain, Alzheimer’s disease, schizophrenia, and stroke), inhibits tumor and vascular cell proliferation, has expression driven by MAPK signaling (e.g. via JNK/MAPK8, ERK1/MAPK3, ERK2/MAPK1), and expression lost in the course of progression to (secondary) high grade glioma[16, 25, 38, 53]; *NEAT1* which may play numerous roles in glioma tumorigenesis, proliferation, glycolysis, and immune evasion (via PD-L1) with high expression associated with poorer outcomes[10, 23, 72, 74]; and *CHI3L1*, expressed on healthy astrocytes and overexpressed in numerous tumors, which may play pleiotropic roles in glioma, including PI3K/AKT-mediated cancer cell growth induction, MAPK/ERK1/2/VEGF(R)-mediated angiogenesis, and macrophage (pro-tumor M2) immune suppression[9, 73].

CITE-seq neoplastic gene regulatory networks and putative drivers

Active transcription factors and their transcriptional programs were nominated in an unbiased manner using SCENIC[54]. Among neoplastic CITE-seq cells, similar to what was found by snRNA-seq, SOX2 appeared to have significantly increased activity (and transcript expression) among certain tumor cells, particularly those with greater stemness and/or *CD34*-expression (**sFigure 8N-O**). LMO2 similarly had increased regulon activity and RNA expression among more potent and/or *CD34+* ganglioglioma neoplastic cells (**sFigure 8N-O**). In the healthy brain, LMO2 is normally mostly expressed in microglia; however, it has been found to be overexpressed in gliomas, especially in glioma stem cells, where it may promote tumorigenesis, invasion, and angiogenesis[28, 47, 51, 75]. PRRX1 also seemed to have activity and concentration associated with cell stemness in the ganglioglioma neoplastic cells (**sFigure 8N-O**). This factor tends to be expressed in healthy OPCs but is expressed more highly in glioma, where conflicting data exists as to its role in cancer-associated phenotypes[12, 62]. Another SOX B1 member, SOX3, regulon activity and RNA concentration appeared correlated with stemness and *CD34*-status but only among the neuron-like subset of tumor cells (**sFigure 8N-O**). Interestingly, SOX3 is expressed in late ectodermal cells, and it maintains neural progenitor cells and inhibits astrocyte lineage differentiation[58].

CITE-seq immune landscape

CITE-seq myeloid cells

CITE-seq myeloid cells were initially typed by inspection of cell markers. Clusters 04, 05, and 06 were *PTPRC*-, *ITGAM*-, and *CD14*-expressing, consistent with representing microglia or macrophages (**sFigure 8P**)[51, 75]. Cluster 06 expressed *P2RY12*, *TYROBP*, and *SPI1* but not appreciable *CD44* or *SIGLEC1*, consistent with microglial identity (**sFigure 8P**)[51, 75]. Cluster 04 expressed *TYROBP* and *SPI1* but not *CD44* or *SIGLEC1*, also most likely microglia (**sFigure 8P**)l[51, 75]. Cluster 05 cells generally also expressed *TYROBP* and *SPI1*, likely primarily microglia (**sFigure 8P**)[51, 75].

CITE-seq myeloid cells were also typed in Seurat by label transfer[21]. Using annotated tumor reference atlases differentiating microglia from macrophages[52, 64] for label transfer based on nearest neighbors mapping, the bulk of myeloid cells were consistently more strongly identified with an overall microglial rather than macrophage signature (not shown). On balance, this appears to suggest that the vast majority of ganglioglioma-associated myeloid cells are microglia, consistent with the snRNA-seq findings.

For myeloid cells, a distinction of interest was that between M1 (potentially anti-tumor) and M2 (potentially pro-tumor) polarization[42]. When looking at individual cell markers, we noted clusters 04, 05, and 06 all appeared largely *CD14+CD68+CD163*+, consistent with immunosuppressive/tissue reparative/tumor promoting M2 polarization (**sFigure 8P**). However, when ganglioglioma cells were subject to signature scoring (by UCell) based on established M1 and M2 signatures, the two scores appeared positively correlated both within each cluster and among cells of the supercluster (not shown)[2, 43], consistent with snRNA-seq results showing aberrant co-activation of M1 and M2 transcriptional programs.

We next evaluated classic pro-inflammatory and anti-inflammatory markers on the myeloid cells. Myeloid cells lacked appreciable expression of the classical pro-inflammatory markers *IL1A*, *IL6*, and *CD40* (**sFigure 8P**)[3]. Cluster 06 did contain some cells with *TNF*, another pro-inflammatory cytokine, expression (**sFigure 8P**)[3]. In terms of anti-inflammatory activity, cluster 05 and 06 cells tended to express significant levels *TGFB1*, but myeloid cells did not express appreciable levels of *IL10* (**sFigure 8P**)[3]. In fact, *TGFB1* and *TNF* transcript concentrations appeared to be positively correlated in ganglioglioma myeloid cells (not shown). Moreover, when looking at the broader cellular context, lymphocytes from cluster 02, 07, and 12 all expressed significant *TGFB1* and tumor clusters 08 and 09 expressed substantial *CSF1*, which promote immunosuppressive and pro-inflammatory properties in macrophages, respectively (not shown)[3]. Interestingly, myeloid cell concentrations of lymphocyte co-stimulatory and co-inhibitory ligands appeared positively correlated throughout the supercluster (not shown). These results further support there being ganglioglioma-associated immune cell spectra of aberrant activation and activities.

Myeloid cell intercellular communication was interrogated in an unbiased manner by CellChat and CellPhoneDB[18, 26]. Top-ranked pathways included MHC class II presentation to all myeloid clusters, primarily from other myeloid cells or lymphocytes (not shown). This appears to represent potential for immune cell activation and anti-tumor cell activity. The presence of active pro-inflammatory myeloid cell activity was supported by nomination of CSF1-CSF1R interaction targeting cluster 05 and 06 myeloid cells, cluster 05 and 06 cell CXCL16-CXCR6 interaction with the lymphocyte clusters, and cluster 06 TNF − TNFRSF1B interaction for all immune clusters except 04 and 10 (not shown). Also of note are pro-inflammatory factors among the myeloid cell cluster top differentially expressed genes, such as *CSF1R*, *AIF1*, and *TYROBP1* (**sTable 6**). On the other hand, anti-inflammatory TGFB1-TGFBR1/2 was nominated as a significant interaction (not shown). Also nominated was cluster 05 and 06 interaction with both CD28 and CTLA4 in cluster 12 (T regs, not shown). These results support the presence of both pro-inflammatory and anti-inflammatory intercellular signaling.

We were also interested in interrogating for myeloid cell C1Q and SPP1 status because C1Q expression appears to be associated with T lymphocyte recruitment and activation whereas SPP1 expression appears to be associated with tumor growth and metastasis[32, 33]. Among the top ligand-receptor interactions by CellChat were SPP1 interactions, primarily from neoplastic appearing cells, with receptors on all of the immune clusters except 04 and 10 (not shown). On the other hand, *SPP1* expression was overall low among ganglioglioma myeloid cells whereas C1Q component transcripts were highly expressed in clusters 04 and 06 (not shown). Moreover, complement activation was also nominated as significant interaction from clusters 04, 05, and 06 targeting clusters 05 and 06 by CellChat (not shown). These observations overall suggest predominantly C1Q+SPP1- myeloid cell state, which would be expected to facilitate anti-tumor lymphocyte activities.

CITE-seq lymphocytes

CITE-seq lymphocytes were initially typed by inspection of individual cell markers. Ganglioglioma clusters 02, 07, 10, and 12 were largely *PTPRC*-, *CD2*-, and *CD3*-expressing (**sFigure 8Q-R**), consistent with these clusters collectively representing a lymphocyte supercluster[3]. The bulk of cluster 02 cells appeared *CD2*+*CD3*+*CD4*-*CD8*+, *GZMA*+, and *PRF1*+, consistent with these cells representing cytotoxic T lymphocytes(**sFigure 8Q-R**)[3]. On the other hand, a subpopulation of cluster 02 did not express *CD3* or *CD8* but did express the natural killer cell-associated *KIR2DL4* and *NCAM1* transcripts in addition to cytolytics (**sFigure 8Q-R**)[51, 75]. This appears most consistent with a minority population of NK cells within cluster 02. Cluster 12 contained cells strongly expressing *CD2*, CD3-encoding RNAs, *CD4*, *IL2RA* (alias *CD25*) and *FOXP3*, consistent with this cluster representing regulatory T cells (**sFigure 8Q-R**)[51, 75]. Cluster 07 cells tended to express *CD2*, CD3-encoding RNAs, and *CD4* but not *CD8*- or cytolytic-encoding RNAs, consistent with these cells representing helper T cells (**sFigure 8Q-R**)[51, 75]. Cluster 10 cells were less descript in terms of their expression profiles, with feature counts an order of magnitude lower than that for the other lymphoid cells precluding fine characterization (not shown). Hence, preliminary assessment of ganglioglioma-infiltrating lymphocytes by individual markers from CITE-seq identified a cytotoxic T lymphocyte cluster (with a small minority of NK cells), a regulatory T cell cluster, and a helper T cell cluster.

To fine tune our understanding of ganglioglioma lymphocyte states, we interrogated CITE-seq individual cell markers further. MHC class I antigen presentation to CTL T cell receptors is fundamental to CTL activity including activation. MHC class I component transcript expression was robust for much of the ganglioglioma neoplastic cells (not shown), suggesting the foundation for anti-tumor CTL activity was present. Additionally, CTL activation depends upon CTL co-stimulatory ligand-receptor interactions and is antagonized by CTL co-inhibitory ligand-receptor interactions. Ganglioglioma neoplastic cells had variable expression of CTL co-stimulatory and co-inhibitory ligands (not shown), making it not entirely clear where the balance was in terms of a milieu promoting CTL activation versus CTL inactivation/anergy. Cluster 02 CTLs tended to express higher levels of co-inhibitory receptor-encoding transcripts *PDCD1*, *LAG3*, and *TIGIT* than co-stimulatory receptor RNAs and higher expression of exhaustion-related transcription factor RNAs *EOMES* and *TOX* than the precursor or pre-exhaustion factors *TBX21* and *HNF1A* (not shown)*,* which all together may be consistent with some degree of CTL dysfunction and exhaustion[3]. In addition, cluster 12 T regs had significant upregulation of *TNFRSF18*, which is upregulated upon T reg activation and thought to play a key role in immunological self-tolerance. From these results, it appeared that there was significant potential for pro-inflammatory (and anti-tumor) TIL activity but that there was also significant immunosuppression, including activated tumor-infiltrating regulatory T cells.

CITE-seq lymphocyte typing by UCell multi-gene signature scoring[2] and Seurat-based label transfer[21], largely agreed with the lymphocyte typing by individual markers discussed above. Lymphocytes were scored (rank-based, by UCell) based on subtype signatures previously evaluated in a single nucleus/cell context[3]. As expected, cluster 12 scored highly with the regulatory T cell and anti-inflammatory signatures (not shown)[3]. Also consistent with preliminary assignment, cluster 02 UCell scores were significantly higher for the CD8 T cell activation, pro-inflammatory, and cytolytics effector pathway signatures (not shown)[3]. In addition, particularly because of the ganglioglioma (snRNA-seq and CITE-seq) myeloid cell aberrant M1-M2 program co-activation (as opposed to polarization) observed, we were interested in assessing for T lymphocyte M1 polarizing versus M2 polarizing signatures[3]. Interestingly, CITE-seq lymphocyte M1-polarizing and M2-polarizing signatures were positively correlated (not shown)[3]. Seurat-based label transfer concurred with these results (not shown)[3, 21]. Hence, multi-gene signature scoring and label transfer concurred with CITE-seq lymphocyte typing by individual markers. Furthermore, signature scoring and label transfer revealed aberrantly (positively) correlated M1-polarizing and M2-polarizing signatures among ganglioglioma TILs. This appears consistent with the results seen among the ganglioglioma myeloid cells (with M1 and M2 program co-activation rather than polarization). The aberrant lymphocyte programs may even account for the myeloid cell aberrant activation.

CITE-seq lymphocyte intercellular signaling was interrogated in an unbiased manner by CellChat and CellPhoneDB[18, 26]. Cluster 02 was a particularly strong signal recipient (and relatively minor origin of communication, not shown). This included robust MHC class I and CTL T cell receptor communication from every ganglioglioma cluster except from the apparently normal astrocytes, making this ligand-receptor interaction the most significant in the CITE-seq cells overall (not shown). The second most significant CITE-seq intercellular interaction pathway was MHC class II interaction with receptors on cluster 07 helper T cells, cluster 12 regulatory T cells, and all of the myeloid clusters (not shown), consistent with their prior assignments. Hence, ligand-receptor analysis suggested robustly active antigen processing and presentation in ganglioglioma.

Supplementary Discussion

In this study, we weighed in on numerous controversies in the analysis of snRNA-seq, CITE-seq, and stRNA-seq. For instance, Seurat v4 CCA was uniquely able to largely preserve biological variation while limiting undesirable technical artifacts in performing the difficult task of integration of snRNA-seq data with CITE-seq data (but poor performing with integration by tumor-of-origin within those datasets). This experience may assist others in the task of choosing among the >50 available algorithms for this common data integration task.

Along these lines, we also compared several methods for cellular hierarchy determination. Though RNA velocity-based inference of cellular hierarchy has become popular, this method remains significantly limited by the common occurrence of uninformative phase portraits. We, too, encountered this limitation, particularly with the CITE-seq data. In comparing several RNA velocity-based methods, signaling entropy rate, and CellRank/CytoTRACE, our data supports use of CellRank/CytoTRACE as the gold standard to beat. That said, completing multiple of these methods is advantageous to highlight meaningful (corroborated) results and deemphasize artifacts. For example, we found nuclei/cells of low usefulness due to low complexity (artifact) could be easily filtered out by their aberrantly high CellRank/CytoTRACE pseudotime combined with middling signaling entropy rate.

In addition, we compared several methods for stRNA-seq cell typing. In general these methods fell along the lines of kNN-based label transfer, deconvolution/decomposition, and rank-based signature scoring. In the end, most algorithms performed poorly. However, cell2location proved robust and was supported by UCell-based signature scoring.

Supplementary cited literature

31.	Lange M, Bergen V, Klein M, Setty M, Reuter B, Bakhti M, Lickert H, Ansari M, Schniering J, Schiller HB, Pe’er D, Theis FJ (2022) CellRank for directed single-cell fate mapping

40.	Morgani SM, Su J, Nichols J, Massagué J, Hadjantonakis AK (2021) The transcription factor Rreb1 regulates epithelial architecture, invasiveness and vasculogenesis in early mouse embryos. Elife 10

75.	Human Protein Atlas proteinatlas.org

76.	HuBMAP Azimuth https://azimuth.hubmapconsortium.org/references/

Supplementary methods

Data integration

To try to mitigate undesirable technical artifacts, data integration was performed for snRNA-seq (by library), for CITE-seq (by library), and of snRNA/CITE-seq by method. In each case, several methods were attempted with widely varying parameters, including Seurat v4 CCA[21], Harmony[30], Scanorama[24], scVI[34], scANVI[71], scGen[35], BBKNN[50], and MNN[20]. Scib wrapper was used where possible[36]. Attempts were made to optimize based on NMI, ARI, ASW (cell type), and isolated label scores as described previously[36].

MSigDB cell type validation

MSigDB C8 cell type gene signatures were accessed Feb 2023[60]. Ganglioglioma snRNA-seq or CITE-seq average log2-fold-change was calculated for each cluster compared to the remaining clusters within the data set (within snRNA-seq or CITE-seq only) within Seurat v4. The entire MSigDB C8 collection was used to query these cluster-based profiles. Fgsea v1.22.0 was used for gene set enrichment using defaults[29]. For UCell v2.0.1 gene signature scoring of individual snRNA-seq or CITE-seq cells, the C8 collection was filtered for signatures containing neural terms including ‘BRAIN’, ‘NEUR’, ‘ASTRO’, ‘OLIGOD’, or ‘OPC’.
